# Phosphatidic acid produced by phospholipase D is required for hyphal cell-cell fusion and fungal-plant symbiosis

**DOI:** 10.1101/849232

**Authors:** Berit Hassing, Carla J. Eaton, David Winter, Kimberly A. Green, Ulrike Brandt, Matthew S. Savoian, Carl H. Mesarich, Andre Fleissner, Barry Scott

## Abstract

Although lipid signaling has been shown to serve crucial roles in mammals and plants, little is known about this process in filamentous fungi. Here we analyse the contribution of phospholipase D (PLD) and its product phosphatidic acid (PA) in hyphal morphogenesis and growth of *Epichloë festucae* and *Neurospora crassa*, and in the establishment of a symbiotic interaction between *E. festucae* and *Lolium perenne*. Growth of *E. festucae* and *N. crassa* PLD deletion strains in axenic culture, and for *E. festucae* in association with *L. perenne*, were analysed by light-, confocal- and electron microscopy. Changes in PA distribution were analysed in *E. festucae* using a PA biosensor and the impact of these changes on endocytic recycling and superoxide production investigated. We found that *E. festucae* PldB and the *N. crassa* ortholog, PLA-7, are required for polarized growth, cell fusion and ascospore development, whereas PldA/PLA-8 are dispensable for these functions. Exogenous addition of PA rescues the cell-fusion phenotype in *E. festucae*. PldB is also crucial for *E. festucae* to establish a symbiotic association with *L. perenne*. This study identifies a new component of the cell-cell communication and cell fusion signaling network that controls hyphal morphogenesis and growth in filamentous fungi.

## Introduction

*Epichloë festucae* is a filamentous ascomycete fungus that forms a largely beneficial interaction with the agriculturally important grass *Lolium perenne* (Schardl, 2010, Tanaka *et al*., 2012). This endophyte colonises the apoplast of aerial tissues of the infected plant to form a restricted hyphal network *in planta* (Christensen *et al*., 2008, Scott *et al*., 2012). *E. festucae* also forms an epiphyllous hyphal network on the leaf surface that is connected to the endophytic network following exit of these hyphae through the cuticle by formation of an appressorium-like structure called an expressorium (Becker *et al*., 2016).

Growth of the fungus *in planta* is highly regulated and many well-known signaling molecules and pathways are required for a functional interaction. These include signaling via reactive oxygen species (ROS), the cell wall integrity (CWI) mitogen-activated protein kinase pathway (MAPK) and the stress-activated MAPK pathway (Becker *et al*., 2015, Eaton *et al*., 2010, Tanaka *et al*., 2006). Many signaling pathways require interplay between membrane-bound and cytosolic proteins. In mammals and plants, this frequently involves lipid signaling via different lipid species which target various proteins to the membrane. While studies on lipid signaling have received considerable attention in mammalian cells, plants and yeasts, few studies have focused on the role of this process in filamentous fungi. Phosphatidic acid (PA), produced by phospholipase D (PLD)-catalysed hydrolysis of phosphatidyl choline (PC), is an important second messenger in plant and mammalian cells, where it is required for the correct temporal and spatial localization and activity of many proteins (Selvy *et al*., 2011, Jenkins & Frohman, 2005). Through these interactions PA influences cell growth, cytoskeletal rearrangement, vesicle formation and trafficking as well as superoxide production (Selvy *et al*., 2011, Wang, 2005). In plants, PA is additionally involved in pathogen and stress responses, stomatal opening and in signaling by the hormone, abscisic acid (Zhang *et al*., 2009, Yao & Xue, 2018). PA also has an important role in both endo- and exocytosis vesicle trafficking. Specifically, PA is involved in vesicle transport from the endoplasmic reticulum (ER) to the Golgi apparatus, including vesicle budding from the Golgi apparatus via ADP ribosylation factor Arf1 signaling and epidermal-growth factor (EFR)-mediated endocytosis (Chen *et al*., 1997, Ktistakis *et al*., 1996, Shen *et al*., 2001). Furthermore, in chromaffin cells PA accumulates at sites of exocytosis where it is required for the formation of fusion-competent granules (Tanguy *et al*., 2019, Zeniou-Meyer *et al*., 2007). In mammalian and plant cells, PA also contributes to superoxide production by activating the multi-protein NADPH oxidase (Nox) complex, via both the direct activation of the catalytic subunit as well as through the recruitment of the regulatory subunits. Depletion of PLDα in *Arabidopsis* leaves and the inhibition of PLD in tobacco pollen tubes reduces NADPH oxidase activity (Potocký *et al*., 2012, Sang *et al*., 2001). In addition, the direct interaction between RBOHD and PA is required to achieve stomatal closure upon abscisic acid signaling (de Jong *et al*., 2004, Zhang *et al*., 2009). In animal cells PA has been found to interact with multiple components of the Nox complex, such as the catalytic subunit gp91^phox^ (or Nox2) (Kanai *et al*., 2001, Karathanassis *et al*., 2002, Ago *et al*., 2003, Qualliotine-Mann *et al*., 1993, Taylor *et al*., 2004, Taylor *et al*., 2012). In a cell-free system, PA was shown to be crucial for the activation of the mammalian NADPH oxidase (Qualliotine-Mann *et al*., 1993, Palicz *et al*., 2001).

The role of PA in yeast and filamentous fungi has been analysed by studying the phenotype of PLD deletion strains. In *Saccharomyces cerevisiae*, deletion of *SPO14,* the homolog of mammalian PLD1, results in a loss of fusion of Golgi-derived vesicles to the prospore membrane during sporulation (Rudge *et al*., 1998, Riedel *et al*., 2005, Nakanishi *et al*., 2006). In addition, Spo14 is necessary for morphogenesis and polarized growth in response to pheromone and Sec14-independent secretion (Hairfield *et al*., 2001, Harkins *et al*., 2008, Rudge *et al*., 2002). In *Candida albicans*, PLD1 is necessary for the dimorphic transition and full virulence (McLain & Dolan, 1997, Dolan *et al*., 2004). In filamentous fungi, the role of PLDs to date has only been analysed in *Aspergillus fumigatus* and *Fusarium graminearum*. *A. fumigatus* has three PLD isoforms, one of which is required for internalization of the fungus into host epithelial cells and for full virulence in certain immunosuppressed mice (Li *et al*., 2012). *F. graminearum*, also has three *pld* genes, but only one, Fgpld1, is crucial for vegetative development and host virulence. Deletion of *Fgpld1* resulted in reduced colony size, defects in sporulation and sexual development, decreased production of the mycotoxin deoxynivalenol (DON), and reduced virulence on wheat (Ding *et al*., 2017). The molecular mechanisms behind these phenotypic changes however, remain unclear.

The genome of *E. festucae* encodes two putative PC-hydrolysing PLD genes, which we have named *pldA* and *pldB*. The aim of this study was to examine the role of these two proteins, and their orthologs from the model organism *Neurospora crassa*, in hyphal morphogenesis, growth and development in axenic culture, and to test if they are required for establishment of a mutualistic symbiotic association between *E. festucae* and *L. perenne*.

## Results

*The* E. festucae *genome encodes two putative phosphatidylcholine-hydrolysing phospholipase D enzymes*

To identify PLD-encoding genes in the *E. festucae* Fl1 genome, a tBLASTn analysis was performed using *S. cerevisiae* Spo14 as a query sequence. This search identified two putative homologs: EfM3.055250 (E-value of 1e-89) and EfM3.032570 (E-value of 0), hereafter referred to as *pldA* and *pldB*, respectively. An analysis of previous transcriptome datasets showed that both genes are moderately expressed in axenic culture (*pldA*: 10.39 RPMK, *pldB*: 35.96 RPMK) (Hassing *et al*., 2019). The expression of *pldB* is downregulated two-fold *in planta* relative to axenic culture, while the expression of *pldA* is not significantly different (Hassing *et al*., 2019). PldA and PldB are 877 and 1,713 amino acid residues in length, respectively (Fig. 1A) and share 39.8% amino acid sequence identity. Both sequences contain four conserved domains characteristic of PLDs, including the HxKx_4_D(x_6_GSxN) (HKD) motif in domains II and IV (Ponting & Kerr, 1996, Koonin, 1996), necessary for phosphatidyltransferase activity (Sung *et al*., 1997, Liscovitch *et al*., 1999, Liscovitch *et al*., 2000). PldB in addition contains a sequence homologous to a PtdIns[4,5]P_2_-binding polybasic motif located between domains II and III (Sciorra *et al*., 1999, Sciorra *et al*., 2002) (Fig. 1A). Analysis with InterproScan (Jones *et al*., 2014) and SMART (Letunic *et al*., 2015) identified slightly overlapping, N-terminal phosphoinositide-binding PX and PH-domains in PldB but not in PldA. Both domains are present in mammalian and fungal PLDs, but only in the Pld*ζ* group of PLDs found in plants (Selvy *et al*., 2011).

**Fig. 1.**
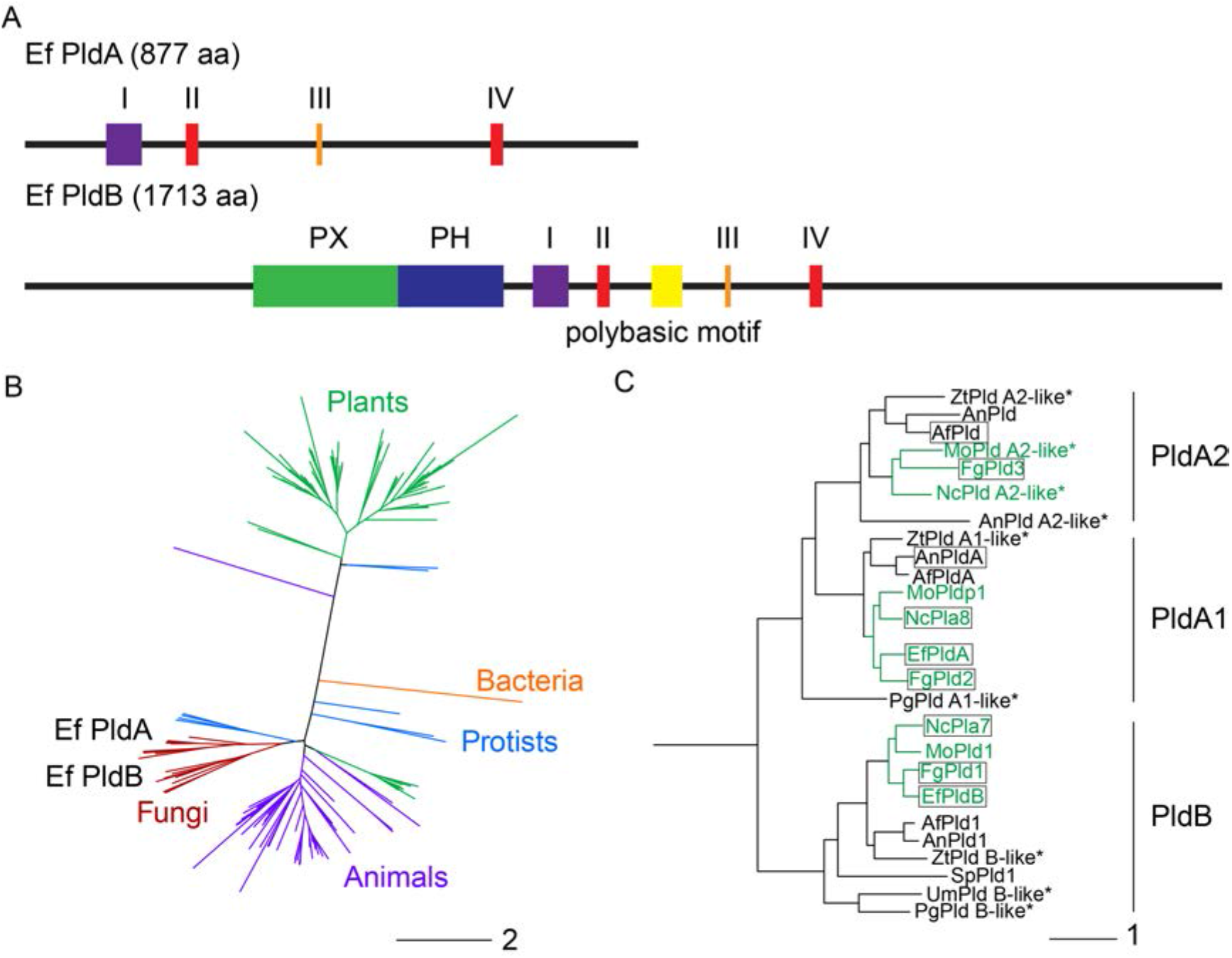
Structure of *E. festucae* PldA and PldB and phylogenetic analysis of phospholipase D in fungi. (A) Domain structure of PldA and PldB, annotated according to InterProScan (Jones *et al*., 2014), SMART (Letunic *et al*., 2015), and the publications of (Sung *et al*., 1997) and (Sciorra *et al*., 1999, Sciorra *et al*., 2002). Green box: PX (phox homology) domain; blue box: PH (pleckstrin homology) domain; purple box: motif I; red box: motif II and IV; orange box: motif III; yellow box: polybasic motif; aa: amino acids. (B) Phylogenetic analysis of the distribution of phospholipase Ds (PLDs) making use of all sequences available on the Esembl genome browser (Zerbino *et al*., 2018). Bar indicates aa substitutions per site. (C) Phylogenetic analysis of PLDs in filamentous fungi with sequences highlighted in green belonging to the Sordariomycetes. Boxed proteins have previously been characterised. Proteins marked with * are described as “putative/hypothetical Pld” in NCBI databases and have therefore been renamed for clarity. Unique identifiers can be found below. *Ef*: *Epichloë festucae* (EfPldB: EfM3.032570, EfPldA: EfM3.055250); *Fg*: *Fusarium graminearum* (FgPld1: XP_011318823.1, FgPld2: XP_011317835.1, FgPld3: XP_011324813.1); *Mo*: *Magnaporthe oryzae* (MoPld1: XP_003717990.1, MoPldp1 XP_003712119.1, MoPld A2-like: XP_003712056.1*); *Nc*: *Neurospora crassa* (NcPLA-7: XP_957594.3, NcPLA-8: XP_001728077.1, NcPld A2-like: XP_962376.3*); *Af*: *Aspergillus fumigatus* (AfPld1: KEY78740.1, AfPldA; XP_748951.1, AfPldA2 like: XP_755989.2); *An: Aspergillus nidulans* (AnPld1: ANIA_10413, AnPldA; ANIA_06712, AnPld: ANIA_02586, AnPld A2-like: ANIA_07334*); *Zt: Zymoseptoria triciti* (ZtPldB1: XP_003857513*, ZtpldA_like: XP_003848870*, ZtPldA2_like: XP_003855368*); *Pg: Puccinia graminis* (PgPld B-like XP_003322083.2*, PgPldA_like: XP_003330640.2*); *Sp: Schizosaccharomyces pombe* (SpPld1: CBE61318.1); *Um: Ustilago maydis* (UmPld B-like: KIS71943*). Bar indicates aa substitutions per site.

Phylogenetic analysis of all PLD sequences available at Ensembl (Zerbino *et al*., 2018), revealed a separation between plant and animal/fungal PLDs, presumably due to the presence of the different protein domains. Using only selected sequences we observed a separation between PldB-like and PldA-like PLDs within the fungal kingdom, demonstrating that both isoforms are conserved among filamentous fungi. All analysed fungal genomes with the exception of *E. festucae*, contained a third PLD isoform, which separated as a subgroup of PldA-like PLDs, here named PldA2-like (Figs. 1B & 1C).

An amino acid sequence alignment of PldA and PldB with the corresponding homologs from *N. crassa, M. oryzae, F. graminearum* and *A. fumigatus*, revealed that PldA and PldB homologs have high sequence identity (an average of 68% and 58%, respectively) (Fig. S1A). For PldB, the sequences were most similar within the catalytic core (Fig. S1B).

### PldB is required for hyphal morphogenesis, growth and fusion

To investigate whether *E. festucae* PldA and PldB are required for hyphal morphogenesis and growth, each gene was individually deleted from the WT background by gene replacement. Three hygromycin-resistant Δ*pldA* strains (#T47, #T49, #T82) and three geneticin-resistant Δ*pldB* strains (#T26, #T50, #T87) were identified (Fig. S2).

Although deletion of *pldA* had no obvious effect on colony growth and morphology, deletion of *pldB* led to a significant reduction in colony size and an increase of aerial hyphae (Fig. 2). Analysis by differential interference contrast (DIC) light microscopy confirmed that the hyphal morphology of Δ*pldA* strains was indistinguishable from WT (Fig. 2B-F). In contrast, Δ*pldB* strains frequently had swollen and highly vacuolated hyphae and hyphal tips (Fig. 2F). Staining of the colony with the chitin stain CFW revealed occasional accumulation of the stain in the cytoplasm of hyphae, which is indicative of a defect in the recycling of cell wall components (Higuchi *et al*., 2009). Intra-hyphal hyphae (IHH) were also occasionally detected. Although both strains formed hyphal bundles and coils like WT, no cell fusion could be observed in the Δ*pldB* strains (Figs. 2E & S3). To verify this cell fusion defect, Δ*pldB* strains expressing either GFP or mCherry under the regulation of a constitutive promoter, were grown in close proximity to one another and analysed for cytoplasmic mixing (Becker *et al*., 2015). WT hyphae with both GFP and mCherry fluorescence were frequently observed, but in the Δ*pldB* strains the hyphae had either GFP or mCherry fluorescence but not both (Fig. 3). Addition of PA to the growth medium rescued the Δ*pldB* hyphal cell fusion defect (Figs. 4 & S4), although this chemical complementation with PA was concentration dependent. However, PA was unable to rescue the hyphal morphology and growth defect of Δ*pldB*. Introduction of the WT *pldB* allele into Δ*pldB* did complement all observed axenic culture phenotypes (Figs. 2 & S3) confirming that the defects were due to the specific deletion of the *pldB* gene.

**Fig. 2.**
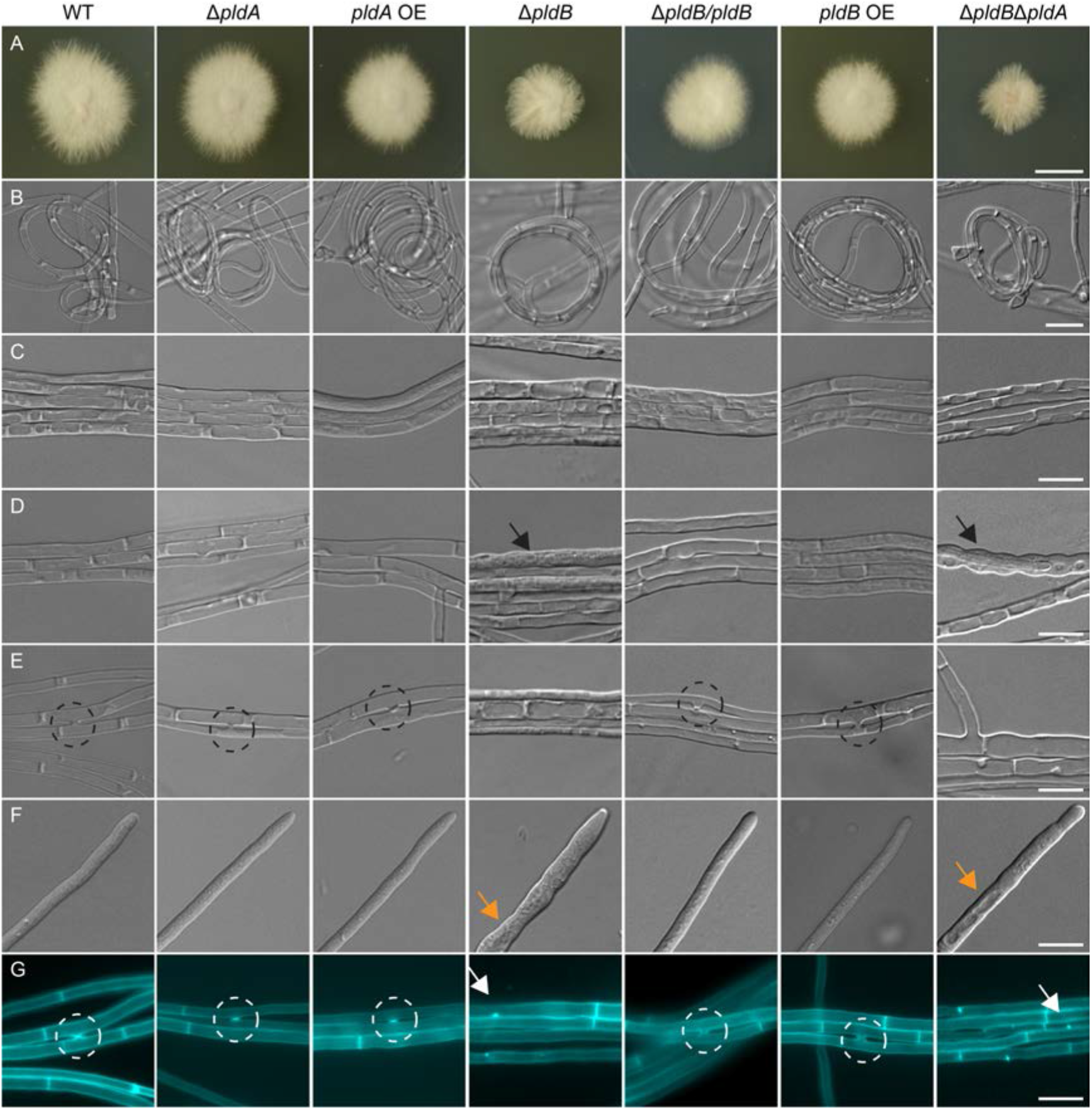
Culture phenotype of wild-type *E. festucae* and Δ*pldA* and Δ*pldB* strains. The phenotypes of the wild-type (WT) strain and three independent mutants of each genotype were analysed. The images shown of Δ*pldA*#T47, *pldA* overexpression strains (OE), OE#T4, Δ*pldB*#T87, Δ*pldB*T#87/*pldB#*T4, *pldB* OE#T21, Δ*pldB*Δ*pldA*#T3, are representative of all strains analysed. (A) Representative image of the whole colony morphology after seven days on 2.4% potato-dextrose agar. Bar=1 cm; (B-G) Representative differential interference contrast microscopy images of hyphal morphology after seven days of incubation on 1.5% H_2_O agar; Bar=10 µm; (B) hyphal coil; (C) hyphal bundle; (D) intra-hyphal hyphae; (E) hyphal fusion; (F) hyphal tip; (G) hyphae stained with Calcofluor white (CFW) to examine cell wall composition. Black arrows: intra-hyphal hyphae; dashed circles: hyphal fusion; orange arrows: vacuolated hyphal tips; white arrows: cytoplasmic CFW accumulations.

**Fig. 3.**
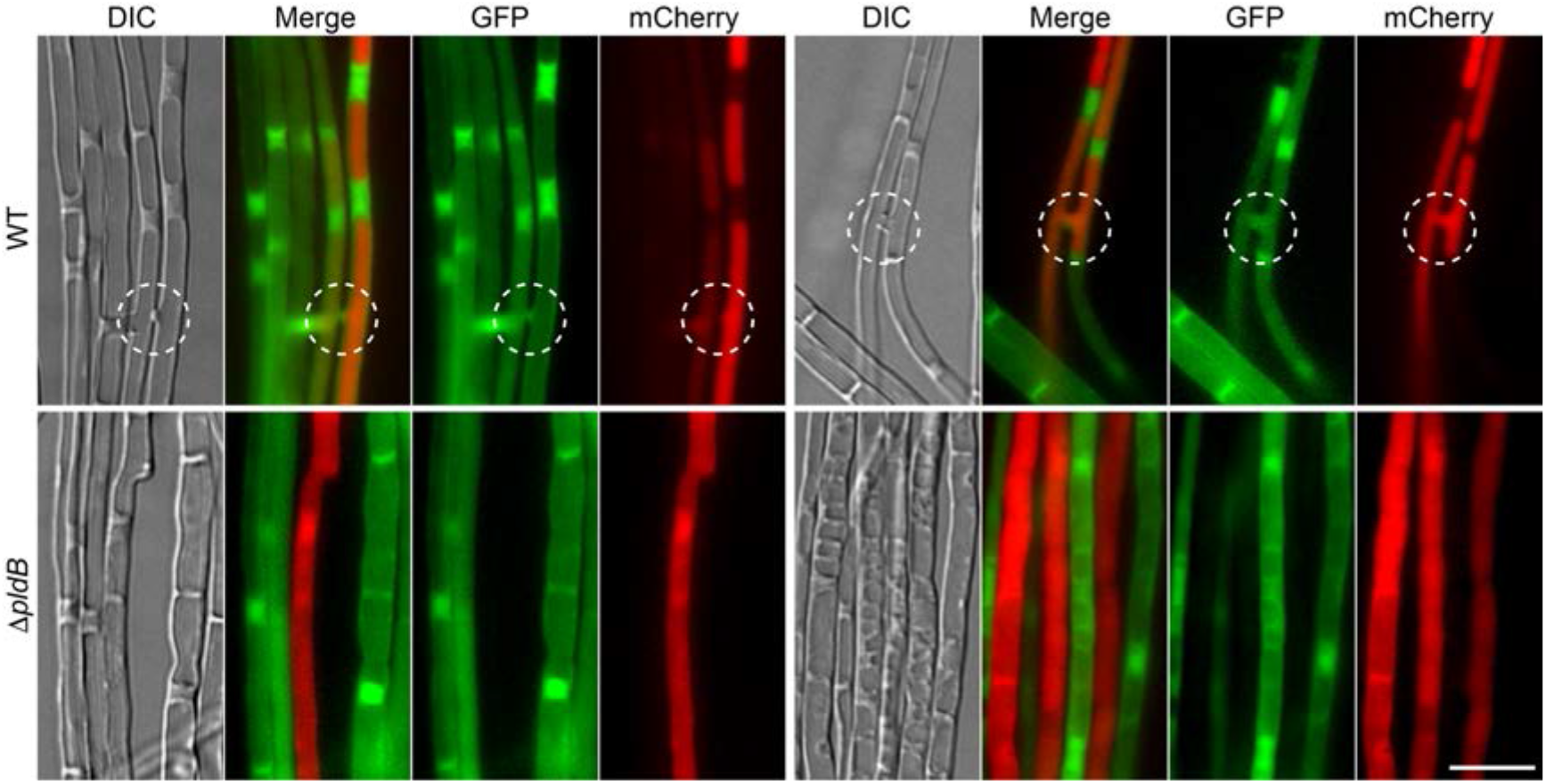
Cytoplasmic mixing of wild-type and Δ*pldB* strains of *E. festucae* expressing cytoplasmic GFP and mCherry. Wild-type (WT) strains expressing cytoplasmic GFP (pBH28) and mCherry (pCE126) (WT GFP#T1/WT mCherry#T1) and Δ*pldB* strains (Δ*pldB*#T26 GFP#T1/Δ*pldB*#T26 mCherry T2) expressing cytoplasmic GFP or mCherry were co-cultured on 1.5% H_2_O agar for 7 days. Strains were examined for hyphae expressing both GFP and mCherry to indicate hyphal fusion followed by cytoplasmic mixing. White, dashed circles: hyphal fusion. Bar=10 µm.

**Fig. 4.**
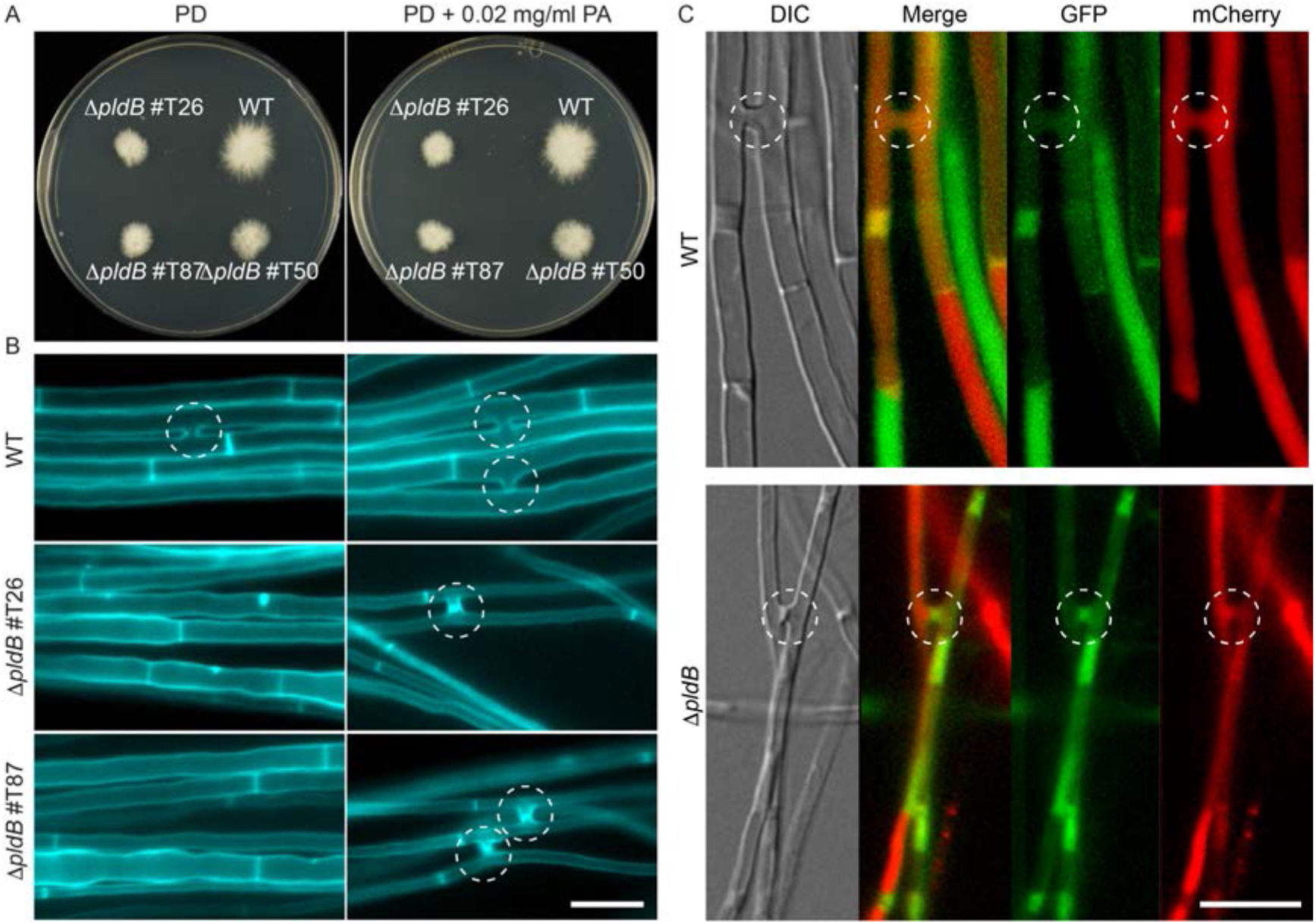
Exogenous addition of phosphatidic acid restores hyphal cell to cell fusion in *E. festucae* Δ*pldB* strains. (A) Wild-type (WT) and Δ*pldB* strains were grown on potato dextrose (PD) agar plates overlaid with 7 ml of PD agar containing 0.02 mg/ml phosphatidic acid (PA) and cultured for 7 days. (B) WT and Δ*pldB* strains were grown on H_2_O-agar plates overlaid with 7 ml of H_2_O-agar containing 0.02 mg/ml PA, cultured for 7 days, and stained with Calcofluor white before analysis to highlight sites of cell fusion. Dashed circles: hyphal fusion. Bar=10 µm. (C) Wild-type (WT) strains expressing cytoplasmic GFP (pBH28) and mCherry (pCE126) (WT GFP#T6/WT mCherry#T2) and Δ*pldB* strains (Δ*pldB*#T26 GFP#T1/Δ*pldB*#T26 mCherry T2) expressing cytoplasmic GFP or mCherry were co-cultured on 1.5% H_2_O-agar plates overlaid with 7 ml of H_2_O-agar containing 0.02 mg/ml PA for 7 days. Strains were examined for hyphae expressing both GFP and mCherry to indicate hyphal fusion followed by cytoplasmic mixing. White, dashed circles: hyphal fusion. Bar=10 µm.

To determine whether there was any functional redundancy between PldA and PldB, double deletion strains were generated by deleting *pldA* in the Δ*pldB*#T26 and #T87 backgrounds. Two strains with a *pldA* deletion were obtained (Δ*pldB*Δ*pldA*#T3 and Δ*pldB*Δ*pldA*#T5) (Fig. S2) and both had axenic culture phenotypes comparable to the single Δ*pldB* strains (Fig. 2).

To determine whether overexpression of these genes would influence hyphal morphology and growth, constructs with *pldA* and *pldB* under the control of the *A. nidulans gpdA* promoter were transformed into the WT strain. Strains with a range of overexpression values (*pldA* OE strains: #T3, #T4, #T9, *pldB* OE strains: #T6, #T21, #T22; Fig. S5), as determined by RT-qPCR, were grown alongside WT in axenic culture but found to have no obvious phenotype differences (Fig. 2).

*The PldB ortholog PLA-7 is required for hyphal fusion and spore production in* N. crassa

To test if the role of PldB in vegetative cell fusion is conserved in other fungi, mutants carrying a deletion of the genes homologous to PldA (NCU10400, PLA-8) and PldB (NCU03955, PLA-7) were obtained from the *N. crassa* gene knock-out collection. While strains are usually available as homokaryotic isolates, the Δ*pla-7* mutant was recorded in the collection as heterokaryotic. This indicated that *pla-7* is either essential or required for successful sexual crossing. Single spore isolation resulted in homokaryotic strains, indicating that *pla-7* is not essential.

In *N. crassa* vegetative cell fusion occurs at two stages of mycelial development: at an early stage when germinating conidia (germlings) home towards each other and undergo consecutive fusion events to form supracellular structures, and at a later stage, within mature colonies, when hyphal branches fuse to form cross connections that increase the interconnectivity of the mycelium. The cell fusion phenotype of germinating conidia or mature hyphae of the Δ*pla-8* mutant was indistinguishable from the WT strain. In contrast, germination of Δ*pla-7* mutant spores, was significantly delayed, and germling homing and fusion was completely absent. Addition of PA, using the same concentration range as used for *E. festucae*, failed to rescue the germling fusion phenotype of *N. crassa Δpla-7* (Fig. S6). No hyphal fusion was detected in mature Δ*pla-7* mutant colonies (Fig. 5). Together these observations indicate that PLA-7 has a conserved essential function in vegetative cell fusion.

**Fig. 5.**
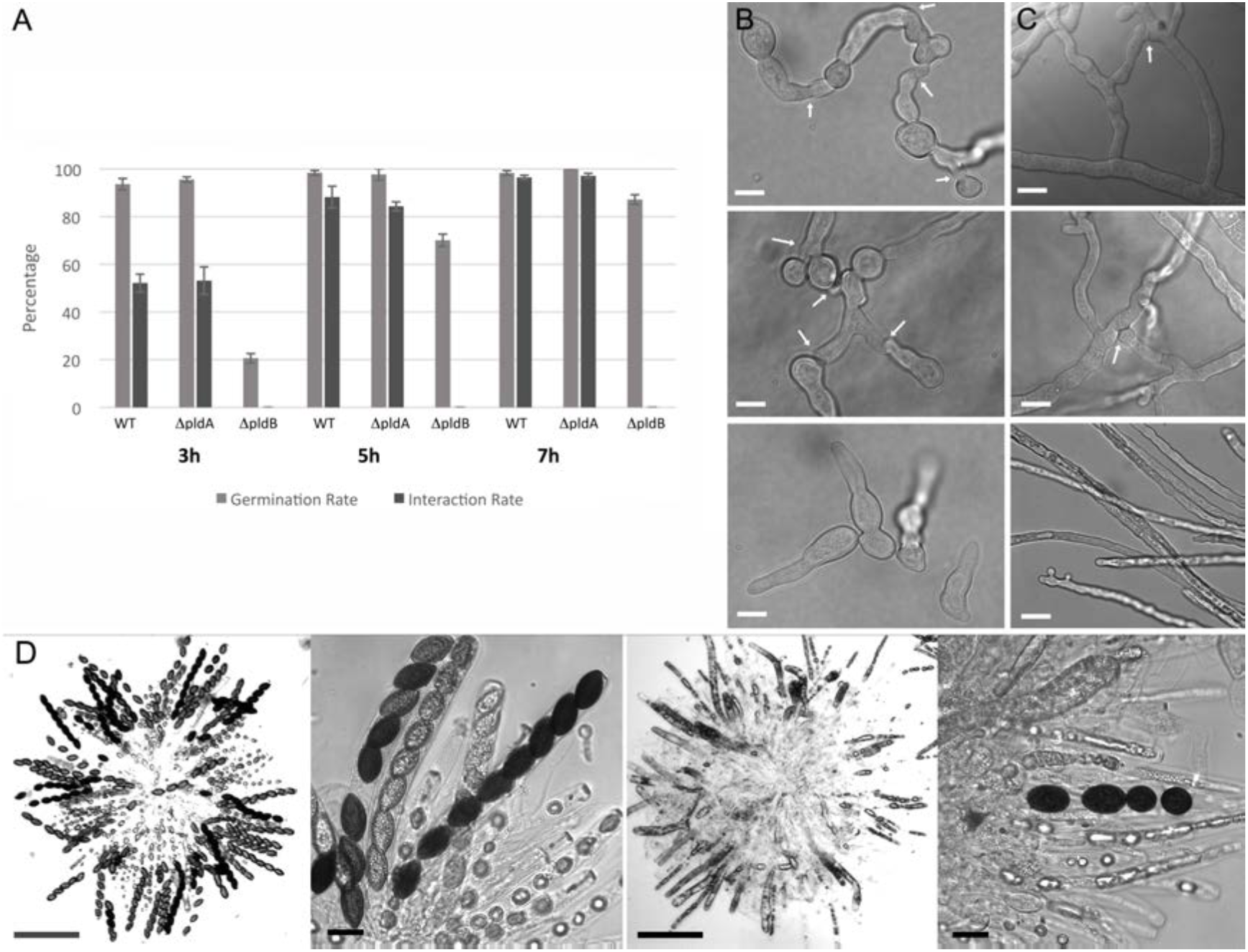
PLA-7 is essential for cell fusion and female fertility in *N. crassa*. (A) Conidia of the wild-type WT) reference strain or the Δ*pla-7* and Δ*pla-8* mutants were cultured on Vogeĺs minimal medium. Spore germination and germling fusion were quantified at the indicated time points (percentage of all spores/spore germlings). (B) After 5 hours of incubation WT and Δ*pla-8* spore germlings form a supracellular network via cell-cell fusion (top and middle image, respectively. Arrows indicate fusion points. Germling fusion is absent in the Δ*pla-7* mutant (bottom image). Bar = 5 µm. (C) Within mature wild-type colonies, hyphal fusion (arrow) increases the interconnectedness. Hyphae in Δ*pla-7* colonies fail to form a network and grow in a straight fashion. Bar=10 µm. (D) Rosettes of asci were squeezed out of 2-week-old fruiting bodies. While the majority of asci obtained from the wild-type reference cross contains eight black ascospores (left and center left image), crosses fertilized with Δ*pldB* conidia produced only low numbers of ascospores and many asci contained aberrant structures (right and center right image). Bar=200 μm (left and center right image) and 20 μm (center left and right image).

Unlike *E. festucae, N. crassa* readily forms sexual structures in axenic culture thereby enabling the role of PLA-7 in sexual development to be analysed by crossing Δ*pla-7* with the WT strain. While the Δ*pla-7* strain failed to produce female reproductive structures, crosses between WT as a female and Δ*pla-7* as the male resulted in ascospores. However, ascus formation was aberrant and sometimes asci contained less than the usual eight ascospores. Overall WT/Δ*pla-7* crosses produced significantly fewer spores than WT/WT pairings (Fig. 5). In addition, the fraction of mutant progeny was just 17%, compared to an expected 50% for a 1:1 segregation of Δ*pla-7* to *pla-7^+^*in this cross.

### PldB localises to septa and the plasma membrane

PLDs analysed to date localise to various cellular compartments, including the Golgi apparatus, vesicles, endosomes, the plasma membrane (PM), and the cytosol (Colley *et al*., 1997, Hughes & Parker, 2001, Du *et al*., 2003, Selvy *et al*., 2011). To investigate the localization of *E. festucae* PldA and PldB, N-terminal GFP translational fusion constructs were generated and transformed into WT protoplasts. Expression of fusion proteins of the expected size was verified by western blot analysis (Fig. S7). To confirm the functionality of the PldB localization constructs, the constructs were transformed into Δ*pldB* strains, which restored WT-like growth in axenic culture (Fig. S8).

The PldA-GFP fusion protein localized to the cytoplasm in hyphae of different developmental stages, a result comparable to that of strains expressing cytosolic GFP driven by the same promoter (Fig. 6). The PldB-GFP fusion protein was also found to localise to the cytoplasm of hyphae of all ages. However, in young hyphal bundles, as well as in mature hyphae, the signal was also detected in septa and occasionally associated with the PM (Fig. 6), a result indicative of a transient localization of PldB to the membrane (Selvy *et al*., 2011).

**Fig. 6.**
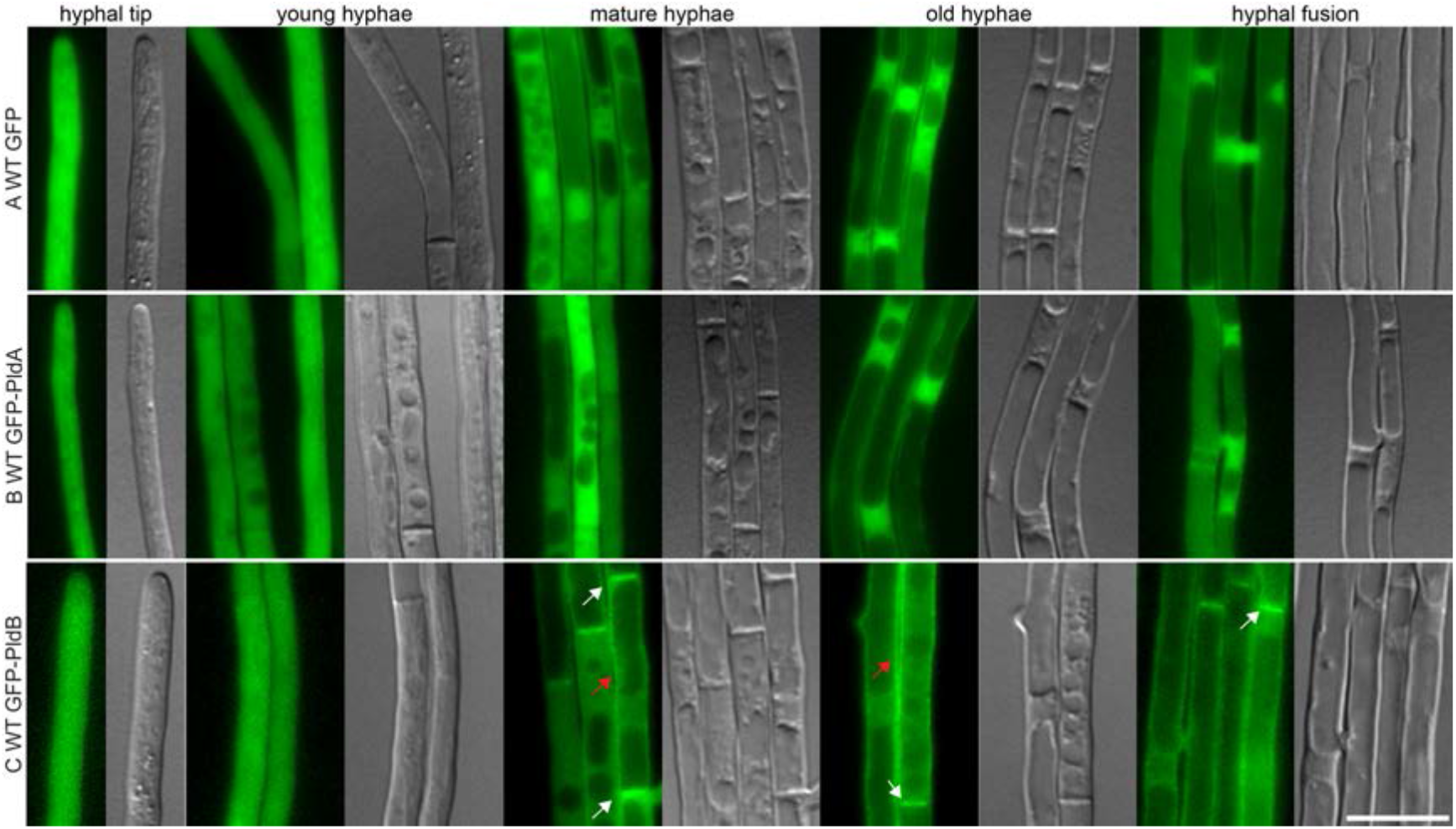
Localization of GFP-PldA and GFP-PldB in *E. festucae* in axenic culture. (A-C) Wild-type (WT) strains expressing either (a) cytoplasmic GFP (pBH28, #T6), (B) GFP-PldA (pBH37, #T4), or (C) GFP-PldB (pBH54, #T14). Fusion proteins were analysed in axenic culture by fluorescence microscopy after approx. five days of incubation. The images shown here are representative of multiple transformants containing these constructs. White arrows: Septa localization of GFP-PldB; Red arrows: PM localization of GFP-PldB. Bar=10 µm.

*PldB is crucial for the* E. festucae-L. perenne *symbiotic interaction and expressoria formation*

To analyse the role of PldA and PldB in the interaction of *E. festucae* with its host, deletion and overexpression (OE) strains were inoculated into seedlings of *L. perenne* and the phenotype of the mature plants examined at 10 weeks post-infection. Plants infected with Δ*pldA*, or *pldA* and *pldB* OE strains, were indistinguishable from those infected with WT. In contrast, plants infected with Δ*pldB* strains were severely stunted, with tiller lengths significantly less than that of plants infected with WT (Fig. 7). While the tiller number of plants infected with Δ*pldB*#T87 was significantly greater than WT, there were no significant differences in tiller number for the other two, independent Δ*pldB* mutants. Introduction of a *pldB* WT allele into the Δ*pldB* strains restored the WT plant interaction phenotype. For the Δ*pldB*Δ*pldA* double deletion strain tiller number was increased and tiller length reduced, compared to WT (Fig. 7).

**Fig. 7.**
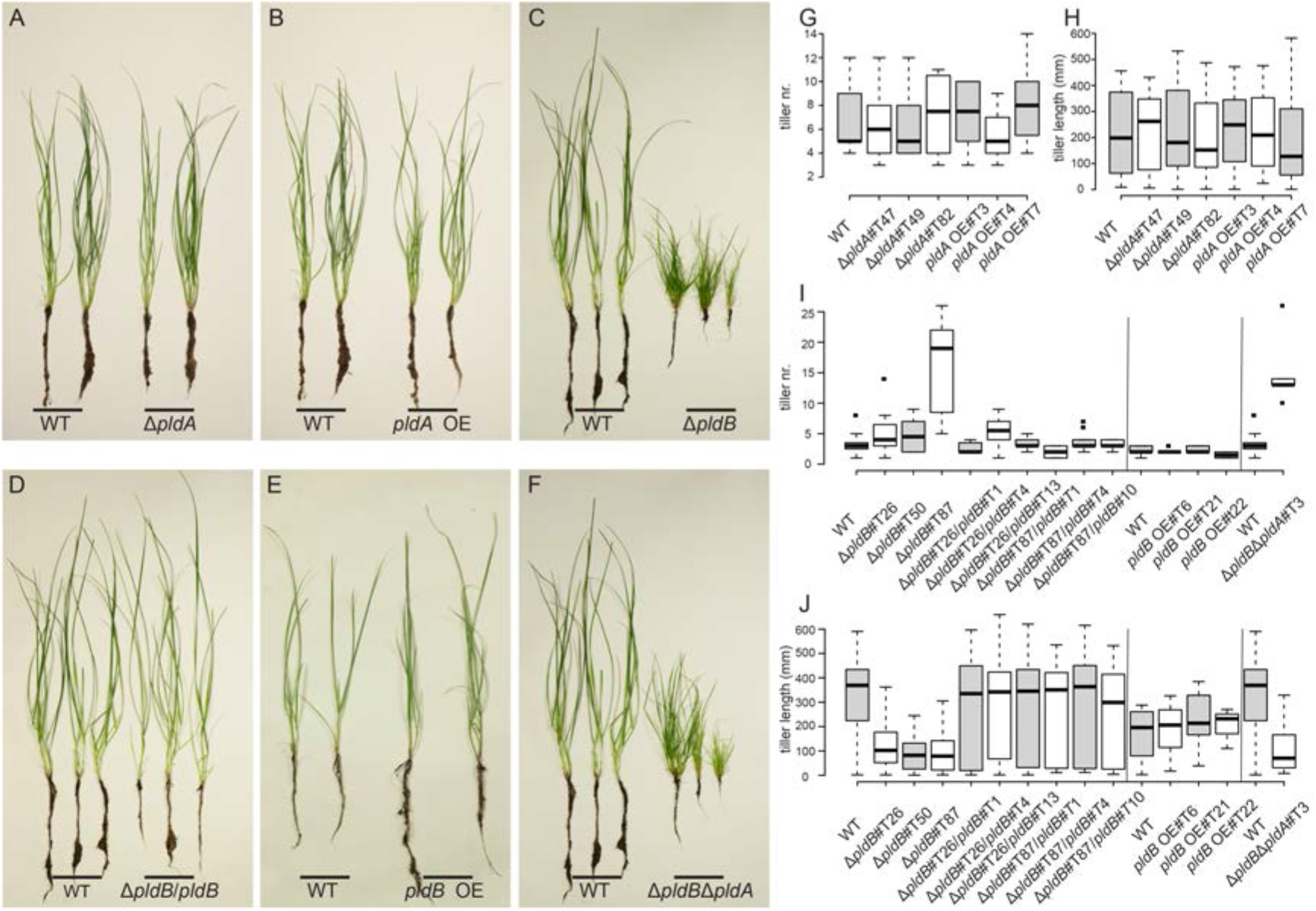
Plant phenotype of *L. perenne* infected with wild-type and Δ*pldA* and Δ*pldB* strains. Plants infected with the wild-type (WT) strain and three independent transformants for each mutation, with the exception of the double deletion strain, were analysed by microscopy. The images shown here are representative of plants infected with all analysed strains. Strains depicted here are the deletion mutants Δ*pldA*#T47, Δ*pldB*#T87, Δ*pldBΔpldA*#T3, and Δ*pldB*#T87/*pldB*#T4 and overexpression (OE) strains *pldA* OE#T4 and *pldB* OE#T21. (A-F) Plant phenotype of WT and mutant-infected plants 8 weeks post-planting (10 weeks post-inoculation); (G-J) boxplot representation of tiller number (G and I) and tiller length (H and J) of infected plants; (G and H) data of Δ*pldA*-infected plants, WT (n=5), Δ*pldA* (n=6/6/4), *pldA* OE (n=6/8/8); (I and J) data of Δ*pldB*-infected plants, WT (n=11/6/11), Δ*pldB* (n=8/6/7), *pldB* complementation (n=11/10/10/9/13/10), *pldB* OE (n=5/6/2) and Δ*pldBΔpldA* double deletion (n=7). Note that the Δ*pldB* and complementation strains and overexpression strains were analysed in different experiments and therefore can only be compared to the WT of the same experiment. This is indicated by black lines in the graphs. One-way ANOVAs were used to test for differences in plant phenotypes between WT and mutant strains. In each case, the ANOVA was fitted with R, and a Bonferroni correction was applied to all p-values to account for multiple testing. Error bars represent the standard deviation. Only Δ*pldB* and Δ*pldBΔpldA* strains had a significantly reduced tiller length (Δ*pldB* #T26: P=1.01E-7; #T50: P=1.22E-8; #T87: P=2.04E-13; Δ*pldBΔpldA*#T3: P=1.09E-12) and only Δ*pldB*#T87 and Δ*pldBΔpldA*#T3 had an increased tiller number (P=5.57E-12; P=1.95E-10).

To examine hyphal growth *in planta*, pseudostem cross sections were fixed and embedded in resin, stained with toluidine blue, and the number of hyphae per intercellular space determined by light microscopy. Plants infected with the Δ*pldB* strains had a significantly greater number (P < 1E-10) of hyphae per intercellular space (3.02) than plants infected with WT strain (1.28). The number of hyphae per intercellular space for plants infected with Δ*pldA* (1.27), *pldA* OE (1.37) and *pldB* OE (1.27) strains were not significantly different to WT (Figs 8A & S9). TEM ultrastructure analysis of pseudostem cross sections revealed that the Δ*pldB* and Δ*pldB*Δ*pldA* double deletion strains occasionally colonise the vascular bundles, a phenotype seldom observed for WT (Fig. 8B-E). Hyphae from either of these two mutants were often less electron-dense or completely vacuolated and frequently formed IHH when growing in the intercellular space (Fig. 8B-E). Another distinct phenotype of both mutants was their apparent reduced ability to degrade the plant cuticle compared to WT and the Δ*pldA* strains, leading to a proliferation of hyphae immediately below the cuticle surface (Fig. 8).

**Fig. 8.**
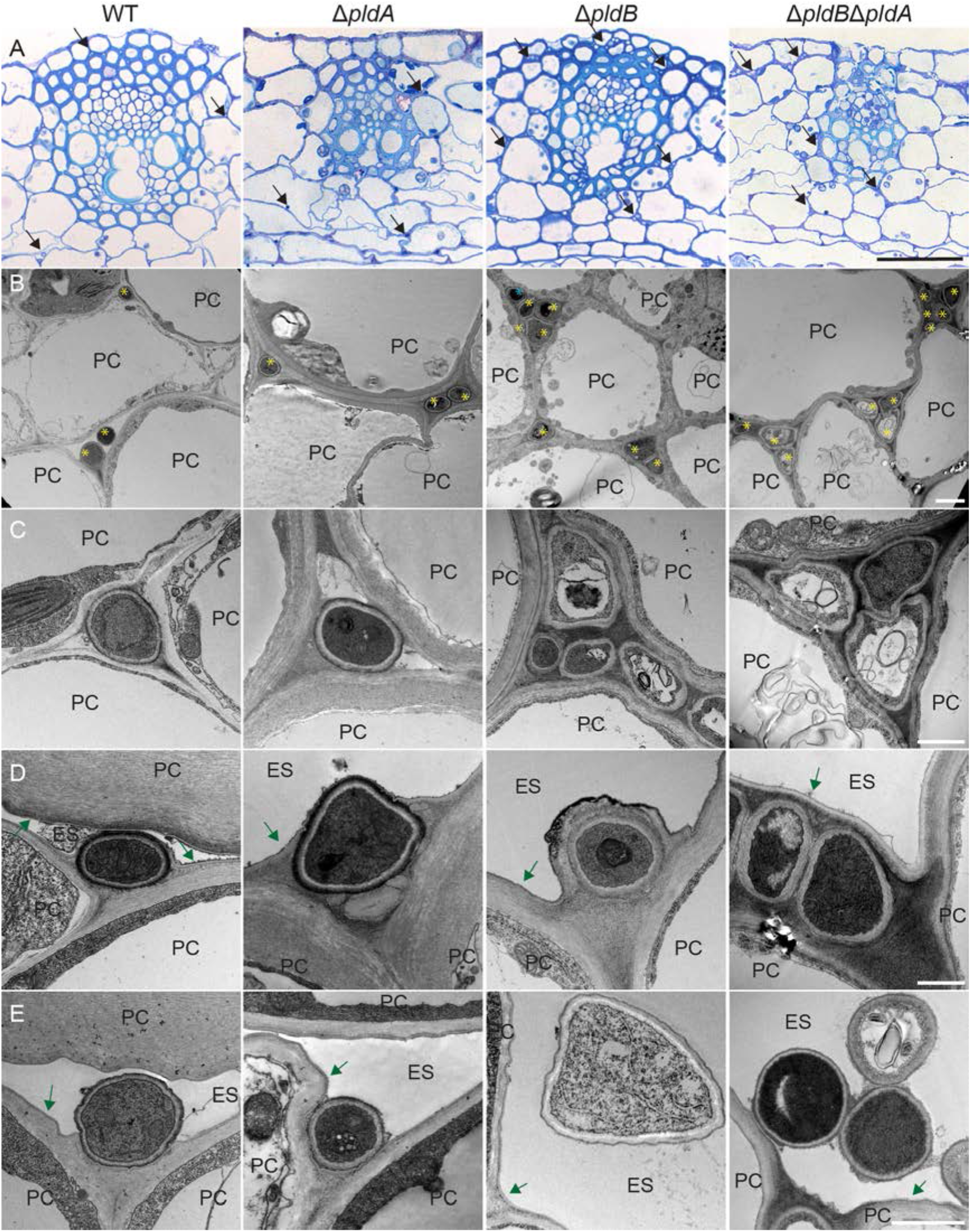
Light microscopy and transmission electron microscopy of *L. perenne* pseudostem cross sections infected with wild-type, Δ*pldA* and Δ*pldB* strains of *E. festucae*. Plants infected with wild-type (WT) and three independent Δ*pldA* and Δ*pldB* strains were analysed. Here only Δ*pldA*#T47, Δ*pldB*#T26 and Δ*pldBΔpldA*#T3 infected plants are shown as representative of all analysed strains. As plants infected with *pldA* and *pldB* OE strains were asymptomatic no TEM analysis was performed. (A) Vascular bundles of WT and mutant-infected plants stained with toluidine blue and imaged by light microscopy. Bar=50 µm. (B) Representative TEM micrograph of growth of WT and mutant strains in infected plants. Bar=2 µm. (C) Representative TEM micrographs of the hyphal structure of WT and mutant strains in infected plants. Bar=1 µm. (D) Representative TEM micrographs of subcuticular hypha of WT and mutant strains in infected plants. Bar=1 µm. Note the cuticle degradation on top of the WT and Δ*pldA* hyphae. (E) Representative TEM micrographs of epiphyllous hyphae of WT and mutant strains in infected plants. Bar=1 µm. Black arrow/yellow star: hyphae, blue star: intra-hyphal hypha, green arrow: cuticle, PC: plant cell. ES: extracellular space.

To further investigate the cellular phenotype of the host interaction, longitudinal sections of the pseudostem were infiltrated with WGA-AF488, which binds chitin, and aniline blue, which binds β-glucans, and analysed by confocal laser scanning microscopy. The Δ*pldA*, and *pldA* and *pldB* OE strains had a hyphal morphology and restricted growth phenotype *in planta* indistinguishable from WT (Fig. 9). In contrast, growth of the Δ*pldB* strains was unrestricted with multiple hyphae in the intercellular space, which formed bundles and multicellular lobed structures. Furthermore, Δ*pldB* strains were unable to efficiently form expressoria, resulting in a proliferation of hyphae immediately below the cuticle. However, these mutant hyphae were still able to occasionally breach the cuticle to grow as epiphytic hyphae on the surface of the leaf (Fig. 9). The Δ*pldBΔpldA* double mutant had a very similar hyphal morphology and growth phenotype *in planta* to the Δ*pldB* strain. As expected, the *in planta* hyphal morphology and growth defects were rescued by introduction of a WT allele of *pldB*. (Fig. 9).

**Fig. 9.**
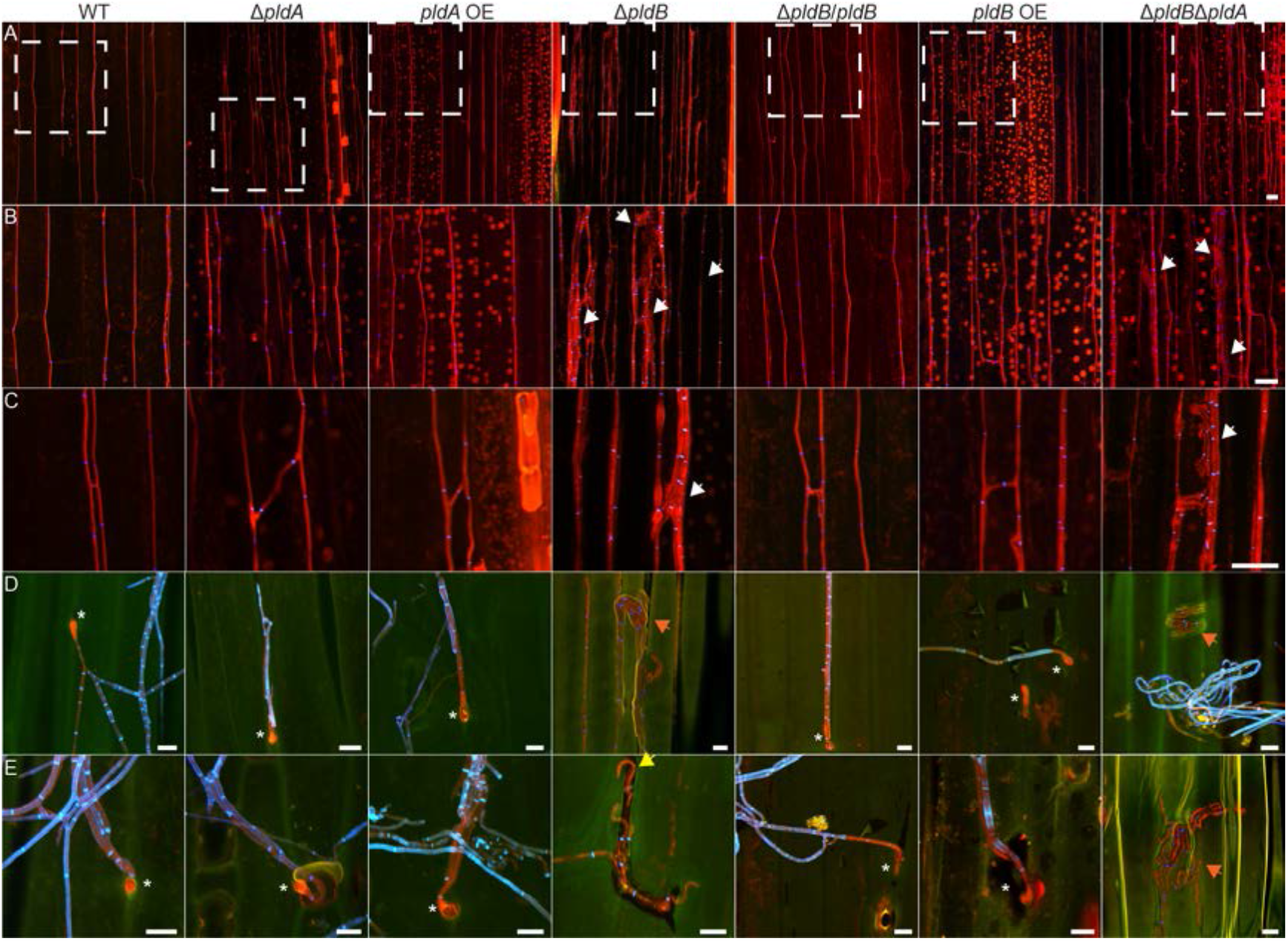
Confocal depth image series of the cellular phenotype of *L. perenne* infected with *E. festucae* wild-type, Δ*pldA* and Δ*pldB* strains. Three independent transformants of each described genotype were analysed. Images of only one strain are shown, as they are representative of all strains analysed. Strains depicted here are the deletion strains Δ*pldA*#T47, Δ*pldB*#T87 and Δ*pldBΔpldA*#T3, the complementation strain Δ*pldB*#T87/*pldB*#T4 and the overexpression (OE) strains *pldA* OE#T4 and *pldB* OE#T21. Infected *L. perenne* pseudostem samples were stained with WGA-AF488 (chitin-binding) and aniline blue (*β*-glucan-binding) and visualised with confocal laser scanning microscopy. In images showing epiphyllous hyphae (D and E), autofluorescence of the cuticle was captured in green pseudocolor. (A) Representative image (*z*=6 µm) of growth of wild-type (WT), Δ*pldA* (#T47), Δ*pldB* (#T87), *pldA* OE#T4, *pldB* OE#T21, *pldB* complementation (Δ*pldB*#T87/*pldB* C4) and Δ*pldBΔpldA* deletion strains (#T3) *in planta*. (B) Magnified image of the dashed, white-boxed area in (A). (C) Higher magnification image (*z*=4 µm) of representative hyphal branching and fusion *in planta*. (D and E) Representative images of an expressorium formed by WT and mutant strains (*z*=4 µm); subcuticular hyphae retained the staining pattern of endophytic hyphae; white arrow: endophytic hyphal bundles in Δ*pldB* and double deletion strains; yellow arrow: hyphae escaping from below the cuticle through rip in cuticle; orange arrow: subcuticular hyphae, note the green hue on hyphae; white star: expressorium. Bar=10 µm.

### Deletion of pldB, but not pldA, results in delocalization of a phosphatidic acid biosensor from the plasma membrane

To analyse the effect of the *pldA* and *pldB* deletions on the localization of PA in the cell, a PA biosensor was constructed comprising a Spo20 PA-binding domain fused to GFP (Potocký *et al*., 2014). This construct (pBH50) was transformed into WT, Δ*pldA* and Δ*pldB* strains, and the resulting transformants shown to express fusion proteins of the correct molecular weight (Fig. S10). In the WT strain, the PA biosensor localised to the PM and septa in young as well as mature hyphae (Fig. 10). Interestingly, the signal in newly formed hyphal branch tips was more intense than in the tips of hyphae at the colony growth front. PA localization in the Δ*pldA* strains was indistinguishable from WT. However, in the Δ*pldB* strains, the signal was largely cytoplasmic, and localization to the PM and septa was only detected in mature hyphae (Fig. 10A). In addition, the PA biosensor accumulated in an organelle in the cytoplasm of the Δ*pldB* strains that was shown by DAPI staining to be the nucleus (Fig. 10B). To examine the localization of PA in fungal hyphae *in planta*, a WT strain expressing the PA biosensor was inoculated into *L. perenne* seedlings and GFP expression examined in mature plants. GFP fluorescence was detected evenly along the PM and septa of both endophytic as well as epiphytic hyphal tips and mature hyphae, a pattern identical to that observed for PA localization in hyphae in axenic culture (Fig. 10C).

**Fig. 10.**
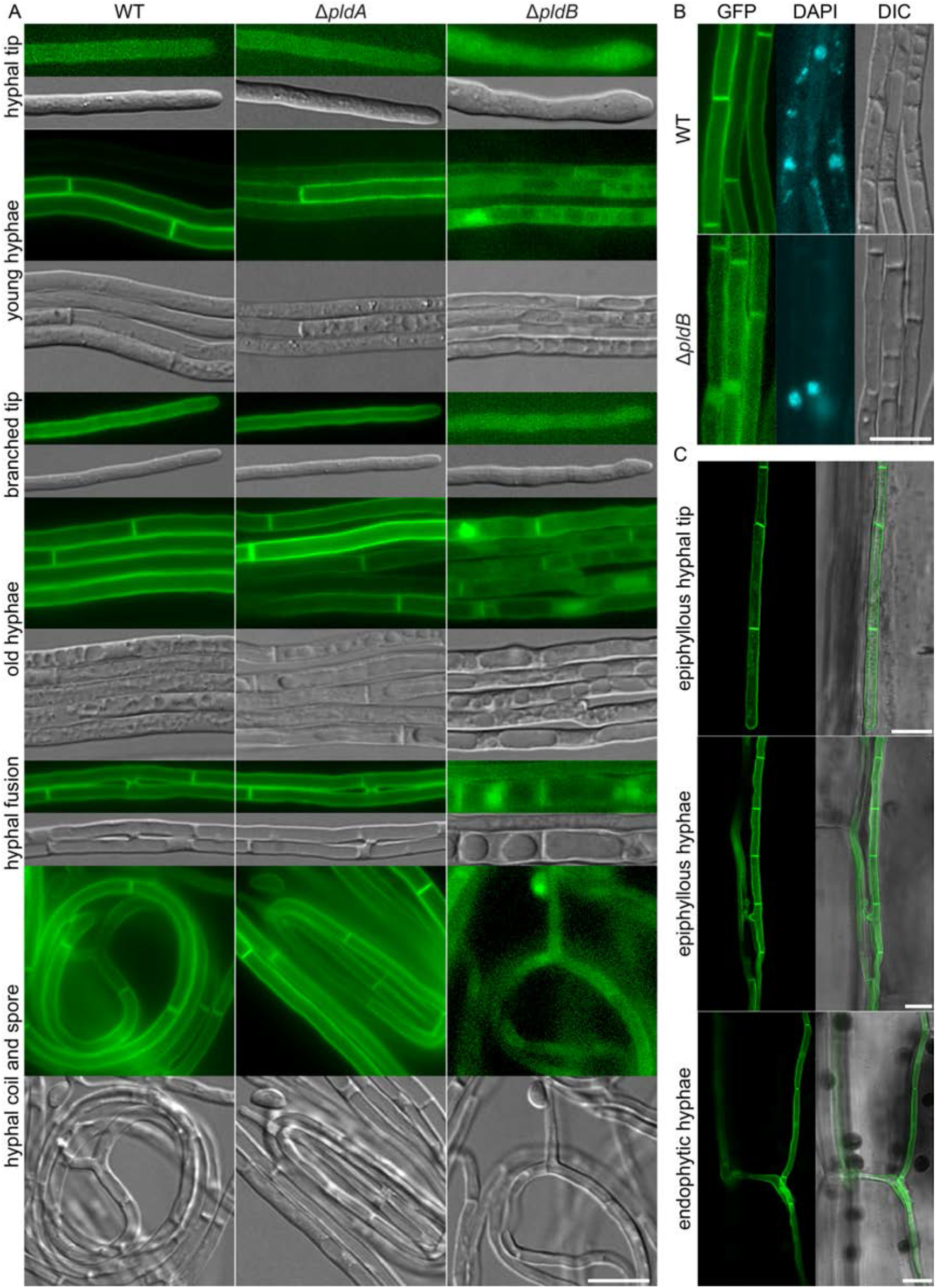
Localization of a phosphatidic acid probe in *E. festucae* wild-type and Δ*pldA* and Δ*pldB* strains growing in axenic culture and *in planta.* Phosphatidic acid biosensor (PAS, pBH50) constructs were transformed into wild-type (WT) and two different Δ*pldA* (#T47 and #T82) and Δ*pldB* (#T26 and #T87) strains, and multiple transformants analysed. Here, the localization in WT GFP-PAS #T4, Δ*pldA*#T82 GFP-PAS#T10 and Δ*pldB*#T26 GFP-PAS#T1 are shown as representative of all strains analysed. (A) Strains expressing the PA biosensor were analysed after approx. 5 days of incubation and hyphae of different ages and developmental stages are shown. (B) WT GFP-PAP and Δ*pldB*#T26 GFP-PAS#T1 were incubated as described and stained with DAPI to visualize nuclei. (C) Localization of the PA biosensor in WT GFP-PAS-infected, mature (>10 weeks) *L. perenne* plants. Representative pictures of the localization in endophytic and epiphyllous hyphae, as well as in epiphyllous hyphal tips, are shown. Bar=10 µm.

### The absence of PldB has no effect on localization of Nox complex components

In mammalian systems PA produced by PLD has been shown to activate the Nox complex by promoting the recruitment of the Nox complex subunits to the membrane as well as by direct interaction of PLD1 or PLD2 with Nox-activating proteins (Regier *et al*., 2000, Kanai *et al*., 2001, Karathanassis *et al*., 2002, Ago *et al*., 2003, Chae *et al*., 2008, Selvy *et al*., 2011, Jang *et al*., 2012). We therefore tested for a direct interaction between PldB and components of the Nox complex (NoxR, BemA, RacA, Cdc24) and other potential regulators (Cdc42, PkcA (EfM3.006500), MssD (EfM3.031950)) by yeast two-hybrid assays. However, PldB did not interact with any of these proteins (Fig. S11). We next examined whether deletion of *pldA* or *pldB* would affect the localization of components of the Nox complex, including the catalytic transmembrane protein NoxA, the regulatory proteins NoxR, BemA, and Cdc24, and the small GTPase RacA using GFP translational fusions of each (Fig. S12) (Takemoto *et al*., 2011). In the WT background all fusion proteins localised as previously reported and this pattern of localization was unchanged in both the Δ*pldA* and Δ*pldB* strains (Fig. S12).

### Absence of PldB results in mislocalized reactive oxygen species production

While the localization of the components of the NoxA complex was not altered, PA produced by PLD has been shown to directly activate Nox complexes in mammalian and plant systems (Karathanassis *et al*., 2002, Taylor *et al*., 2004, Taylor *et al*., 2012, Potocký *et al*., 2012). To investigate the role of PA in the activation of fungal NADPH oxidases, the production and localization of superoxide was analysed in Δ*pldA* and Δ*pldB* strains in culture, using NBT and a ROS-ID^®^ Superoxide kit.

When WT colonies were stained with NBT, a dark blue formazan precipitate was formed that was predominantly localised to hyphal tips, with only a small number (∼5%) having either an aberrant staining pattern or no detectable signal (Fig. 11). A similar pattern of staining was observed for the Δ*pldA* strains, as well as the *pldA* and *pldB* OE strains. In comparison, this aberrant pattern of NBT staining of tips was increased to approx. 20% in Δ*noxA*, and approx. 40% in Δ*pldB*. Normal tip staining was restored in the Δ*pldB*/*pldB* complementation strains (Fig. 11).

**Fig. 11.**
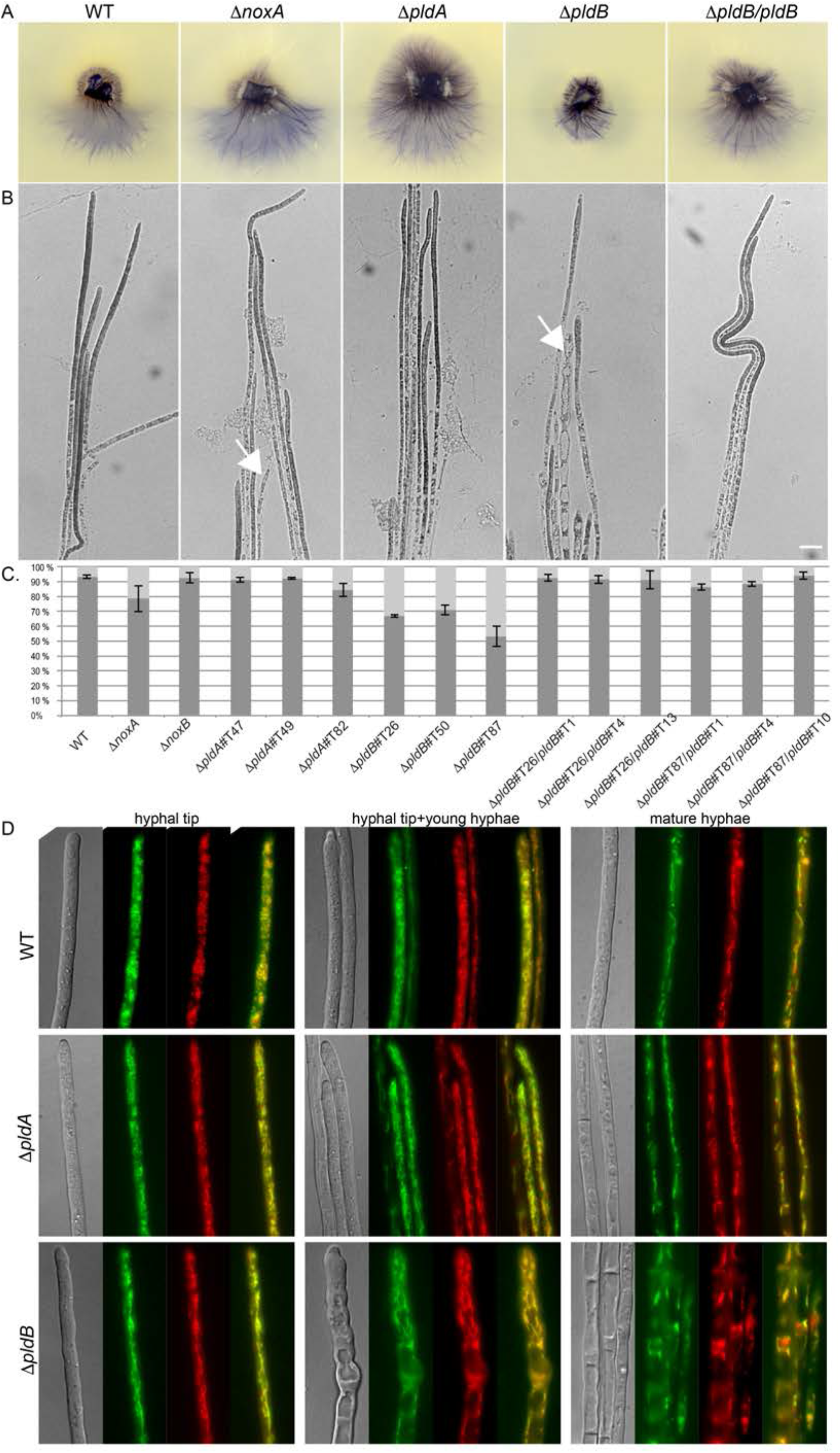
Analysis of superoxide production in *E. festucae* wild-type, Δ*pldA* and Δ*pldB* strains. (A) Whole-colony image of seven-day old colonies stained with 150 µl of 6.1 mM Nitroblue-Tetrazolium (NBT) solution (Sigma) for five hours. Three independent Δ*pldA* and Δ*pldB* strains were analysed, and here, representative images of WT, Δ*noxA, ΔpldA*#T47, Δ*pldB*#T87, and Δ*pldB*#T87/*pldB*#T4 are shown as representative of all strains analysed. The staining was performed twice. (B) Representative microscopic images of hyphal tips in (A); White arrows: aberrantly stained hyphae. Bar=10 µm. (C) Analysis of the occurrence of aberrant hyphal tip staining; 3x50 hypha of two different colonies of the same strain were analysed and the averages plotted (100% = 50 hyphal tips). Statistical testing revealed that the Δ*noxA* and Δ*pldB* strains were significantly different from WT (Δ*noxA*: P=2.28E-2; Δ*pldB* #T26: P=7.5E-7; Δ*pldB*#T50: P=1.10E-5; Δ*pldB*#T87: P<1E-8). Error bars represent the standard deviation. (D) Localization of superoxide production by staining with 0.16 mM of ROS-ID® Superoxide probe and co-localization with mitochondrial staining reagent-Green. The Δ*pldA* strains T47 and T82 and the Δ*pldB* strains T26 and T87 were analysed and images of Δ*pldA*#T47 and Δ*pldB*#T87 shown here are representative of all strains analysed. Bar=10 µm.

Hyphae were also stained with the ROS-ID^®^ superoxide probe. In WT tips the probe localised to puncta and in mature hyphae the dye accumulated in long filaments containing more intense puncta. Given these structures colocalised with the mitochondrial Reagent-Green marker, the superoxide detected appears to be generated as a by-product within mitochondria. Similar observations were made with Δ*pldA* strains. For the Δ*pldB* strains the ROS probe localised to long filamentous structures in tips, which, with the exception of mature hyphae, were not seen in WT or the Δ*pldA* mutant. These ROS positive structures also colocalised with the mitochondrial Reagent-Green marker demonstrating they are also mitochondria. It is likely that these unique distributions reflect distorted mitochondria and result from excessive vacuole formation characteristic of the Δ*pldB* strains (Fig. 11).

### The deletion of pldB may influence endocytic recycling

As mammalian PLD1 and PLD2 as well as PA have roles in both endo- and exocytosis, the vesicle trafficking phenotypes of the Δ*pldA* and Δ*pldB* mutants were examined by introducing GFP fusions of a suite of proteins known to localize to specific structures/stages of secretion and endosomal recycling, including: late Golgi cisternae (Vps52), the Spitzenköerper (Rab11), the exocyst (Sec3, Exo70), the endocytic collar (fimbrin) and early endosomes (Rab5) (Kilaru *et al*., 2015, Guo *et al*., 2015, Sánchez-León *et al*., 2015, Upadhyay & Shaw, 2008).

In WT strains, Vps52 localized to small, mobile puncta. While the apical region was mainly free of these, small accumulations or aggregates at the very apex of the tip were occasionally observed. Rab11 assumed a far more discreet distribution and was observed exclusively as a spot at the hyphal tip, presumably marking the Spitzenköerper. Although cytoplasmic signals could make clear identification difficult in some instances, the exocyst component Exo70 also collected at the hyphal tip but in a broader crescent shape than Rab11. Sec3 was observed as a crescent that emanated from a spot-like concentration also at the hyphal tip. These distributions differed markedly from the endocytic collar protein Fimbrin, which appeared as puncta that associated at the membrane and to a much lesser extent the cytoplasm, proximal to, but not at, the hyphal apex. An additional endocytic protein, Rab5, was also absent from the tip. This protein appeared as different sized accumulations which were motile (Fig. 12). The localization of Vps52, Rab11, Exo70, Sec3 and Rab5 GFP fusion proteins in Δ*pldA* and Δ*pldB* strains was indistinguishable from WT (Fig. 12). However, the characteristic sub-apical collar localization of fimbrin in WT was frequently absent in the Δ*pldB* mutant with the protein instead concentrating along the apical tip as a robust and broad crescent (Figs. 12 & S13).

**Fig. 12.**
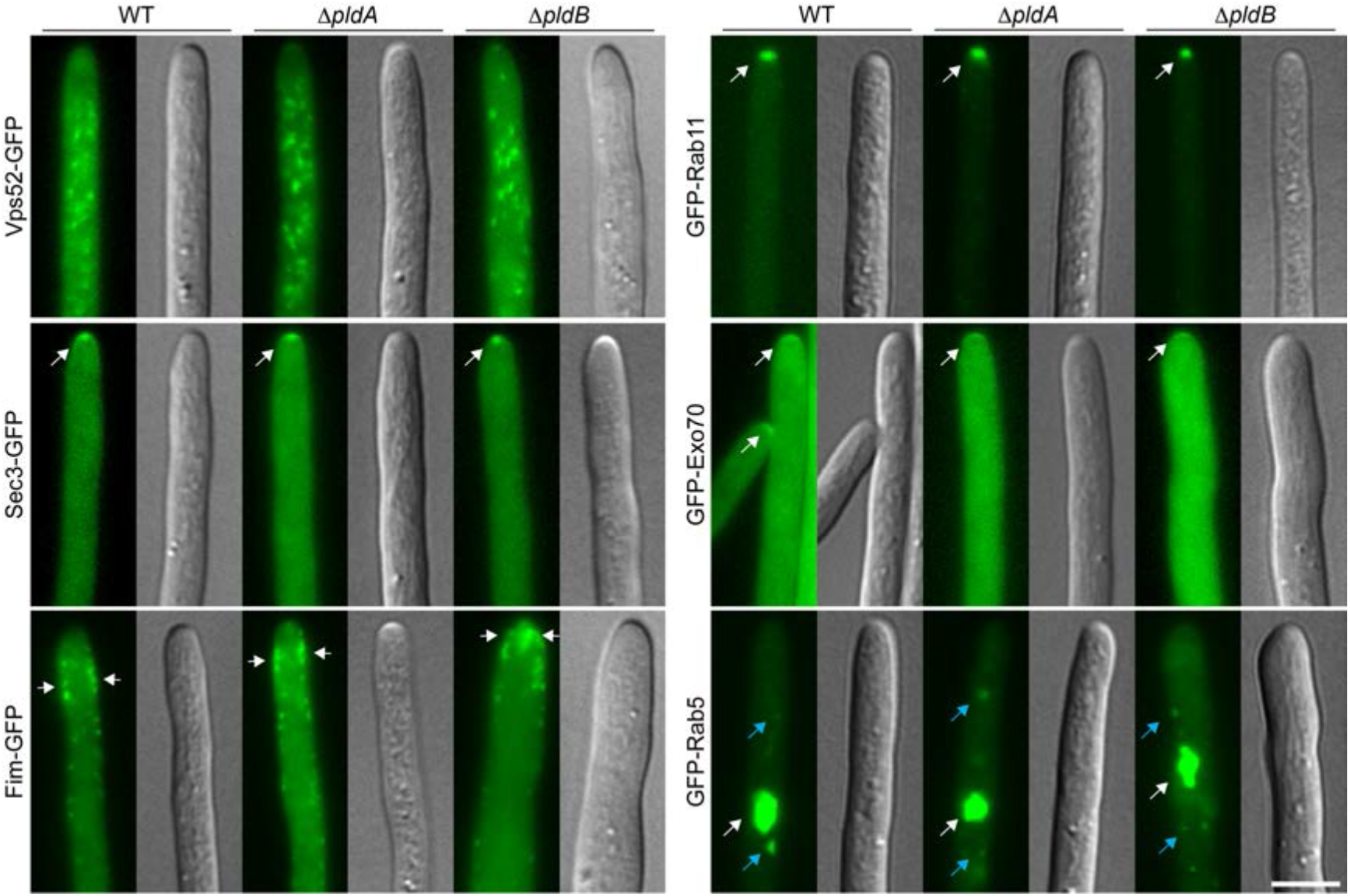
Localization of exo- and endocytosis markers in *E. festucae* wild-type, Δ*pldA* and Δ*pldB* strains. Constructs encoding translational fusion constructs between the protein of interest and GFP were transformed into wild-type (WT) and two different Δ*pldA* (#T47 and #T82) and Δ*pldB* (#T26 and #T87) strains, and multiple transformants analysed. Strains were grown for 5-7 days on H_2_O agar before analysis. Localization of (A) Vps52-GFP (EfM3.005680, pKG55) in WT (#T15), Δ*pldA*#T82 (#T6) and Δ*pldB*#T26 (#T7). (B) GFP-Rab11 (EfM3.017510, pBH71) in WT (#T17), Δ*pldA*#T82 (#T2) and Δ*pldB*#T87 (#T2). (C) Sec3-GFP (EfM3.039460, pBH69) in WT (#T9), Δ*pldA*#T82 (#T1) and Δ*pldB*#T26 (#T6). (D) GFP-Exo70 (EfM3.022070, pBH67) in WT (#T15), Δ*pldA*#T82 (#T6) and Δ*pldB*#T26 (#T6). (E) Fimbrin-GFP (EfM3.072070, pBH66) in WT (#T2), Δ*pldA*#T82 (#T3) and Δ*pldB*#T26 (#T1). (F) GFP-Rab5 (EfM3.068910, pBH68) in WT (#T1), Δ*pldA*#T82 (#T15) and Δ*pldB*#T26 (#T2). Blue arrow: Localisation of GFP-Rab5 to small moving vesicles. White arrows: Localisation of the respective fusion proteins. Bar=10 µm.

## Discussion

Membrane-bound conversion of phosphatidyl choline (PC) to phosphatidic acid (PA) has an important role for morphogenesis, growth and development in animals, plants and yeast, but very little is known about the importance of this process in filamentous fungi (Selvy *et al*., 2011). Here we show that phospholipase D (PLD)-dependent synthesis of PA is required for cell fusion in both *E. festucae* and *N. crassa*, highlighting the importance of lipid signaling for the fusion process. Cell fusion in fungi is crucial for hyphal network formation and multi-cellular development enabling nutrient transfer and signaling. This lipid conversion is also vital for development of a mutualistic symbiotic association between *E. festucae* and *L. perenne*.

The genome of *E. festucae* encodes two putative PC-hydrolysing PLDs, PldA and PldB, which differ significantly in their domain structure. PldB has PH and PX domains, which are also present in yeast, mammalian and plant Pld*ς* homologs (Selvy *et al*., 2011). These domains bind specific phosphoinositides that are required for localization of the protein (Selvy *et al*., 2011, Stahelin *et al*., 2004, Sciorra *et al*., 2002, Hodgkin *et al*., 2000). PldB also contains a PtdIns[4,5]P_2-_binding polybasic motif domain that is present in many PLDs of higher eukaryotes (Selvy *et al*., 2011). This motif is required for activation of the protein, but not for its localization (Sciorra *et al*., 1999, Sciorra *et al*., 2002). All of these domains are absent in PldA, although this isoform has retained the phosphatidyltransferase activity domains. Consistent with this domain structure, PldB localises to lipid-rich structures, in septa and the PM, while PldA localized exclusively to the cytosol of the hyphae. *S. cerevisiae* Spo14 localizes to the cytoplasm, to endosomes, the nucleus, and is recruited to the prospore membrane during sporulation (Rudge *et al*., 1998, Li *et al*., 2000, Garcia-Lopez *et al*., 2011). Recruitment of mammalian PLD1 and PLD2 to perinuclear membranes, the Golgi apparatus, endosomes, the PM, and membrane ruffles was found to be highly dependent on the activation status and developmental stage (Colley *et al*., 1997, Du *et al*., 2003, Hughes & Parker, 2001, Selvy *et al*., 2011). Thus, PldA and PldB may also assume other distributions during different stages of development.

Deletion of *pldB* resulted in reduced colony growth and increased aerial hyphae, phenotypes similar to those observed in *F. graminearum* Δ*pld1* strains (Ding *et al*., 2017). Microscopic analysis of the Δ*pldB* strains in culture showed that hyphae were often variable in diameter, abnormally shaped, frequently swollen and had an uncommonly high degree of vacuolation. Intra-hyphal hyphae were also occasionally detected. In contrast, growth of Δ*pldA*, *pldA* OE and *pldB* OE were indistinguishable from WT as found for mutants of the *pldA* homologs in *A. nidulans* (*AnpldA*) and *F. graminearum* (*Fgpld2*) (Ding *et al*., 2017, Hong *et al*., 2003). These observations highlight the importance of PldB, but not PldA, for normal polarized growth. While genetic complementation was able to restore the WT-like phenotype, the exogenous addition of PA did not, indicating that PldB itself is required for normal hyphal growth. Interestingly, similar phenotypes were observed for the *E. festucae* Δ*racA* mutant (Tanaka *et al*., 2008, Kayano *et al*., 2018). RacA is a small GTPase, which contributes to actin polymerization and Nox complex activation and has been found to interact with PLDs in mammalian cells (Jang *et al*., 2012, Hess *et al*., 1997, Bishop & Hall, 2000). Indeed, PLD2 has been found to act as a guanine exchange factor (GEF) for mammalian Rac2 (Mahankali *et al*., 2011).

Deletion of either *pldB* in *E. festucae* or *pla-7* in *N. crassa* abolished hyphal fusion in axenic culture, demonstrating that PldB/PLA-7 is a newly discovered component of the cell-to-cell communication and cell fusion signaling network between cells of Ascomycete filamentous fungi (Read *et al*., 2010, Fischer & Glass, 2019). To date, approximately 70 mutants with a fusion defect have been identified, and among these are proteins required for the cell wall integrity and pheromone response MAP kinase pathways, the STRIPAK complex, ROS signaling, as well as phosphatases and transcription factors (Fischer & Glass, 2019). The first step in the cell fusion process is chemotropic growth of germlings or hyphae towards each other, followed by degradation of the cell wall and fusion of membranes. Because the PA molecule has a cone shape, it can influence membrane curvature and could therefore be mechanistically required for membrane fusion during hyphal cell fusion (Selvy *et al*., 2011, Fischer & Glass, 2019). However, in Δ*pla-7* mutants, chemotropic growth was not observed, indicating that instead of a requirement of PA for membrane fusion, PA may actually be required for signaling during chemotropic growth. Interestingly, defects in ergosterol biosynthesis also result in cell-to-cell communication and cell fusion defects (Weichert *et al*., 2016). Nevertheless, exogenous addition of PA was able to restore hyphal fusion to the *E. festucae ΔpldB* strains. Whether this chemical complementation can be attributed to the restoration of signaling pathways required for chemotropic growth or changes in the ability of the membranes to fuse, remains to be determined. In mammals, PA interacts with, and recruits, proteins such as Rho and Arf GTPases, which may influence vesicle transport and actin cytoskeleton organization (Jang *et al*., 2012). Indeed, formation of an actin cytoskeleton is an important component of the hyphal fusion process as shown by treatment of *N. crassa* with the actin-polymerization inhibitor latrunculin B, which abolished hyphal fusion (Roca *et al*., 2010). The PLD proteins also bind actin and interact with a multitude of proteins in mammals (Lee *et al*., 2001, Kusner *et al*., 2002), including the Rho family GTPases, CDC42 and RAC1, the homologs of which are required for cell polarity and hyphal fusion in *N. crassa*, respectively (Lichius & Lord, 2014, Araujo-Palomares *et al*., 2011). In *S. cerevisiae* Cdc42 activates Spo14 (PLD1) and PA activates the p21-activated kinase Ste20 during polarized growth in response to pheromone (Harkins *et al*., 2008). While our yeast two-hybrid analysis did not reveal interactions between *E. festucae* PldB and Rac1 or Cdc42, these results do not exclude the possibility as detecting interactors with the yeast homolog, Spo14 (PLD1), is inherently difficult (Riedel *et al*., 2005). Whether the lack of fusion observed here is due to a defect in the ability of PLD or PA to interact with the actin cytoskeleton or on other signaling pathways required for fusion such as the CWI MAP kinase pathway or ROS signaling, remains to be elucidated.

Deletion of the *pldB* homolog in *N. crassa* resulted in a female-sterile phenotype, characterised by the failure to produce pre-fruiting bodies. Only crosses where the female is WT resulted in the formation of mature perithecia and ascospores, a phenotype commonly observed for other fusion-negative mutants (Fischer and Glass, 2019). This developmental stage could not be analysed in *E. festucae* as it only undergoes sexual reproduction in association with the host plant (Bultman & Leuchtmann, 2009). A defect in perithecia formation was also observed for the *F. graminearum Δpld1* strain (Ding *et al*., 2017).

In *N. crassa*, asci resulting from a cross between a Δ*pla-7* male and a WT female, were reduced in number and malformed, a phenotype not commonly observed for fusion mutants. *N. crassa* possesses a mechanism called meiotic silencing by unpaired DNA (MSUD), in which the RNA generated from unpaired DNA at meiosis is silenced (Shiu *et al*., 2001, Hammond, 2017). As *pla-7* is absent in the genome of the male partner in these crosses, MSUD likely results in the complete silencing of *pla7* expression during meiosis. Therefore, these results suggest that PLA-7 has a role in ascus and/or ascospore development, as has been show for Spo14, the homolog from *S. cerevisiae* (Rudge *et al*., 1998). While meiosis itself is not affected, deletion of *SPO14* results in the inability of vesicles to fuse to the prospore membrane due to the loss of activation and recruitment of the PA-binding *t*-SNARE Spo20 to the prospore membrane (Connolly & Engebrecht, 2006, Nakanishi *et al*., 2006). However, while Spo14 is absolutely essential for spore formation, PLA-7 is not, as some viable ascospores were obtained in these crosses, although at a much lower frequency of segregation (17%) than expected (50%). Whether this outcome correlates with the silencing of Pla7 during meiosis or is due to other functions of PLA-7 or PA remains to be determined.

PLA-7 also appears to have a role in the asexual life cycle of *N. crassa*, as conidiospore germination is delayed in the Δ*pla-7* mutant, a phenotype also observed for the *F. graminearum Δpld1* mutant (Ding *et al*., 2017). As *E. festucae* strain Fl1 conidiates very poorly it was difficult to assess this phenotype in this species (Becker *et al*., 2015). These results highlight the importance of PLA-7 for both the asexual and sexual lifecycles of *N. crassa*.

The severe host interaction phenotype observed for the Δ*pldB* mutant was very similar to that reported previously for several other *E. festucae* symbiotic mutants (Tanaka *et al*., 2006, Tanaka *et al*., 2013, Eaton *et al*., 2010, Becker *et al*., 2015, Takemoto *et al*., 2006, Green *et al*., 2017). Instead of wild-type like restricted hyphal growth, the Δ*pldB* mutant had a proliferative pattern of growth *in planta*. There was an increase in the number of hyphae in the host intercellular spaces of the leaves and colonization of the host vascular bundles, phenotypes not seen in WT associations. A transcriptome analysis of *E. festucae* symbiosis-defective mutant strains in association with *L. perenne* revealed an upregulation of genes in primary metabolism, host cell wall degradation, and peptide and sugar transporters; changes indicative of a starvation response (Eaton *et al*., 2015, Chujo *et al*., 2019). Indeed, all fusion-negative *E. festucae* strains obtained to date have a stunted host phenotype (Scott *et al*., 2018). However, given *E. festucae* mutants in cAMP/PKA signaling (Voisey *et al*., 2016), apoplastic iron homestasis (Johnson *et al*., 2013), and chromatin regulation (Chujo *et al*., 2019), exhibit a proliferative growth phenotype *in planta* but are not defective in cell fusion, an alternative more general hypothesis is needed to explain these results. One possibility is alkalinisation of the apoplast (Scott *et al*., 2018); a change that activates signaling through the PacC-mediated pH responsive and MAP kinase pathways, triggering invasive growth (Fernandes *et al*., 2017).

In addition to the hyphal fusion defect, the Δ*pldB* mutant was unable to form expressoria, appressorium-like structures that allow endophytic hyphae below the cuticle to swell up and penetrate this layer to emerge and grow on the surface of the leaf. Instead, the hyphae proliferate below the surface of the cuticle, a phenotype also observed for *E. festucae nox* complex, and some other symbiotic mutants that are defective in a conserved signaling network for cell fusion and multi-cellular development (Becker *et al*., 2016, Green *et al*., 2017, Green *et al*., 2019, Scott *et al*., 2018). Development of a functional appressorium in *M. oryzae* requires both Nox1 and Nox2 (Egan *et al*., 2007). Nox2 is necessary for septin-mediated actin ring reorientation at the septal pore, and Nox1 for maintenance of the F-actin network during penetration and elongation (Ryder *et al*., 2013). Given the similarity in expressorium phenotypes of the *E. festucae* Δ*pldB* and *nox* complex mutant strains, PldB or the PA product may be involved in Nox complex assembly and/or activation leading to a change in the actin cytoskeleton network. Indeed, PA has been shown to be involved in the recruitment of the cytosolic components of the Nox complex in plant and mammalian systems (Regier *et al*., 2000, Kanai *et al*., 2001, Karathanassis *et al*., 2002, Ago *et al*., 2003, Chae *et al*., 2008). However, deletion of either *pldA* or *pldB* in *E. festucae* did not alter the cellular localization of the Nox complex components, NoxA, NoxR, BemA, Cdc24 or RacA.

PA has also been shown to stimulate mammalian NADPH oxidase activity in a cell-free system through direct interaction with gp91^phox^, and in plant cells, by direct interaction with the NADPH oxidase (Palicz *et al*., 2001, Qualliotine-Mann *et al*., 1993, Taylor *et al*., 2004, Taylor *et al*., 2012, Zhang *et al*., 2009). While the *E. festucae* Δ*pldB* strains showed reduced NBT tip staining, indicative of reduced production of superoxide, this synthesis was confined to the mitochondria. Given the more filamentous structure of these structures in hyphal tips of the Δ*pldB* mutant strains compared to WT, it appears that PldB has a role in organelle structural organisation rather than superoxide production itself. Deletion of components of the Nox complex in *E. festucae*, *A. nidulans*, *F. graminearum* and *Sclerotinia sclerotium* also results in reduced NBT staining (Kayano *et al*., 2018, Semighini & Harris, 2008, Tanaka *et al*., 2006, Wang *et al*., 2014, Kim *et al*., 2011). Whether these reported changes in NBT staining are the result of changes in the structure of the mitochondria remains to be determined. Therefore, while there are many phenotypic similarities between Δ*pldB* and Nox component deletion mutants, a direct connection between PA and the Nox complex could not be demonstrated.

To analyse the effect of PLD deletion on PA concentration and distribution, a PA molecular biosensor was generated using the defined PA-binding domain of Spo20 (Potocký *et al*., 2014, Nakanishi *et al*., 2004). Microscopic analysis revealed that the PA biosensor localised to the PM in *E. festucae* WT and Δ*pldA* strains, as has been described in *S. cerevisiae* and in tobacco root hairs (Potocký *et al*., 2014, Nakanishi *et al*., 2004). In Δ*pldB* strains, localization of the PA biosensor was reduced at the PM in hyphal tips and young hyphae and preferentially localised to the cytoplasm and nuclei. The preferential localization to the nuclei is similar to what is observed when cells are treated with the PLD inhibitor *n*-butanol, resulting in a decrease in the PM/nucleus fluorescence intensity ratio (Potocký *et al*., 2014, Zeniou-Meyer *et al*., 2007). Therefore, an increased nuclear as well as cytoplasmic localization likely reflects reduced PLD activity. As the PA biosensor was never detected in nuclei of either the WT or Δ*pldA* strains, PldA is most likely to be the isozyme responsible for PA synthesis in *E. festucae*.

Both PLD and PA also have a role in vesicle trafficking, as demonstrated by their roles in endo- and exocytosis in yeast and mammalian cells (Li *et al*., 2000, Selvy *et al*., 2011, Donaldson, 2009). The detection of CFW-stained puncta in the Δ*pldB* strain is indicative of defects in vesicular trafficking (Higuchi *et al*., 2009, Upadhyay & Shaw, 2008). Using a range of GFP-labelled proteins associated with various steps in vesicle transport, we examined whether the localization of these proteins in Δ*pldB* was different to WT. However, apart from fimbrin, all the other GFP fusion proteins had the same localization as in WT. Fimbrin localised to actin patches at the endocytic collar, where it is known to crosslink with actin filaments. These filaments coat endocytic vesicles and provide mechanical force for membrane invagination (Riquelme *et al*., 2018, Aghamohammadzadeh & Ayscough, 2009). In the Δ*pldB* deletion strains, the Fimbrin-GFP fusion protein, while also detected at the endocytic collar, was found to collect at the hyphal tip more often than in WT hyphae. Fimbrin is known to oscillate between the hyphal tip and the endocytic collar and only concentrates at the endocytic collar in actively growing hyphae (Upadhyay & Shaw, 2008). While this increased tip localization of fimbrin in the Δ*pldB* strains could be a result of the slower growth rate of the Δ*pldB* strain it could also be indicative of a key role for PldB in endocytosis, as has been described in mammalian cells (Donaldson, 2009). Interestingly, deletion of fimbrin in *A. nidulans*, results in swollen hyphae, reduced spore germination and an accumulation of CFW in the cell, defects that are associated with loss of cell polarity (Upadhyay & Shaw, 2008). While fimbrin itself is unlikely to bind PA or interact with PLDs these observations suggest that PA or PLDs are involved with localization of the endocytic machinery and/or establishment of cell polarity.

In conclusion, we have demonstrated that PldB/PLA-7, one of two PLDs found in *E. festucae* and *N. crassa*, has an important role in hyphal morphogenesis and development. Crucially, PldB generated synthesis of PA is required for normal polarized hyphal growth, cell fusion and ascospore formation. Additionally, we have shown that PldB is required for expressoria formation and development of a mutualistic symbiotic interaction between *E. festucae* and *L. perenne*. Given the changes in lipid membrane structure and composition following the PLD dependent synthesis of PA, it will be of considerable future interest to identify which proteins are recruited to these lipid sites to initiate cell-to-cell communication and cell fusion.

## Experimental procedures

### Bioinformatic analyses

*E. festucae* sequences were obtained from the Kentucky Endophyte Database (Schardl *et al*., 2013). Protein sequences were annotated using InterproScan (Jones *et al*., 2014) and SMART (Letunic *et al*., 2015). Domains in the PLDs were annotated based on previous publications (Sciorra *et al*., 1999, Sciorra *et al*., 2002, Sung *et al*., 1997). BLASTp analyses were performed against fungal reference proteins at NCBI. Alignments were generated using MAFFT version 7 (Katoh *et al*., 2017) and shaded according to percentage identity in Jalview v1. The Ensembl genome database (Zerbino *et al*., 2018) was used to compare *E. festucae* PldA and PldB proteins to other PLD sequences by first obtaining a pre-computed protein alignment for the gene family with ID “EGGT00050000001203” and then adding the *Epichloë* sequences to this alignment using the “profile” method implemented in Muscle v3.8.31 (Edgar, 2004). A maximum likelihood phylogeny was estimated from the combined alignment using RAxML v 8.2.12 (Stamatakis, 2014) and the BLOSUM62 model of amino acid substitution. Box plots were generated using BoxPlotR (http://shiny.chemgrid.org/boxplotr/). All statistical tests are explained in detail in the Methods S1.

### Strains and growth conditions

*Escherichia coli* strains were grown in Lysogeny Broth (LB) (Miller, 1972) or on LB solidified with 1.5% (w/v) agar, supplemented with 100 µg/ml ampicillin or 50 µg/ml kanamycin at 37°C. *S. cerevisiae* strains were grown in YPDA liquid medium (Green, 2016) or on YPDA supplemented with 2% (w/v) agar overnight or for up to 3 days, respectively, at 30°C. Following transformation, strains were selected on SD synthetic -T/L dropout medium (Green, 2016), and incubated at 30°C until colonies appeared. For yeast two-hybrid assays strains were grown in liquid SD-T/L broth, then plated on SD-T/L/H and -T/L/H/A dropout media (Green, 2016), and were incubated for up to 5 days at 30°C. *E. festucae* strains were grown on Potato-Dextrose (PD) agar (2.4% PD, 1.5% agar, PDA) supplemented with 150 µg/ml hygromycin B and/or 200 µg/ml geneticin for up to 7 days at 22°C. For chemical complementation *E. festucae* strains were inoculated on 20 ml PDA and plates overlaid with 7 ml PDA containing 0.005, 0.02, 0.05 and 0.1 mg/ml 3-sn-phosphatidic acid sodium salt (Sigma) respectively, and grown for 7 days. *N. crassa* strains were cultivated on Vogel’s minimal medium supplemented with 2% sucrose as the carbon source and with 1.5% agar for solid media (Vogel, 1956). For propagation and spore production, strains were grown on agar slants and incubated at 30°C for 5-7 days. Crosses were performed on Westergaard’s medium at 25°C (Westergaard & Mitchell, 1947). For single spore isolations, conidia were plated on BDES medium agar plates (Brockman & de Serres, 1963) containing 200 µg/ml hygromycin. After an overnight incubation, individual colonies were isolated and transferred to agar slants.

All strain details are listed in Table S1.

### Plant growth and endophyte inoculation

*E. festucae* strains were artificially inoculated into *L. perenne* cv Samson seedlings and plants grown as described previously (Latch & Christensen, 1985, Becker *et al*., 2015).

### DNA isolation, PCR and sequencing

Genomic DNA was purified from freeze-dried *E. festucae* mycelia as described previously (Byrd *et al*., 1990). Plasmid DNA was extracted from *E. coli* strains using the High Pure plasmid isolation kit (Roche, Basel, Switzerland). PCR amplification of short DNA fragments (<3000 bp in length) was conducted using OneTaq^®^ DNA polymerase (New England Biolabs (NEB), Ipswich, USA) and amplification of DNA fragments for cloning purposes was performed with the Phusion^TM^ DNA polymerase (NEB) according to the manufacturer’s instructions. DNA fragments were separated by agarose-gel electrophoresis and purified with the Wizard SV Gel and PCR Clean-Up System (Promega, Madison, USA). Sequence authenticity was confirmed at the Massey Genome Centre. Sequence data was assembled (ClustalW) and analysed using MacVector v14.5.2.

### Preparation of constructs

All constructs were designed using MacVector v14.5.2 and synthesised using Gibson assembly (Gibson *et al*., 2009). Primer sequences are listed in Supporting Information Table S2 and descriptions of construct designs in Methods S2.

### Transformation of organisms

Chemically competent *E. coli* DH5*α* cells were transformed as described previously (Hanahan, 1983). *S. cerevisiae* AH109 cells were made competent using lithium acetate and transformed via heat-shock at 42°C for 20 min (Gietz & Woods, 2002). *E. festucae* protoplasts were generated as described elsewhere (Young *et al*., 2005), and transformed with 1.2-5 µg of each plasmid (Itoh *et al*., 1994). Protoplasts were allowed to recover overnight on regeneration (RG, PD with 0.8 M sucrose) medium supplemented with 1.5% agar before being overlaid with RG medium solidified with 0.8% agar and the appropriate antibiotic. For the deletion of *pldA* a linear 5’*pldA*-*hph*-3’*pldA* replacement cassette was amplified by PCR from pCE131 and transformed into wild-type (WT) protoplasts. For the deletion of *pldB*, the linear 5’*pldB*-*nptII*-3’*pldB* replacement cassette was amplified by PCR in two fragments from pCE121 with an overlapping region of approx. 600 bp (Rahnama *et al*., 2016). Protoplasts of the Δ*pldB* strain were allowed to recover for two days before being overlaid with 20-50 µg/ml hygromycin B.

### Protein extraction and western blot analysis

*E. festucae* strains were grown shaking in PD liquid medium for 3-4 days at 21°C. The washed and snap frozen mycelium was ground to a fine powder and resuspended in 0.2-1 mL of lysis buffer (50 mM Tris-HCL, 100 mM NaCl, 10 mM EDTA, 0.2 mM PMFS, 0.1 mM DTT, 1 µl/mL NP40, 1 µl/mL Protease inhibitor cocktail (Roche)). Cell debris was removed by centrifugation and 15-20 µl of the protein suspension was resolved on a 7 or 10% SDS-polyacrylamide gel. After electrophoresis proteins were transferred to a PVDF membrane. The membrane was incubated with a 1:3,000 dilution of primary rabbit anti-GFP antibody (ab290, Abcam, Cambridge, UK) or a 1:2,000 dilution of primary rabbit anti-mCherry antibody (ab167453, Abcam) overnight. This was followed by incubation with a 1:10,000 dilution of secondary goat anti-rabbit horseradish peroxidase (HRP) antibody (ab6721, Abcam) for one hour. Detection of HRP was performed using the Amersham ECL Western blotting detection reagent (GE Healthcare, Chicago, USA).

### RNA extraction and qRT-PCR

RNA was extracted from snap-frozen and ground mycelia using TRIzol (Invitrogen, Carlsbad, USA), and cDNA synthesis performed with the QuantiTect Reverse Transcriptase kit (Qiagen, Venlo, the Netherlands). RT-qPCRs were performed with SYBR Green (Invitrogen) on a LightCycler 480 System (Roche), as described by the manufacturer. Each sample was analysed with two technical replicates. *E. festucae* translation elongation factor 2 (*EF-2*, EfM3.021210) and 40S ribosomal protein S22 (*S22*, EfM3.016650) genes were used as references (Lukito *et al*., 2015). The gene expression level in the overexpression strains was calculated relative to the expression in WT (Lukito *et al*., 2015). Primer sequences used for this analysis (BH185-BH188) are listed in Table S2.

### Microscopy

For analysis of the hyphal morphology and localization of proteins, *E. festucae* strains were grown on a microscopic slide covered with 1.5% H_2_O agar for 5-7 days and analysed with an Olympus IX83 inverted fluorescence microscope. Cell walls were stained using 3 µl of a 3 mg/ml Calcofluor White (CFW) stock solution and nuclei with 3 µl of a 0.071 mM 4’,6-diamidino-2-phenylindole (DAPI) solution. For quantification of fusion events, colonies were incubated for 14 days before analysis. For chemical complementation, *E. festucae* strains were inoculated on 20 ml 1.5% H_2_O agar plates with a microscopic slide, overlaid with 7 ml 1.5% H_2_O agar containing 0.005 mg/ml, 0.02 mg/ml, 0.05 mg/ml, 0.1 mg/ml 3-sn-Phosphatidic acid sodium salt (Sigma) respectively, and grown for 7 days. Quantitative spore germination and fusion assays in *N. crassa* were conducted as described in (Fleißner *et al*., 2009). To observe hyphal fusion, strains were grown on Vogel’s minimal medium overnight at 30°C. Agar blocks of 1 cm^2^ were cut from the cultures and analysed by light microscopy.

For transmission electron microscopy (TEM), infected plant tissue was fixed and embedded as described by (Spiers & Hopcroft, 1993). Samples were analysed with a FEI Tecnai G2 Biotwin TEM. In addition, sections were stained with 0.05% toluidine blue in phosphate buffer and heat-fixed at approx. 100°C for 10 sec. These sections were examined with a Zeiss Axiophot Microscope with Differential Interference Contrast (DIC) Optics and Colour CCD camera.

To evaluate hyphal growth *in planta*, infected pseudostem samples were stained with aniline blue diammonium salt (Sigma-Aldrich, St. Louis, USA) and wheat germ agglutinin conjugated to AlexaFluor®488 (WGA-AF488, Molecular Probes/Invitrogen) (Becker *et al*., 2016, Becker *et al*., 2018), and examined with a Leica SP5DM6000B confocal microscope (488 nm argon and 561 nm DPSS laser, 40*×* oil immersion objective, NA = 1.3) (Leica Microsystems, Wetzlar, Germany).

### Superoxide detection

For nitroblue tetrazolium (NBT, Sigma-Aldrich) staining, colonies were grown and stained as reported previously (Tanaka *et al*., 2006). Colonies were photographed and imaged with a Zeiss Axiophot microscope as described. To quantify hyphal staining patterns, two colonies of each strain of interest were stained and three times approx. 50 hyphae evaluated for each.

Superoxide was detected using the ROS-ID^®^ Superoxide (ENZO LifeSciences, Farmingdale, USA) kit. The stock was diluted 1:30 (0.16 mM) in H_2_O and 3 µl was added to 7-day old colonies, followed by a 10 min incubation in the dark. For mitochondria identification a stock of Mitochondrial Staining Reagent-Green (ab176830, Cytopainter, Abcam) was diluted 1:500 in H_2_O and 3 µl added to the colonies. After 20 min incubation in the dark the ROS-ID ® Superoxide probe was added.

### Yeast two-hybrid

All plasmids used in yeast two-hybrid assays are listed in Table S1. Yeast strain AH109 was transformed as described above with each assay repeated twice as previously described (Green, 2016).

## Acknowledgements

This research was supported by grants from the Tertiary Education Commission to the Bio-Protection Research Centre, the Royal Society of New Zealand Marsden Fund (MAU1301) and by Massey University. BH was supported by a Massey University PhD studentship and BS by an Alexander von Humboldt Research Award. We thank Niki Murray and Pani Vijayan (Manawatu Microscopy and Imaging Centre) for support with microscopy, Arvina Ram and Alyesha Candy for technical assistance and Christopher Schardl for provision of *Epichloë* genome sequences.

## Author contributions

BH, CJE, CHM, AF and BS planned and designed the research. BH, CJE, KAG, DW, UB performed the experiments. BH, CJE, KAG, DW, MSS, CHM, AF and BS analyzed the results. CHM, AF and BS supervised the project. BH, CHM, AF and BS wrote the manuscript.

