## Supplemental Methods, Figures and Tables for "Phosphatidic acid produced by phospholipase D is required for hyphal cell-cell fusion and fungal-plant symbiosis"

The following Supporting Information is available for this article:

A

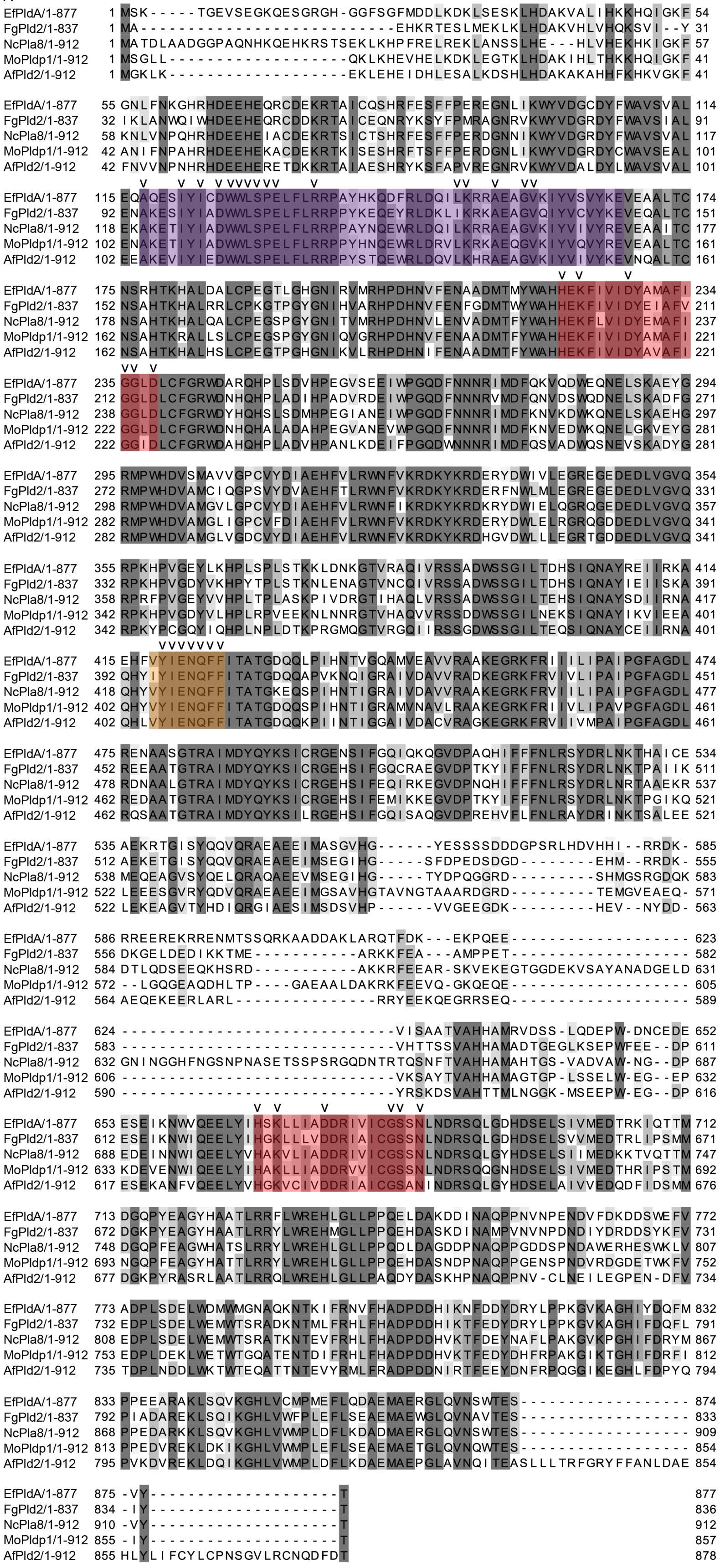
EfPldA/1-877 1 MSK-----TGEVSEGKQESGRGH-GGFSGFMDDLKDKLSESKLHDAKVALIHKKHQIGKF 54
FgPld2/1-837 1 MA---------------------------EHKRTESLMEKLKLHDAKVHLVHQKSVI--Y 31
NcPla8/1-912 1 MATDLAADGGPAQNHKQEHKRSTSEKLKHPFRELREKLANSSLHE---HLVHEKHKIGKF 57
MoPldp1/1-912 1 MSGLL-------------------QKLKHEVHELKDKLEGTKLHDAKIHLTHKKHQIGKF 41
AfPld2/1-912 1 MGKLK-------------------EKLEHEIDHLESALKDSHLHDAKAKAHHFKHKVGKF 41

EfPldA/1-877 55 GNLFNKGHRHDEEHEQRCDEKRTAICQSHRFESFFPEREGNLIKWYVDGCDYFWAVSVAL 114
FgPld2/1-837 32 IKLANWQIWHDEEHEQRCDDKRTAICEQNRYKSYFPMRAGNRVKWYVDGRDYFWAVSIAL 91
NcPla8/1-912 58 KNLVNPQHRHDEEHEIACDEKRTSICTSHRFESFFPERDGNNIKWYVDGRDYFWAVSVAL 117
MoPldp1/1-912 42 ANIFNPAHRHDEEHEKACDEKRTKISESHRFTSFFPERDGNLIKWYVDGRDYCWAVSEAL 101
AfPld2/1-912 42 FNVVNPNHRHDEEHERETDKKRTAIAESHRYKSFAPVREGNRVKWYVDALDYLWAVSVAL 101

EfPldA/1-877 115 EQAQESIYICDWWLSPELFLRRPAYHKQDFRLDQILKRRAEAGVKIYVSVYKEVEAALTC 174
FgPld2/1-837 92 ENAKESIYIADWWLSPELFLRRPPYKEQEYRLDKLIKRKAEAGVQIYVCVYKEVEQALTC 151
NcPla8/1-912 118 EKAKETIYIADWWLSPELFLRRPPAYNQEWRLDQILKRRAEAGVKIYVIVYREVEAAITC 177
MoPldp1/1-912 102 ENAKETIYIADWWLSPELFLRRPPYHNQEWRLDRVLKRKAEQGVKIYVQVYREVEAALTC 161
AfPld2/1-912 102 EEAKEVIYIEDWWLSPELFLRRPPYSTQEWRLDQVLKHRAEAGVKIYVIVYKEVNQALTC 161

EfPldA/1-877 175 NSRHTKHALDALCPEGTLGHGNIRVMRHPDHNVFENAADMTMYWAHHEKFIVIDYAMAFI 234
FgPld2/1-837 152 NSAHTKHALRRLCPKGTPGYGNIHVARHPDHNVFENFGDMTWYWAHHEKFIVIDYEIAFV 211
NcPla8/1-912 178 NSAHTKHALQALCPEGSPGYGNITVMRHPDHNVLENVADMTFYWAHHEKFLVIDYEMAFI 237
MoPldp1/1-912 162 NSAHTKRALLSLCPEGSPGYGNIQVGRHPDHNVFENAADMTFYWAHHEKFIVIDYAMAFI 221
AfPld2/1-912 162 NSAHTKHALHSLCPEGTPGHGNIKVLRHPDHNIFENAADMTFYWAHHEKFIVIDYAVAFI 221

EfPldA/1-877 235 GGLDLCFGRWDARQHPLSDVHPEGVSEEIWPGQDFNNNRIMDFQKVQDWEQNELSKAEYG 294
FgPld2/1-837 212 GGLDLCFGRWDNHQHPLADIHPADVRDEIWPGQDFNNNRVMDFQNVDSWQDNELSKADFG 271
NcPla8/1-912 238 GGLDLCFGRWDYHQHSLSDMHPEGIANEIWPGQDFNNNRIMDFKNVKDWKQNELSKAEHG 297
MoPldp1/1-912 222 GGLDLCFGRWDNHQHALADAHPEGVANEVWPGQDFNNNRIMDFKNVEDWKQNELGKVEYG 281
AfPld2/1-912 222 GGIDLCFGRWDAHQHPLADVHPANLKDEIFPGQDWNNNRIMDFQSVADWQSNEVSKADYG 281

EfPldA/1-877 295 RMPWHDVSMAVVGPCVYDIAEHFVLRWNFVKRDKYKRDERYDWIVLEGREGEDEDLVGVQ 354
FgPld2/1-837 272 RMPWHDVAMCIQGPSVYDVAEHFTLRWNFVKRDKYKRDERFNWLMLEGREGEDEDLVGVQ 331
NcPla8/1-912 298 RMPWHDVAMGVLGPCVYDIAEHFVLRWNFIKRDKYKRDKRYDWIELQGRQGEDEDLVGVQ 357
MoPldp1/1-912 282 RMPWHDVAMGLIGPCVFDIAEHFVLRWNFVKRDKYKRDERYDWLELRGRQGDDEDLVGVQ 341
AfPld2/1-912 282 RMPWHDVAMGLVGDCVYDIAEHFVLRWNFVKRDKYKRDHGVDWLLLEGRTGDDEDLVGVQ 341

EfPldA/1-877 355 RPKHPVGEYLKHPLSPLSTKKLDNKGTVRAQIVRSSADWSSGILTDHSIQNAYREIIRKA 414
FgPld2/1-837 332 RPKHPVGDYVKHPYTPLSTKNLENAGTVNCQIVRSSADWSSGILTDHSIQNAYIEIISKA 391
NcPla8/1-912 358 RPRFPVGEYVVHPLTPLASKPIVDRGTIHAQLVRSSADWSSGILTEHSIQNAYSDIIRNA 417
MoPldp1/1-912 342 RPKHPVGDYVLHPLRPVEEKNLNNRGTVHAQVVRSSDDWSSGILNEKSIQNAYIKVIEEA 401
AfPld2/1-912 342 RPKYPCGQYIQHPLNPLDTKPRGMQGTVRGQIIRSSGDWSSGILTEQSIQNAYCEIIRNA 401

EfPldA/1-877 415 EHFVYIENQFFITATGDQQLPIHNTVGQAMVEAVVRAAKEGRKFRIIILIPAIPGFAGDL 474
FgPld2/1-837 392 QHYIYIENQFFITATGDQQAPVKNQIGRAIVDAVVRADKEGRKFRVIVLIPAIPGFAGDL 451
NcPla8/1-912 418 QHYVYIENQFFITATGKEQSPIHNTIGRAIVDAVVRAAKEGRKFRVIILIPAIPGFAGDL 477
MoPldp1/1-912 402 QHYVYIENQFFITATGDQQSPIHNTIGRAMVNAVLRAAKEGRKFRIIVLIPAVPGFAGDL 461
AfPld2/1-912 402 QHLVYIENQFFITATGDQQKPIINTIGGAIVDACVRAGKEGRKFRVIIVMPAIPGFAGDL 461

EfPldA/1-877 475 RENAASGTRAIMDYQYKSICRGENSIFGQIQKQGVDPAQHIFFFNLRSYDRLNKTHAICE 534
FgPld2/1-837 452 REEAATGTRAIMDYQYKSICRGEHSIFGQCRAEGVDPTKYIFFFNLRSYDRLNKTPAIIK 511
NcPla8/1-912 478 RDNAALGTRAIMDYQYKSICRGEHSIFEQIRKEGVDPNQHIFFFNLRSYDRLNRTAAEKR 537
MoPldp1/1-912 462 REDAATGTRAIMDYQYKSICRGEHSIFEMIKKEGVDPTKYIFFFNLRSYDRLNKTPGIKQ 521
AfPld2/1-912 462 RQSAATGTRAIMDYQYKSILRGEHSIFGQISAQGVDPREHVFLFNLRAYDRINKTSALEE 521

EfPldA/1-877 535 AEKRTGISYQQVQRAEAEEIMASGVHG------YESSSSDDDGPSRLHDVHHI--RRDK- 585
FgPld2/1-837 512 AEKETGISYQQVQRAEAEEIMSEGIHG-----SFDPEDSDGD--------EHM--RRDK- 555
NcPla8/1-912 538 MEQEAGVSYQELQRAQAEEVMSEGIHG-----TYDPQGGRD---------SHMGSRGDQK 583
MoPldp1/1-912 522 LEEESGVRYQDVQRAEAEEIMGSAVHGTAVNGTAAARDGRD---------TEMGVEAEQ- 571
AfPld2/1-912 522 LEKEAGVTYHDIQRGIAESIMSDSVHP------VVGEEGDK---------HEV--NYDD- 563

EfPldA/1-877 586 RREEREKRRENMTSSQRKAADDAKLARQTFDK---EKPQEE------------------- 623
FgPld2/1-837 556 DKGELDEDIKKTME-----------ARKKFEA---AMPPET------------------- 582
NcPla8/1-912 584 DTLQDSEEQKHSRD-----------AKKRFEEAR-SKVEKEGTGGDEKVSAYANADGELD 631
MoPldp1/1-912 572 --LGQGEAQDHLTP----GAEAALDAKRKFEEVQ-GKQEQE------------------- 605
AfPld2/1-912 564 AEQEKEERLARL--------------RRYEEKQEGRRSEQ-------------------- 589

EfPldA/1-877 624 -----------------------------VISAATVAHHAMRVDSS-LQDEPW-DNCEDE 652
FgPld2/1-837 583 -----------------------------VHTTSSVAHHAMADTGEGLKSEPWFEE--DP 611
NcPla8/1-912 632 GNINGGHFNGSNPNASETSSPSRGQDNTRTQSNFTVAHHAMAHTGS-VADVAW-NG--DP 687
MoPldp1/1-912 606 -----------------------------VKSAYTVAHHAMAGTGP-LSSELW-EG--EP 632
AfPld2/1-912 590 -----------------------------YRSKDSVAHTTMLNGGK-MSEEPW-EG--DP 616

EfPldA/1-877 653 ESEIKNWVQEELYIHSKLLIADDRIVICGSSNLNDRSQLGDHDSELSIVMEDTRKIQTTM 712
FgPld2/1-837 612 ESEIKNWIQEELYIHGKLLLVDDRIAICGSSNLNDRSQLGDHDSELSVVMEDTRLIPSMM 671
NcPla8/1-912 688 EDEINNWIQEELYVHAKVLIADDRIVICGSSNLNDRSQLGYHDSELSIIMEDKKTVQTTM 747
MoPldp1/1-912 633 KDEVENWIQEELYIHAKLLIADDRVVICGSSNLNDRSQQGNHDSELSIVMEDTHRIPSTM 692
AfPld2/1-912 617 ESEKANFVQEELYVHGKVCIVDDRIAICGSANINDRSQLGYHDSELAIVVEDQDFIDSMM 676

EfPldA/1-877 713 DGQPYEAGYHAATLRRFLWREHLGLLPPQELDAKDDINAQPPNVNPENDVFDKDDSWEFV 772
FgPld2/1-837 672 DGKPYEAGYHAATLRRYLWREHMGLLPPQEHDASKDINAMPVNVNPDNDIYDRDDSYKFV 731
NcPla8/1-912 748 DGQPFEAGWHATSLRRYLWREHLGLLPPQDLDAGDDPNAQPPGDDSPNDAWERHESWKLV 807
MoPldp1/1-912 693 NGQPFEAGYHATTLRRYLWREHLGLLPPQEHDASNDPNAQPPGENSPNDVRDGDETWKFV 752
AfPld2/1-912 677 DGKPYRASRLAATLRRQLWREHLGLLPAQDYDASKHPNAQPPNV-CLNEILEGPEN-DFV 734

EfPldA/1-877 773 ADPLSDELWDMWMGNAQKNTKIFRNVFHADPDDHIKNFDDYDRYLPPKGVKAGHIYDQFM 832
FgPld2/1-837 732 EDPLSDELWEMWTSRADKNTMLFRHLFHADPDDHIKTFEDYDRYLPPKGVKAGHIFDQFL 791
NcPla8/1-912 808 EDPLSDELWEMWTSRATKNTEVFRHLFHADPDDHVKTFDEYNAFLPAKGVKPGHIFDRYM 867
MoPldp1/1-912 753 EDPLDEKLWETWTGQATENTDIFRHLFHADPDDHVKTFEDYDHFRPAKGIKTGHIFDRFI 812
AfPld2/1-912 735 TDPLNDDLWKTWTEQATTNTEVYRMLFRADPDDNIRTFEEYDNFRPQGGIKEGHLFDPYQ 794

EfPldA/1-877 833 PPEEARAKLSQVKGHLVCMPMEFLQDAEMAERGLQVNSWTES------------------ 874
FgPld2/1-837 792 PIADAREKLSQIKGHLVWFPLEFLSEAEMAEWGLQVNAVTES------------------ 833
NcPla8/1-912 868 PPEDARKKLSQVKGHLVWMPLDFLKDADMAERGLQVNSWTES------------------ 909
MoPldp1/1-912 813 PPEDVREKLDKIKGHLVWMPLEFLSEAEMAETGLQVNQWTES------------------ 854
AfPld2/1-912 795 PVKDVREKLDQIKGHLVWMPLDFLKDAEMAEPGLAVNQITEASLLLTRFGRYFFANLDAE 854

EfPldA/1-877 875 -VY--------------------T 877
FgPld2/1-837 834 -IY--------------------T 836
NcPla8/1-912 910 -VY--------------------T 912
MoPldp1/1-912 855 -IY--------------------T 857
AfPld2/1-912 855 HLYLIFCYLCPNSGVLRCNQDFDT 878

B

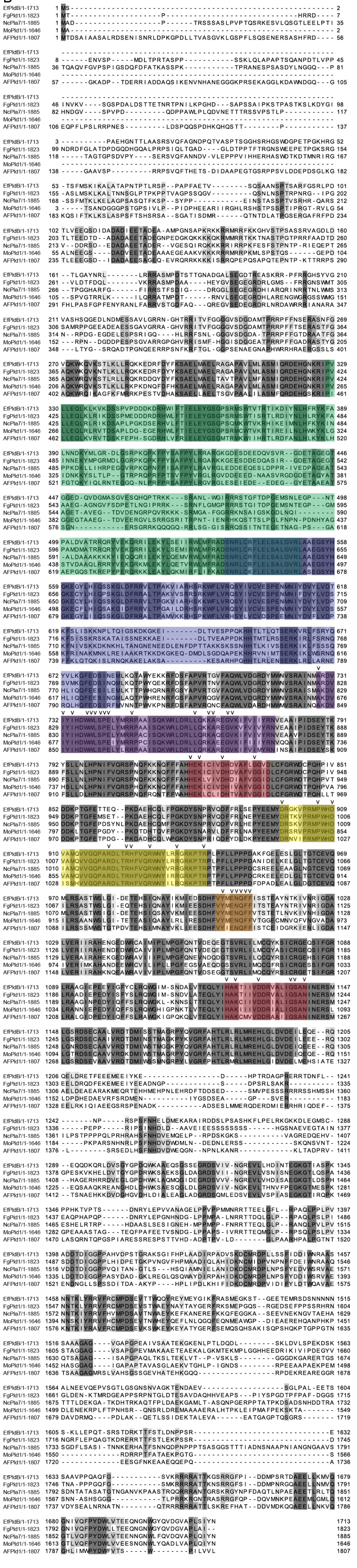

**Fig. S1.** Amino acid sequence alignment of *E. festucae* PldA and PldB with homologs from other Ascomycete fungi.

(A) PldA amino acid sequence alignment.

(B) PldB amino acid sequence alignment. Grey shading according to percentage identity, dark grey: high identity, light grey: low identity. Green box: PH (phox homology) domain; blue box: PX (pleckstrin homology) domain; purple box: motif I; red box: motif II and IV; orange box: motif III; yellow box: polybasic motif; *Ef*: *E. festucae* (EfPldB: EfM3.032570, EfPldA: EfM3.055250); *Fg*: *F. graminearum (*FgPld1: XP_011318823.1, FgPld2: XP_011317835.1); *Mo*: *M. oryzae* (MoPld1: XP_003717990.1, MoPldp1 XP_003712119.1); *Nc*: *N. crassa* (NcPLA-7: XP_957594.3*,* NcPLA-8: XP_001728077.1); *Af*: *A. fumigatus* (AfPld1: KEY78740.1, AfPldA; XP_748951.1). Amino acids marked with “v” are conserved according to Sung *et al.* (1997), Sciorra *et al*. (1999) and Sciorra *et al*. (2002).

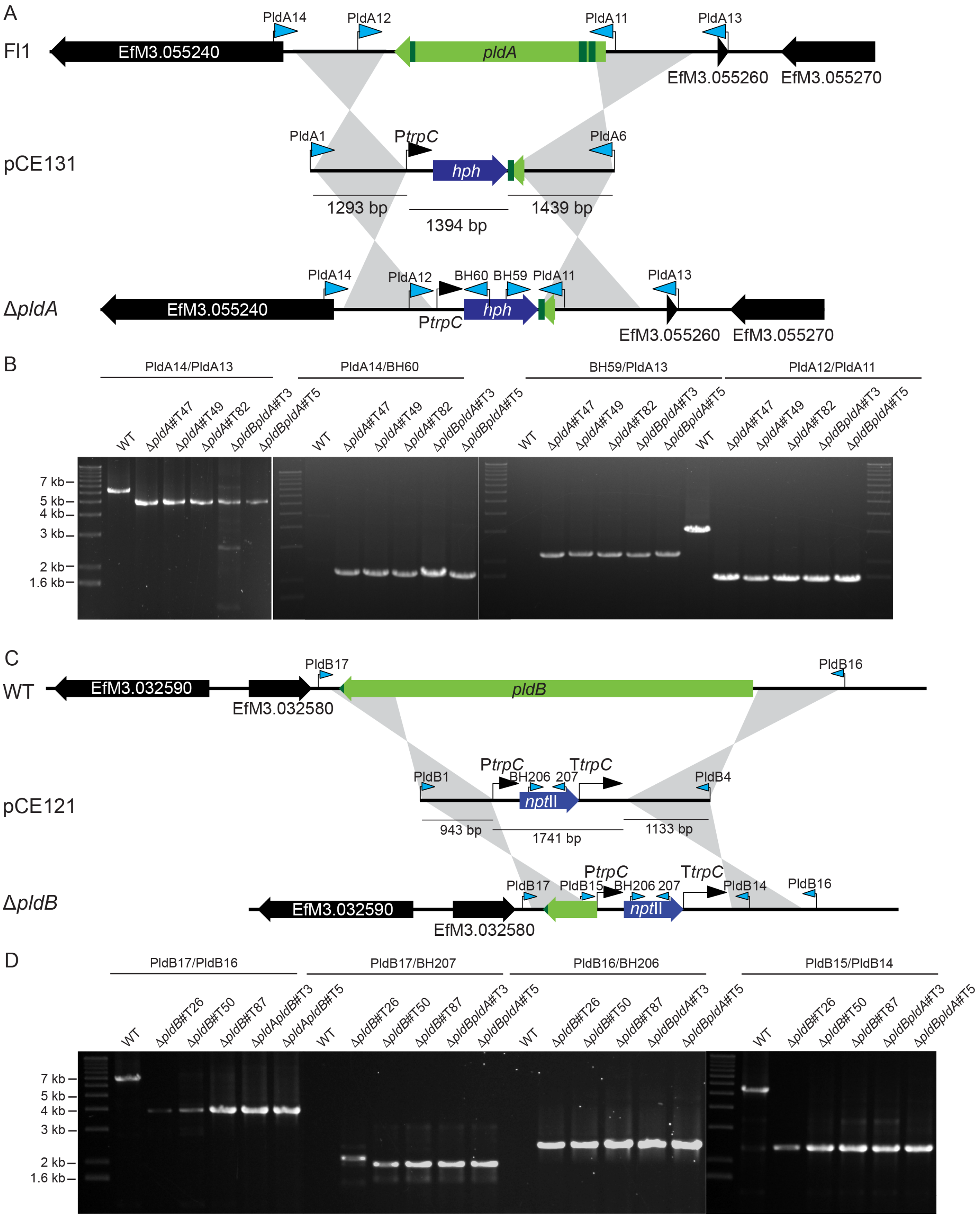

**Fig. S2.** Strategy for the deletion of *E. festucae pldA* and *pldB* and verification of deletion by PCR analysis.

(A) Physical map of the *pldA* wild-type (WT) genomic locus, linear insert of the *pldA* replacement construct, pCE131. Grey shading indicates regions of recombination. Numbers indicate the PCR primer pairs used for Gibson assembly (PldA1/PldA6) and deletion mutant screening (BH59/BH60/PldA11/PldA12/PldA13/PldB14).

(B) PCR screening of Δ*pldA* single and Δ*pldB*Δ*pldA* double deletion candidates. Screening with the PCR primer pair PldA14/PldA13 generated the expected bands of 6,261 bp in WT and 4,865 bp in the deletion mutants, the pair PldA14/BH60 generated expected band of 2,058 bp in deletion mutants and the pair BH59/PldA13 generated expected band of 2,114 bp in deletion mutants. As expected no band was observed for the last two PCRs in WT due to the lack of the *hph* gene. PldA11/PldA12, generated expected bands of 2,947 bp in WT, and 1555 bp in deletion mutants.

(C) Physical map of the *pldB* WT genomic locus, linear insert of the *pldB* replacement construct, pCE121. Grey shading indicates regions of recombination. Numbers indicate the PCR primer pairs used for Gibson assembly (PldB1/PldB4) and deletion mutant screening (BH206/BH207/PldB14/PldB15/PldB16/PldB17)

(D) PCR screening of Δ*pldB* single and Δ*pldB*Δ*pldA* double deletion candidates. Screening with the PCR primer pair PldB17/PldB16 generated expected band of 3,928 bp in deletion mutants and 7,024 bp in WT, the pair PldB17/BH207 generated expected band of 1,969 bp in deletion mutants and the pair BH206/PldB16 generated expected band of 2,465 bp in deletion mutants. As expected, no band was observed for the last two PCRs in WT due to the lack of the *nptII* gene. Screening with the PCR primer pair PldB15/PldB14 generated expected band of 2,394 bp in deletion mutants and 5,490 bp in WT.

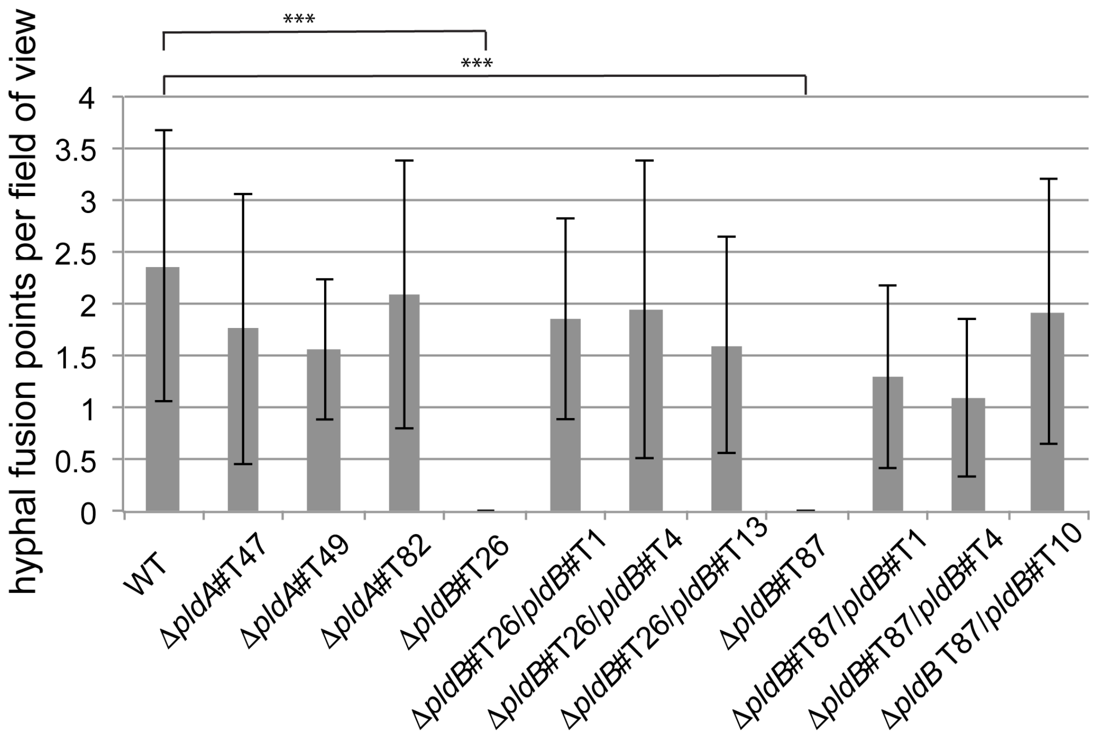

**Fig. S3.** Quantification of hyphal fusion events in *E. festucae* wild-type, Δ*pldA* and Δ*pldB* strains.

Wild-type (WT), deletion and complementation strains were grown on 3% $H_2O$ agar for 14 days and hyphal fusion events in 3 x 10 fields of view were counted per strain. The average of 30 counts, including the standard deviation, is plotted. Statistical tests used for this analysis are provided in Supporting Information Methods S1. Only Δ*pldB* strains #T26 and #T87 were significantly different from WT (P = 2.95e-8 and 3.4e-10, respectively).

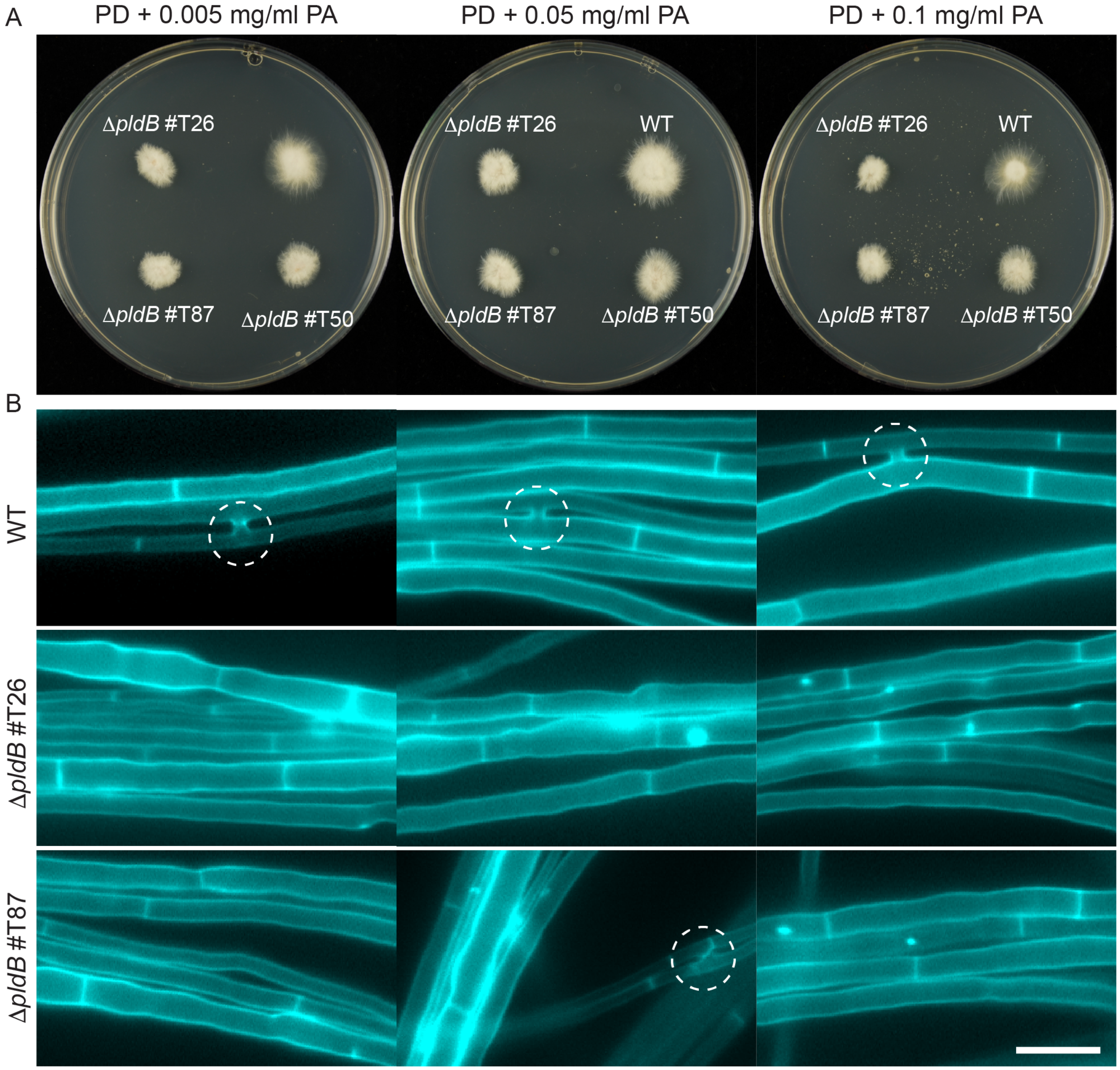

**Fig. S4.** Exogenous addition of phosphatidic acid restores hyphal cell to cell fusion in *E. festucae* Δ*pldB* strains.

(A) Wild-type (WT) and Δ*pldB* strains were grown on potato dextrose (PD) agar plates overlaid with 7 ml of PD agar containing 0.005 mg/ml, 0.05 mg/ml and 0.1 mg/ml phosphatidic acid (PA) and cultured for 7 days.

(B) WT and Δ*pldB* strains were grown on $H_2O$-agar plates overlaid with 7 ml of $H_2O$-agar containing 0.005 mg/ml, 0.05 mg/ml and 0.1 mg/ml PA, cultured for 7 days, and stained with

Calcofluor white before analysis to highlight sites of cell fusion. Dashed circles: hyphal fusion; Bar=10 μm.

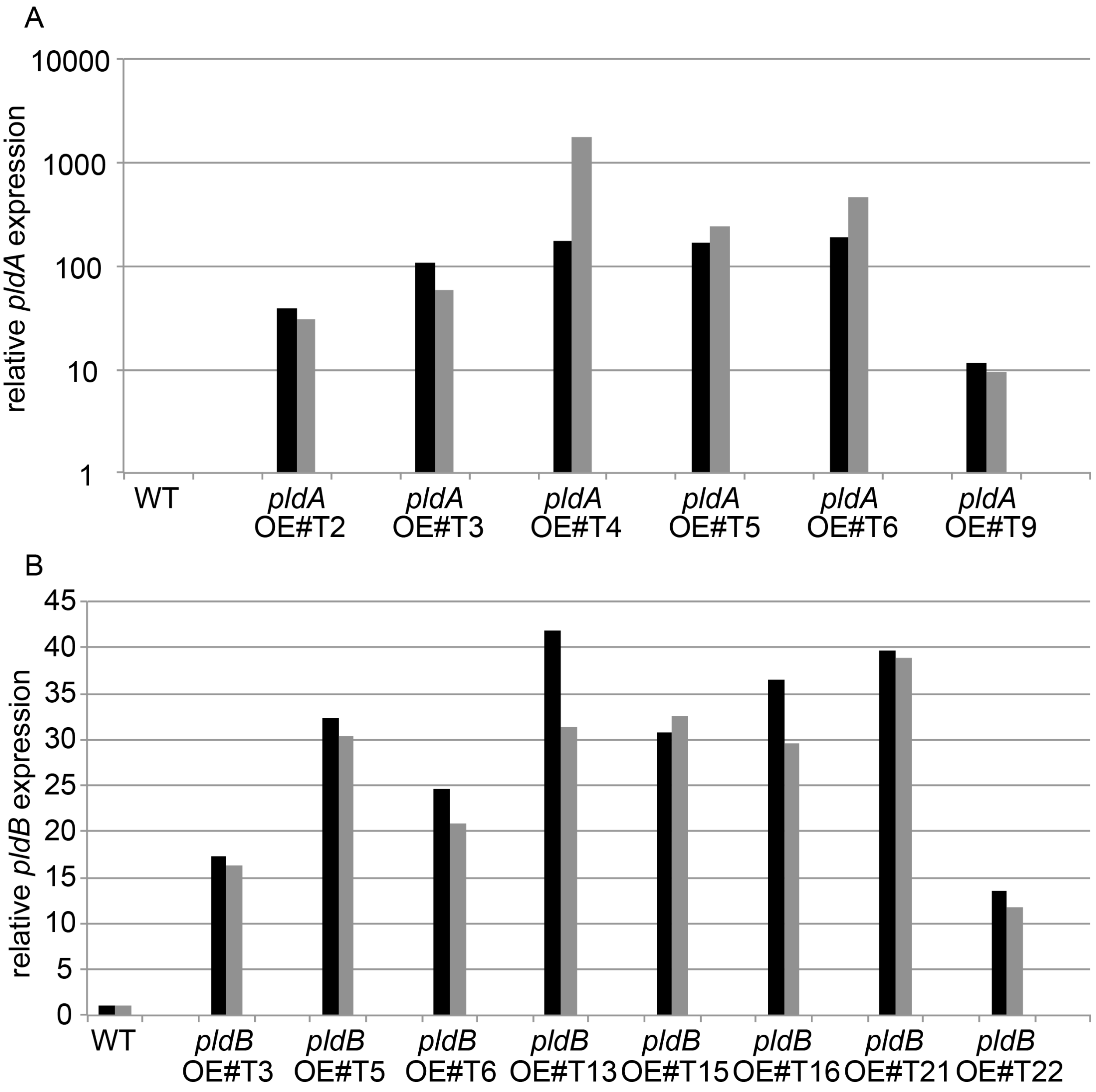

**Fig. S5.** RT-qPCR analysis of *E. festucae pldA* and *pldB* overexpression strains.

Relative expression of *pldA* (A) and *pldB* (B) was determined relative to wild-type (WT) gene expression. The analysis was performed with two technical replicates. Reference genes coding for elongation factor 2 (*EF-2*, black bar) and 40S ribosomal protein S22 (*S22*, grey bar) were used for normalization and the relative expression was calculated as described by Lukito *et al*.

(2015). Notice the different scales in (A) compared to (B). The *pldA* OE strains T3, T4 and T9 and *pldB* OE strains T6, T21 and T22 were chosen for further analysis. For the RT-qPCR the primer combinations BH185/BH186 and BH187/BH188 were used for *pldA* and *pldB*, respectively.

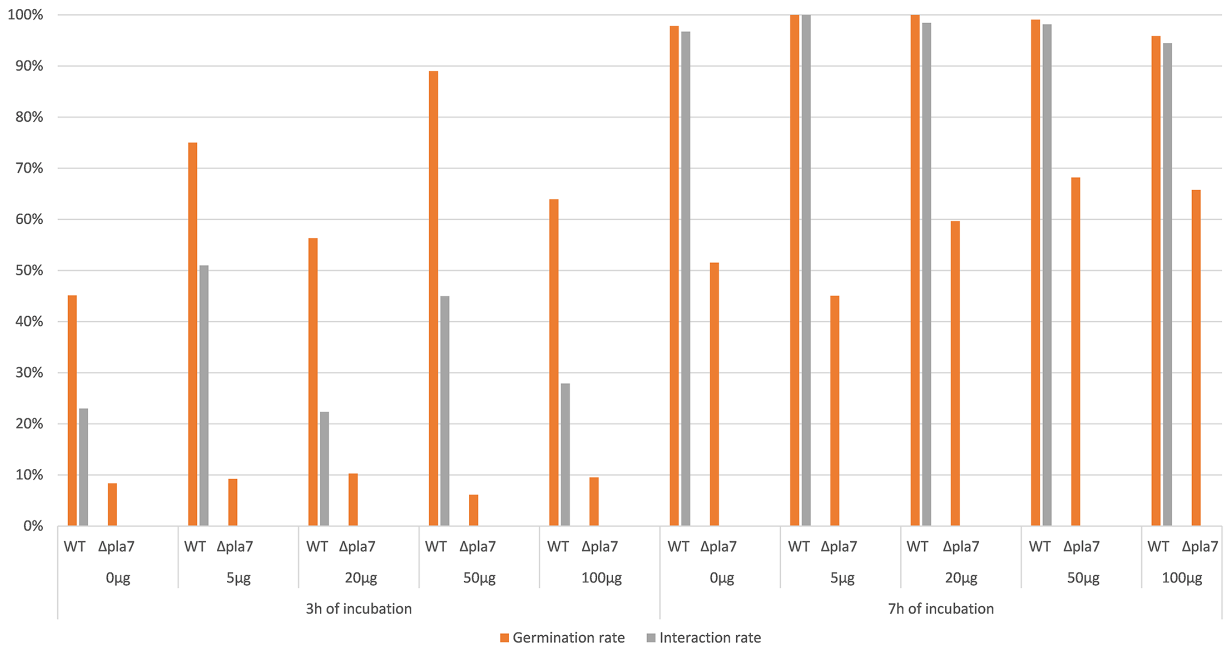

**Fig. S6. The fusion deficiency of the *N. crassa* Δ*pla-7* mutant is not complemented by the addition of PA to the growth medium.** Conidiospores of the wild type (WT) and the Δ*pla-7* mutant were spread on minimal medium containing different concentrations of PA (0, 5, 20, 50, 100 µg/ml). After 3 and 7 h of incubation the germination and interaction rates were determined (orange and grey bars, respectively). 100 germinated spores were analyzed per sample. Exogenous addition of phosphatidic acid does not restore germling fusion in *N. crassa* Δ*pla-7*.

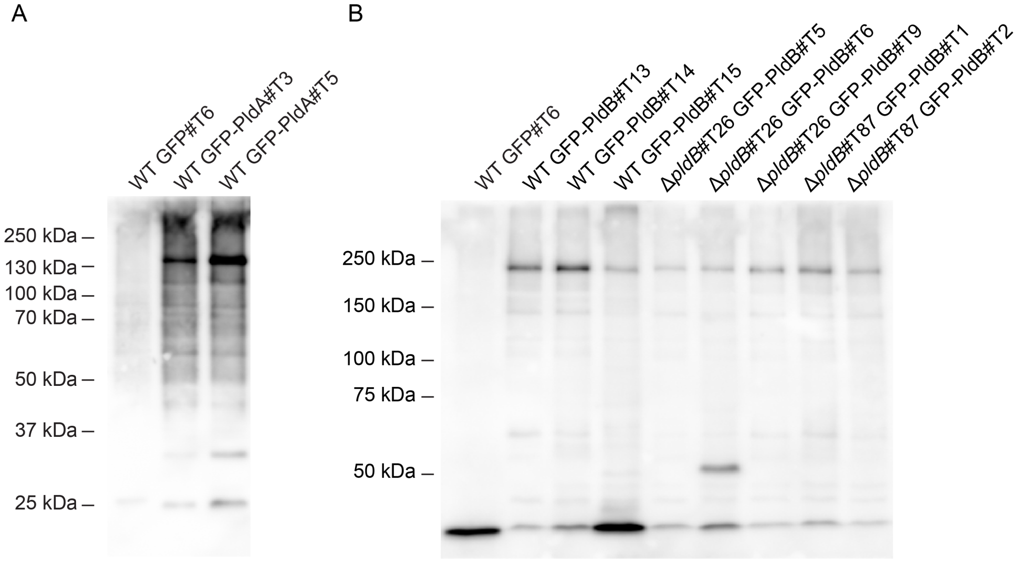

**Fig. S7.** Verification of the expression of GFP-PldA, GFP-PldB fusion proteins in *E. festucae* by western blot.

Strains expressing the fusion proteins were grown in potato-dextrose liquid media for 3 days, and total protein extracted. Proteins were resolved on a 10% (PldA fusions) or 7% (PldB fusions) acrylamide gel, transferred onto a PVDF membrane and probed with an α-GFP. Subsequently a secondary α-rabbit antibody conjugated to horseradish peroxidase was applied and the western blot was developed via a chemiluminescence reaction.

(A) Wild-type (WT) GFP (#T6) and WT GFP-PldA (pBH37; #T3 & #T5).

(B) WT GFP (#T6), WT GFP-PldB (pBH54; #T13, #T14 & #T15), Δ*pldB*#T26 GFP-PldB (#T5, #T6 & #T9) and Δ*pldB*#T87 GFP-PldB (#T1 & #T2).

Expected protein sizes: GFP: 27 kDa; PldA-GFP: 128.3 kDa; PldB-GFP: 220.4 kDa.

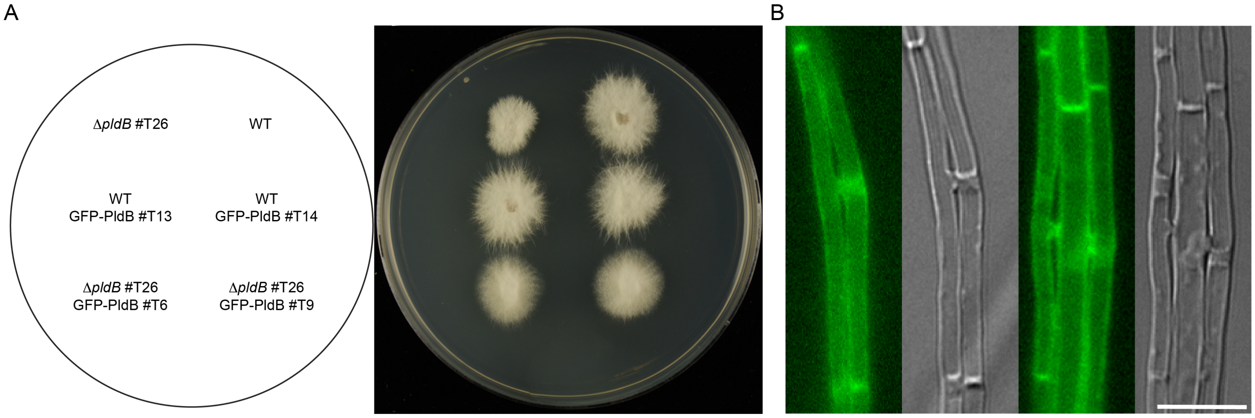

**Fig. S8.** Complementation of the culture growth phenotype of Δ*pldB* strains through the expression of GFP-PldB.

(A) Complementation of the growth defects of Δ*pldB* strains as determined by GFP-PldB expression in axenic culture after growth on potato-dextrose agar for 7 days.

(B) Complementation of the hyphal fusion defect of Δ*pldB*#T26 with GFP-PldB ((#T6 & #T9) after 7 days of growth on $H_2O$ agar. Bar=10 µm.

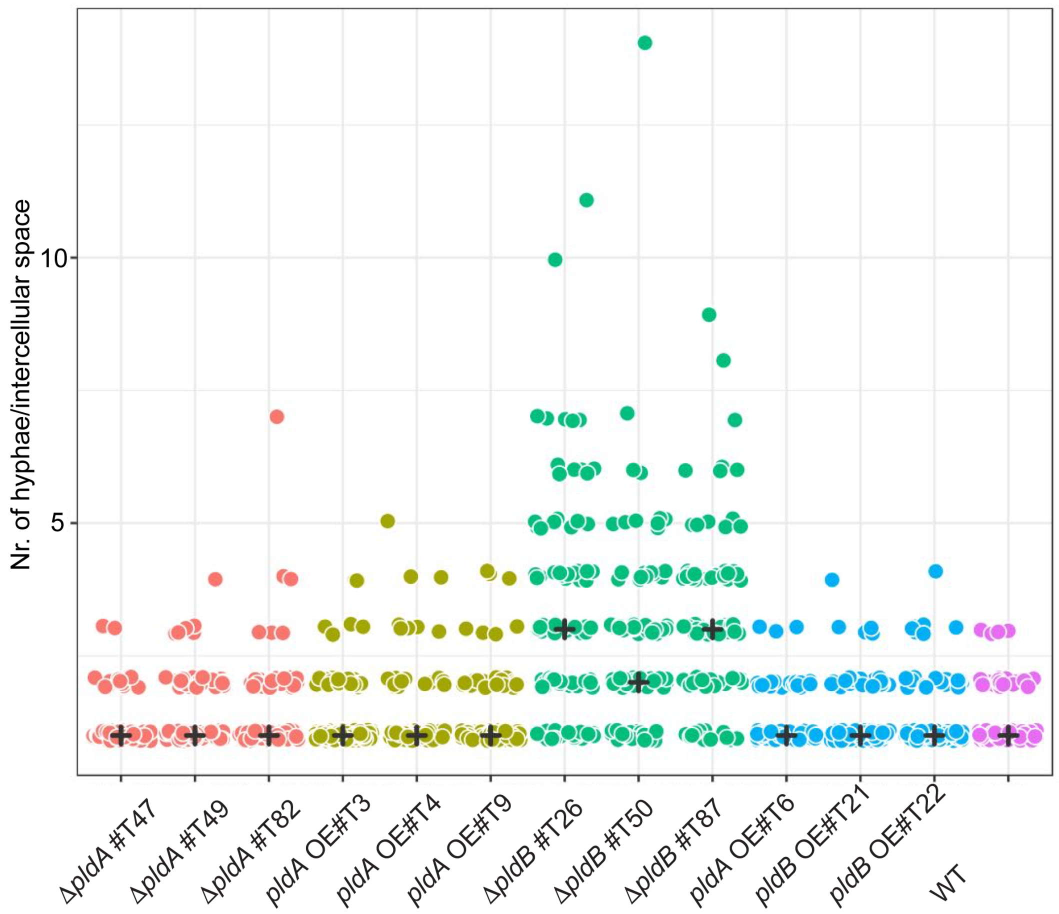

**Fig. S9.** Number of hyphae per intercellular space of *L. perenne* plants infected with *E. festucae* WT, Δ*pldA* and Δ*pldB* strains.

Number of hyphae per intercellular space were counted in fixed, toluidine blue-stained pseudostem cross sections. For each strain, 100 infected intercellular spaces were evaluated, with counting started at the outermost leaf. The data was analysed with a Dunn test, and only data obtained from plants infected with Δ*pldB* strains were significantly different from data of WT-infected plants ($P < 0.0000$). "+" indicates the average number of hyphae/intercellular space per strain.

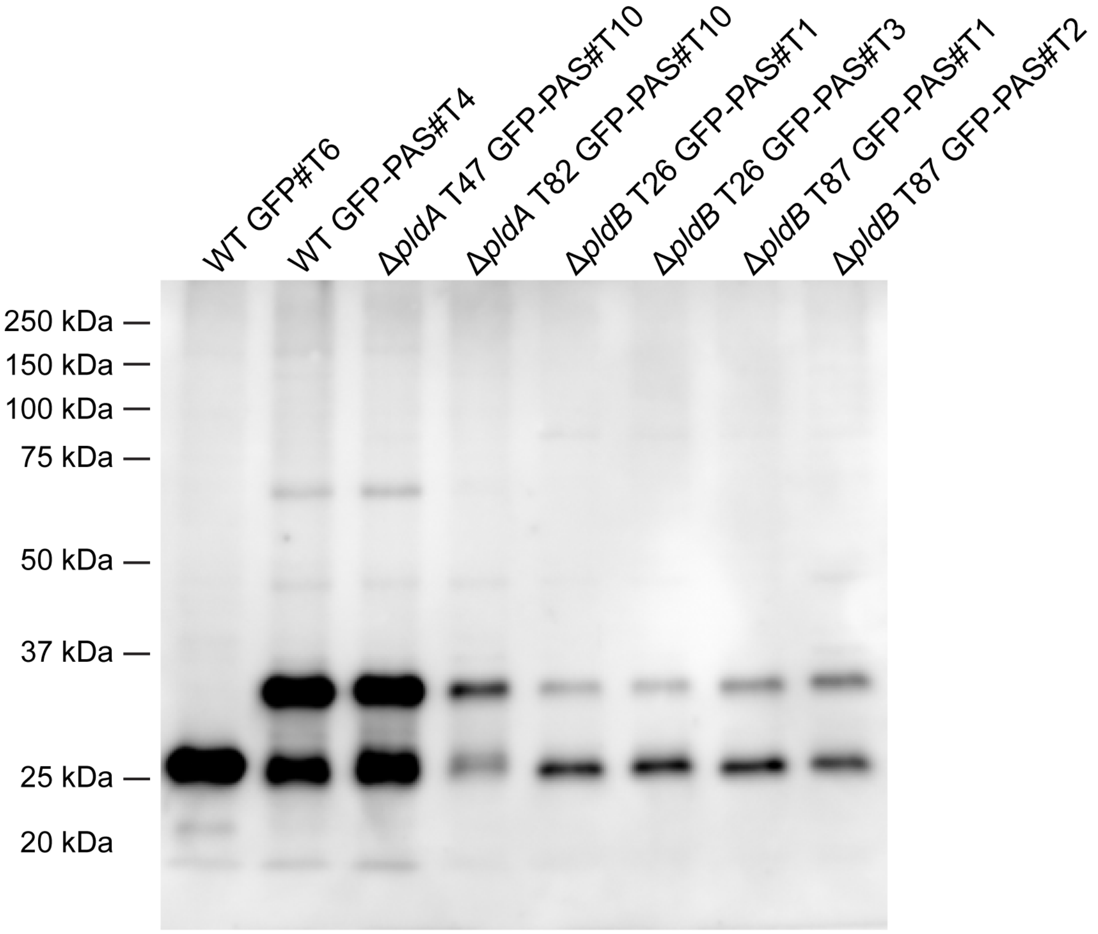

**Fig. S10.** Verification of the expression of the phosphatidic acid molecular probe in the *E. festucae* wild-type, Δ*pldA* and Δ*pldB* deletion strains by western analysis.

Phosphatidic acid biosensor (PAS)-expressing strains were grown in potato-dextrose liquid medium for three days and total protein was extracted. The western analysis was performed with the following strains: wild-type (WT) GFP, WT GFP-PAS#T4, Δ*pldA*#T47 GFP-PAS#T10, Δ*pldA*#T82 GFP-PAS#T10, Δ*pldB*#T26 GFP-PAS#T1 and #T3, Δ*pldB*#T87 GFP-PAS#T1 and #T2. From each extract, 15 µl was resolved on a 10% polyacrylamide gel. Proteins were then transferred onto PVDF membranes and probed with an α-GFP antibody. Subsequently, a secondary α-rabbit antibody conjugated to horseradish peroxidase was applied and the western blot was developed via a chemiluminescence reaction. Expected protein sizes: GFP: 27 kDa; GFP-PAS: 32 kDa.

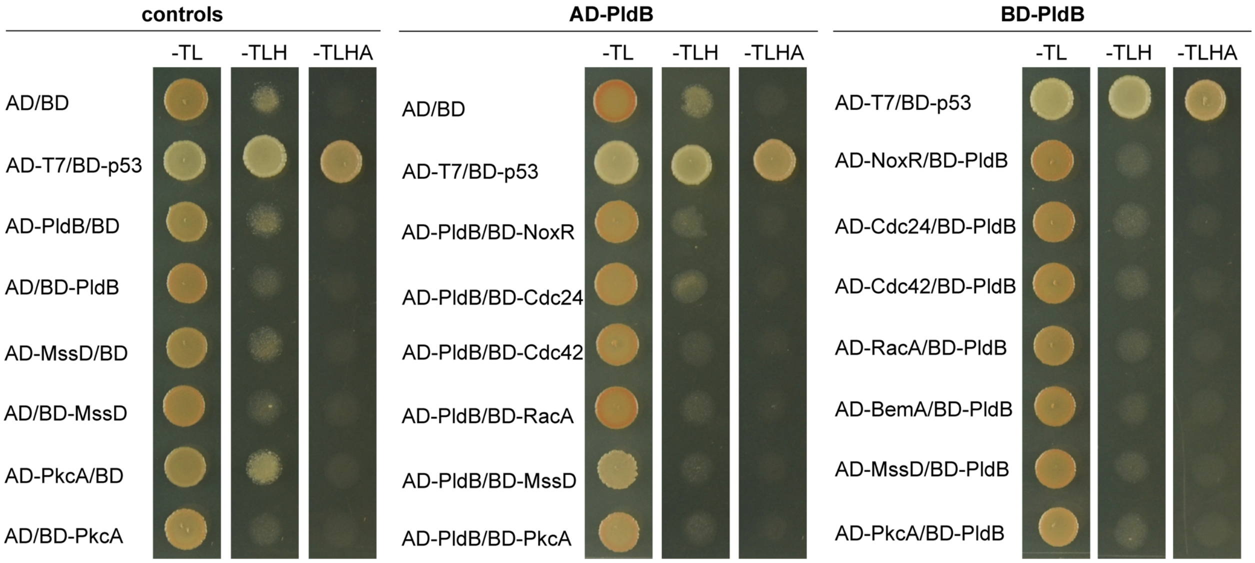

**Fig. S11.** PldB of *E. festucae* does not interact with components of the Nox complex or regulators identified in mammalian cells.

Yeast two-hybrid analysis of the interactions of PldB with *E. festucae* NoxR, Cdc24, Cdc42, RacA, BemA, PkcA (EfM3.006500) and MssD (EfM3.031950). Yeast strain AH109 was transformed with bait and prey vectors and plated on CMD plates lacking leucine and tryptophan (-L/T), leucine, tryptophan and histidine (-L/T/H) or leucine, tryptophan, histidine and adenine (-L/T/H/A). Empty pGADT7/pGBKT7 (AD/BD) were used as negative controls and the vectors pGADT7-T7/pGBKT7-p53 (AD-T7/BH-p53) as a positive control.

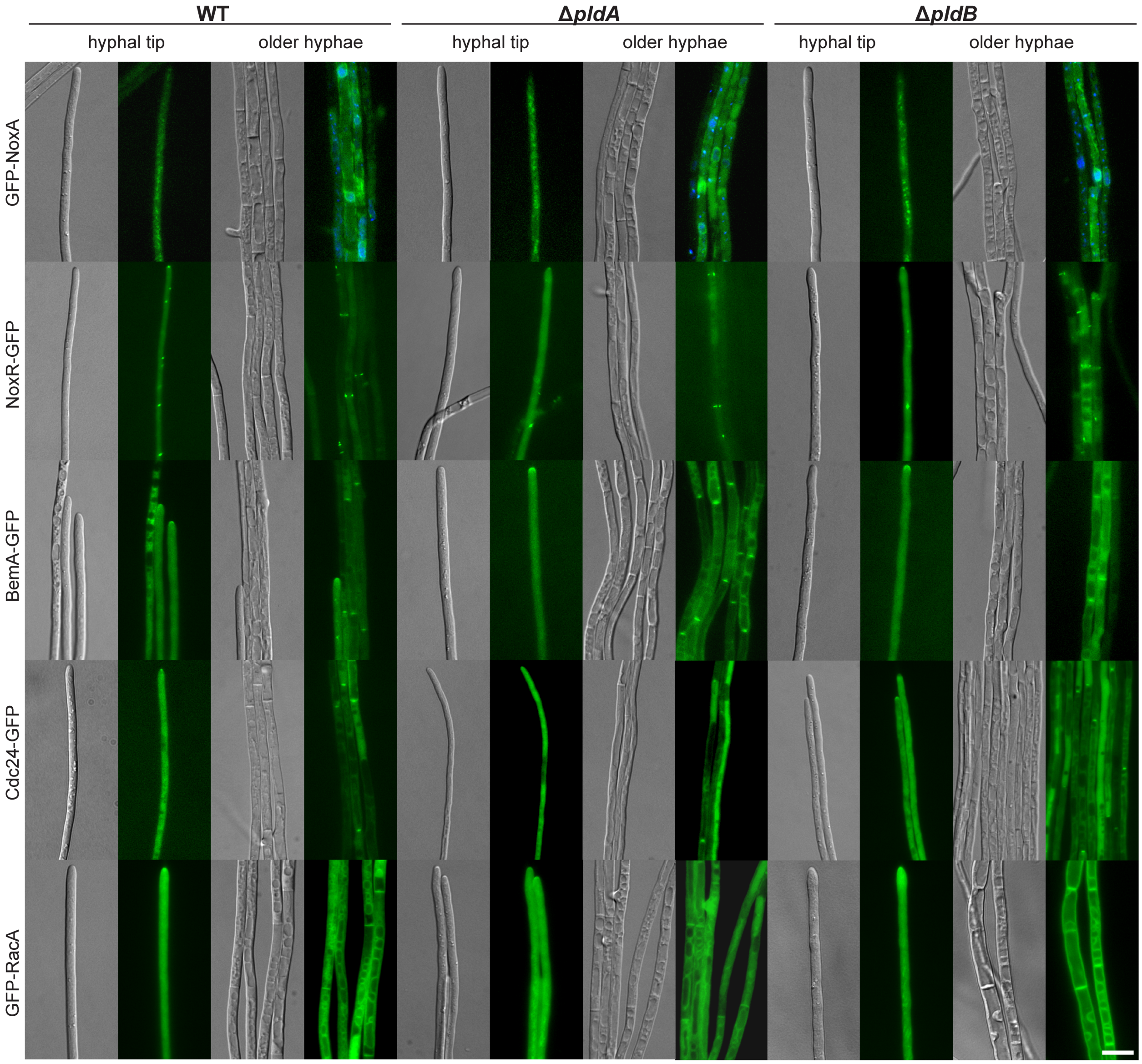

**Fig. S12.** Localisation of Nox complex components in *E. festucae* wild-type, Δ*pldA* and Δ*pldB* strains.

The localisation of NoxA, NoxR, BemA, Cdc24 and RacA GFP fusion constructs was analysed after approx. five days of incubation. Constructs were transformed into wild-type (WT) and two different Δ*pldA* (#T47 and #T82) or Δ*pldB* (#T26 and #T87) strains, and multiple transformants analysed. As not all localization constructs encode a resistance gene, strains were co-transformed with pBH12 ($Hyg^R$) or pII99 ($Gen^R$) when appropriate. Here the localization in the Δ*pldA*#T82 and Δ*pldB*#T26 strains are shown as representative of all the strains analysed. (A) Localization of GFP-NoxA (pGFP-NoxA) in WT (#T4), Δ*pldA*#T82 (#T3) and Δ*pldB*#T26 (#T1), nuclei were visualized using 0.071 mM DAPI stain. The NoxA fusion protein was found to localize to the cytoplasm, vesicles and the nuclear envelope. (B) Localisation of NoxR-GFP (pNPP9) in WT (#T1), Δ*pldA*#T82 (#T29) and Δ*pldB*#T26 (#T1); NoxR localized to hyphal tips, mobile vesicles and in puncta to either side of septa. (C) Localization of BemA-GFP (pCE139) in WT (#T23), Δ*pldA*#T82 (#T3) and Δ*pldB*#T26 (#T3); BemA was observed to localize to the PM of the hyphal tip and to the centre of septa. (D) Localization of Cdc24-GFP (pNPP6) in WT (#T33), Δ*pldA*#T82 (#T1) and Δ*pldB*#T26 (#T1); Cdc24-GFP was found to localize to puncta at the very tip of the hyphae and at the center of septa. (E) Localization of GFP-RacA (pPN154) in WT (#T6), Δ*pldA*#T82 (#T1) and Δ*pldB*#T26 (#T1); The RacA-fusion protein localized to hyphal tips, the PM and internal membranes. Bar=10 μm.

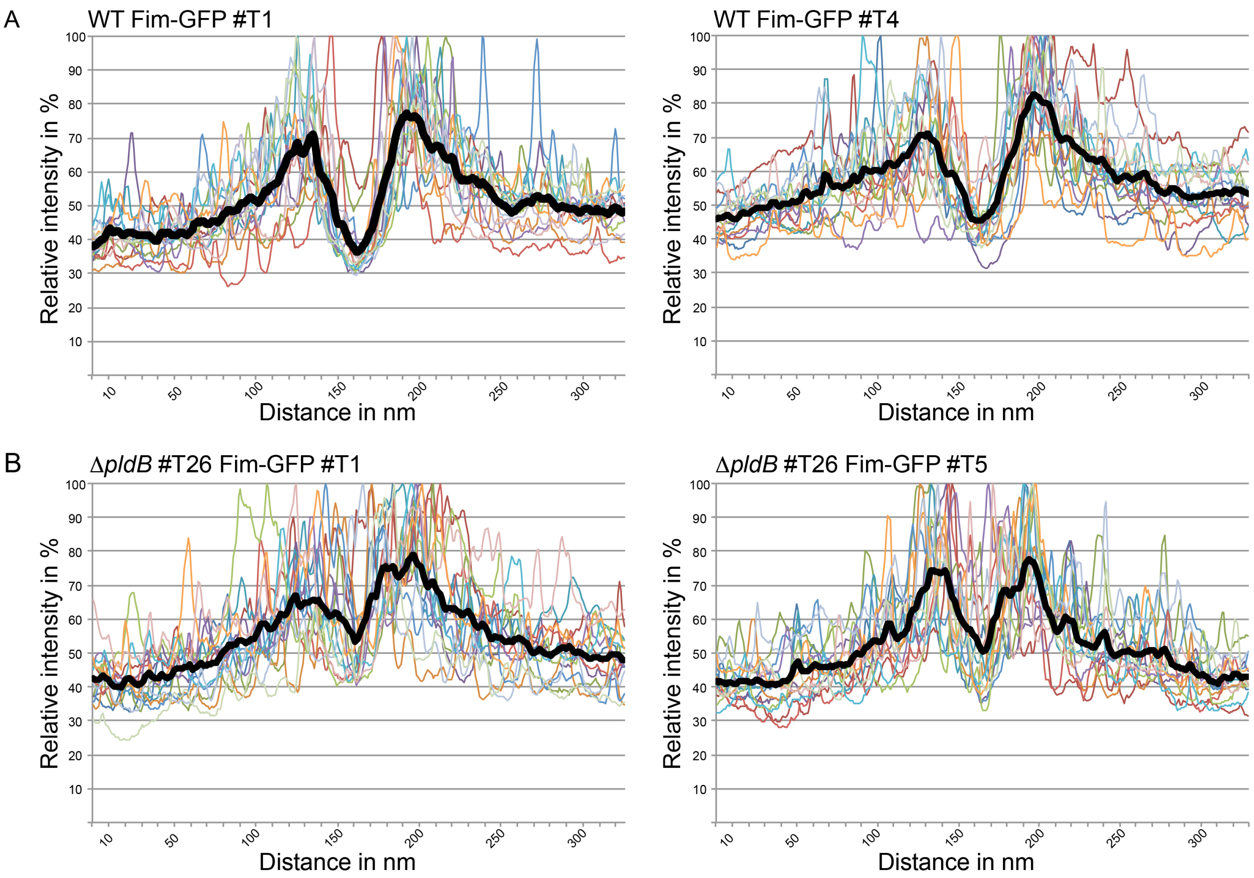

**Fig. S13.** Analysis of the localisation of Fimbrin-GFP in *E. festucae* wild-type and Δ*pldB* strains.

The strains expressing Fimbrin-GFP (pBH66) WT (T1 and T4) and Δ*pldB*#T26 (T1 and T5), were grown on 1.5% $H_2O$ agar for six days before analysis. Images of fluorescing hyphae of each strain were taken at random and the signal intensity was measured by tracing the outer edges of the hyphal tip for a distance of approx. 30 µm in total. For each strain, 15 hyphal tips were analysed using the software Fiji and relative intensities were calculated and plotted in a line graph. The black line indicates the average relative intensity. Expression of Fimbrin-GFP (pBH66) in (A) WT #T1; (B) WT #T4; (C) Δ*pldB*#T26 #T1; (D) Δ*pldB*#T26 #T5.

**Table S1:** Biological material.

| Organism/Strain | Characteristics | Reference |
|---|---|---|
| ***E. coli*** | | |
| DH5α | $F^-$, φ80*lacZ*, ΔM15, Δ(*lacZYA-argF*), U169, *recA1*, *endA1*, *hsdR17* ($r_k^-$, $m_k^-$), *phoA*, *supE44*, $\lambda^-$, *thi-1*, *gyrA96*, *relA1* | Invitrogen |
| ***S. cerevisiae*** | | |
| AH109 | MATa, *trp1-901*, leu2-3, 112, *ura3*-52, *his3-200*, *gal4*Δ, *gal80*Δ, LYS2::GAL1UAS-GAL1TATAHIS3, GAL2UAS-GAL2TATA-ADE2, URA3::MEL1 UASMEL1TATA-*lacZ*, MEL1 | Clontech |
| AH109 (AD, BD) | AH109/pGADT7 & pGBKT7; *LEU2, TRP1* | This study |
| AH109 (AD-T7, BD-p53) | AH109/pGADT7-T7 & pGBKT7-p53; *LEU2, TRP1* | This study |
| AH109 (AD-PldB, BD) | AH109/pBH60 & pGBKT7; *LEU2, TRP1* | This study |
| AH (AD, BD-PldB) | AH109/pGADT7 & pBH61; *LEU2, TRP1* | This study |
| AH109 (AD-MssD, BD) | AH109/pBH63 & pGBKT7; *LEU2, TRP1* | This study |
| AH (AD, BD-MssD) | AH109/pGADT7 & pBH62; *LEU2, TRP1* | This study |
| AH109 (AD-PkcA, BD) | AH109/pBH64 & pGBKT7; *LEU2, TRP1* | This study |
| AH (AD, BD-PkcA) | AH109/pGADT7 & pBH65; *LEU2, TRP1* | This study |
| AH109 (AD-PldB, BD-NoxR) | AH109/pBH60 & pPN113; *LEU2, TRP1* | This study |
| AH109 (AD-PldB, BD-Cdc24) | AH109/pBH60 & pPN173; *LEU2, TRP1* | This study |
| AH109 (AD-PldB, BD-Cdc42) | AH109/pBH60 & pYR9; *LEU2, TRP1* | This study |
| AH109 (AD-PldB, BD-RacA) | AH109/pBH60 & pYR7; *LEU2, TRP1* | This study |

| Organism/Strain | Characteristics | Reference |
|---|---|---|
| AH109 (AD-PldB, BD-BemA) | AH109/pBH60 & pPN172; *LEU2, TRP1* | This study |
| AH109 (AD-PldB, BD-MssD) | AH109/pBH60 & pBH62; *LEU2, TRP1* | This study |
| AH109 (AD-PldB, BD-PkcA) | AH109/pBH60 & pBH65; *LEU2, TRP1* | This study |
| AH109 (AD-NoxR, BD-PldB) | AH109/ pPN112 & pBH61; *LEU2, TRP1* | This study |
| AH109 (AD-Cdc24, BD-PldB) | AH109/ pPN160 & pBH61; *LEU2, TRP1* | This study |
| AH109 (AD-Cdc42, BD-PldB) | AH109/pYR8 & pBH61; *LEU2, TRP1* | This study |
| AH109 (AD-RacA, BD-PldB) | AH109/pYR6 & pBH61; *LEU2, TRP1* | This study |
| AH109 (AD-MssD, BD-PldB) | AH109/pBH63 & pBH61; *LEU2, TRP1* | This study |
| AH109 (AD-PkcA, BD-PldB) | AH109/pBH64 & pBH61; *LEU2, TRP1* | This study |
| ***N. crassa*** | | |
| FGSC 988 | *mat-A* | FGSC |
| FGSC 21624 | *pldA*::*hph*; *mat-A* | FGSC |
| FGSC 16300 | *pldB*::*hph*; *mat-A* | FGSC |
| ***E. festucae*** | | |
| PN2278 (Fl1) | Wild-type isolated from *Festuca longifolia* | Young *et al*. 2005 |
| PN2326 (Δ*noxA)* | WT/Δ*noxA::*P*trpC-hph;* $Hyg^R$ | Tanaka *et al*., 2006 |
| PN2469 (Δ*noxB)* | WT/Δ*noxB::*P*trpC-nptII*-T*trpC;* $Gen^R$ | Tanaka *et al*., 2006 |
| PN3272 (Δ*pldA*#T47) | WT/Δ*pldA::*P*trpC-hph;* $Hyg^R$ | This study |
| PN3273 (Δ*pldA*#T49) | WT/Δ*pldA::*P*trpC-hph;* $Hyg^R$ | This study |
| PN3274 (Δ*pldA*#T82) | WT/Δ*pldA::*P*trpC-hph;* $Hyg^R$ | This study |
| *pldA* overexpression #T3 | WT/pCE129/pII99, $Gen^R$ | This study |
| *pldA* overexpression #T4 | WT/pCE129/pII99, $Gen^R$ | This study |
| *pldA* overexpression #T9 | WT/pCE129/pII99, $Gen^R$ | This study |
| PN3275 (Δ*pldB*#T26) | WT/Δ*pldB::*P*trpC-nptII*-P*trpC;* $Gen^R$ | This study |
| PN3276 (Δ*pldB*#T50) | WT/Δ*pldB::*P*trpC-nptII*-P*trpC;* $Gen^R$ | This study |
| PN3277(Δ*pldB*#T87) | WT/Δ*pldB::*P*trpC-nptII*-P*trpC;* $Gen^R$ | This study |
| *pldB* overexpression #T6 | WT/pBH48/pII99, $Gen^R$ | This study |
| *pldB* overexpression #T21 | WT/pBH48/pII99, $Gen^R$ | This study |

| Organism/Strain | Characteristics | Reference |
|---|---|---|
| *pldB* overexpression #T22 | WT/pBH48/pII99, $Gen^R$ | This study |
| PN3284Δ*pldB*Δ*pldA*#T3 | Δ*pldB*#T87/Δ*pldA::*P*trpC-hph;* $Gen^R$; $Hyg^R$ | This study |
| PN3285Δ*pldB*Δ*pldA*#T5 | Δ*pldB*#T26/Δ*pldA::*P*trpC-hph;* $Gen^R$; $Hyg^R$ | This study |
| WT GFP-NoxA#T1 | WT/pNoxA-GFP/pBH12, $Hyg^R$ | This study |
| WT GFP-NoxA#T4 | WT/pNoxA-GFP/pBH12, $Hyg^R$ | This study |
| WT GFP-NoxA#T5 | WT/pNoxA-GFP/pBH12, $Hyg^R$ | This study |
| WT GFP-NoxA#T7 | WT/pNoxA-GFP/pBH12, $Hyg^R$ | This study |
| WT NoxR-GFP#T1 | WT/pNPP9, $Hyg^R$ | Green pers. comm. |
| WT Cdc24-GFP#T9 | WT/pNPP6, $Hyg^R$ | Green pers. comm. |
| WT Cdc24-GFP#T33 | WT/pNPP6, $Hyg^R$ | Green pers. comm. |
| WT BemA-GFP#T20 | WT/pCE139, pII99, $Gen^R$ | Green pers. comm. |
| WT BemA-GFP#T23 | WT/pCE139, pII99, $Gen^R$ | Green pers. comm. |
| WT BemA-GFP#T24 | WT/pCE139, pII99, $Gen^R$ | Green pers. comm. |
| WT RacA-GFP#T1 | WT/pPN154, $Hyg^R$ | This study |
| WT RacA-GFP#T6 | WT/pPN154, $Hyg^R$ | This study |
| WT RacA-GFP#T7 | WT/pPN154, $Hyg^R$ | This study |
| WT RacA-GFP#T8 | WT/pPN154, $Hyg^R$ | This study |
| WT PA-molecular probe #T4 | WT/pBH50, $Hyg^R$ | This study |
| WT PA-molecular probe #T7 | WT/pBH50, $Hyg^R$ | This study |
| WT PA-molecular probe #T8 | WT/pBH50, $Hyg^R$ | This study |
| WT PA-molecular probe #T9 | WT/pBH50, $Hyg^R$ | This study |
| WT PA-molecular probe #T11 | WT/pBH50, $Hyg^R$ | This study |
| WT GFP-PldA#T3 | WT/pBH37, $Gen^R$ | This study |
| WT GFP-PldA#T4 | WT/pBH37, $Gen^R$ | This study |
| WT GFP-PldA#T5 | WT/pBH37, $Gen^R$ | This study |
| WT GFP-PldA#T9 | WT/pBH37, $Gen^R$ | This study |
| WT GFP-PldA#T14 | WT/pBH37, $Gen^R$ | This study |
| WT PldA-mCherry#T1 | WT/pBH38, $Hyg^R$ | This study |
| WT GFP-PldB#T13 | WT/pBH54 $Gen^R$ | This study |
| WT GFP-PldB#T14 | WT/pBH54 $Gen^R$ | This study |
| WT GFP-PldB#T15 | WT/pBH54 $Gen^R$ | This study |
| WT GFP-PldB#T16 | WT/pBH54 $Gen^R$ | This study |
| WT GFP#T1 | WT/pBH28, $Gen^R$ | This study |
| WT GFP#T6 | WT/pBH28, $Gen^R$ | This study |
| WT mCherry#T1 | WT/pCE126, pBH12, $Hyg^R$ | This study |
| WT mCherry#T2 | WT/pCE126, pBH12, $Hyg^R$ | This study |
| WT Vps52-GFP#T15 | WT/pKG55, $Hyg^R$ | This study |
| WT Vps52-GFP#T16 | WT/pKG55, $Hyg^R$ | This study |

| Organism/Strain | Characteristics | Reference |
|---|---|---|
| WT Fim-GFP#T2 | WT/pBH66, $Hyg^R$ | This study |
| WT Fim-GFP#T4 | WT/pBH66, $Hyg^R$ | This study |
| WT Fim-GFP#T7 | WT/pBH66, $Hyg^R$ | This study |
| WT GFP-Exo70#T2 | WT/pBH67, $Hyg^R$ | This study |
| WT GFP-Exo70#T5 | WT/pBH67, $Hyg^R$ | This study |
| WT GFP-Exo70#T15 | WT/pBH67, $Hyg^R$ | This study |
| WT GFP-Rab5#T1 | WT/pBH68, $Hyg^R$ | This study |
| WT GFP-Rab5#T7 | WT/pBH68, $Hyg^R$ | This study |
| WT GFP-Rab5#T17 | WT/pBH68, $Hyg^R$ | This study |
| WT Sec3-GFP#T9 | WT/pBH69, $Hyg^R$ | This study |
| WT Sec3-GFP#T13 | WT/pBH69, $Hyg^R$ | This study |
| WT Sec3-GFP#T20 | WT/pBH69, $Hyg^R$ | This study |
| WT GFP-Rab11#T1 | WT/pBH71, $Hyg^R$ | This study |
| WT GFP-Rab11#T17 | WT/pBH71, $Hyg^R$ | This study |
| Δ*pldA*#T47 NoxR-GFP | Δ*pldA*#T47/pNPP9/pII99, $Hyg^R$, $Gen^R$ | This study |
| Δ*pldA*#T47 Cdc24-GFP#T1 | Δ*pldA*#T47/pNPP6/pII99, $Hyg^R$, $Gen^R$ | This study |
| Δ*pldA*#T47 Cdc24-GFP#T2 | Δ*pldA*#T47/pNPP6/pII99, $Hyg^R$, $Gen^R$ | This study |
| Δ*pldA*#T47 Cdc24-GFP#T4 | Δ*pldA*#T47/pNPP6/pII99, $Hyg^R$, $Gen^R$ | This study |
| Δ*pldA*#T47 BemA-GFP#T17 | Δ*pldA*#T47/pCE139/pII99, $Hyg^R$, $Gen^R$ | This study |
| Δ*pldA*#T47 BemA-GFP#T23 | Δ*pldA*#T47/pCE139/pII99, $Hyg^R$, $Gen^R$ | This study |
| Δ*pldA*#T47 RacA-GFP#T3 | Δ*pldA*#T47/pPN154/pII99, $Hyg^R$, $Gen^R$ | This study |
| Δ*pldA*#T47 RacA-GFP#T4 | Δ*pldA*#T47/pPN154/pII99, $Hyg^R$, $Gen^R$ | This study |
| Δ*pldA*#T47 RacA-GFP#T15 | Δ*pldA*#T47/pPN154/pII99, $Hyg^R$, $Gen^R$ | This study |
| Δ*pldA*#T47 PA molecular probe#T10 | Δ*pldA*#T47/pBH50/pII99, $Hyg^R$, $Gen^R$ | This study |
| Δ*pldA*#T47 Vps52-GFP#T2 | Δ*pldA*#T47/pKG55, pII99, $Hyg^R$, $Gen^R$ | This study |
| Δ*pldA*#T47 Fim-GFP#T1 | Δ*pldA*#T47/pBH66, pII99, $Hyg^R$, $Gen^R$ | This study |
| Δ*pldA*#T47 Fim-GFP#T4 | Δ*pldA*#T47/pBH66, pII99, $Hyg^R$, $Gen^R$ | This study |
| Δ*pldA*#T47 Fim-GFP#T5 | Δ*pldA*#T47/pBH66, pII99, $Hyg^R$, $Gen^R$ | This study |
| Δ*pldA*#T47 GFP-Rab5#T1 | Δ*pldA*#T47/pBH68, pII99, $Hyg^R$, $Gen^R$ | This study |

| Organism/Strain | Characteristics | Reference |
|---|---|---|
| Δ*pldA*#T47 Sec3-GFP#T2 | Δ*pldA*#T47/pBH69, pII99, $Hyg^R$, $Gen^R$ | This study |
| Δ*pldA*#T82 GFP-NoxA#T2 | Δ*pldA*#T82/pGFP-NoxA/pII99, $Hyg^R$, $Gen^R$ | This study |
| Δ*pldA*#T82 GFP-NoxA#T3 | Δ*pldA*#T82/pGFP-NoxA/pII99, $Hyg^R$, $Gen^R$ | This study |
| Δ*pldA*#T82 GFP-NoxA#T5 | Δ*pldA*#T82/pGFP-NoxA /pII99, $Hyg^R$, $Gen^R$ | This study |
| Δ*pldA*#T82 NoxR-GFP#T29 | Δ*pldA*#T82/pNPP9/pII99, $Hyg^R$, $Gen^R$ | This study |
| Δ*pldA*#T82 Cdc24-GFP#T1 | Δ*pldA*#T82/pNPP6/pII99, $Hyg^R$, $Gen^R$ | This study |
| Δ*pldA*#T82 Cdc24-GFP#T25 | Δ*pldA*#T82/pNPP6/pII99, $Hyg^R$, $Gen^R$ | This study |
| Δ*pldA*#T82 BemA-GFP#T3 | Δ*pldA*#T82/pCE139/pII99, $Hyg^R$, $Gen^R$ | This study |
| Δ*pldA*#T82 BemA-GFP#T9 | Δ*pldA*#T82/pCE139/pII99, $Hyg^R$, $Gen^R$ | This study |
| Δ*pldA*#T82 BemA-GFP#T14 | Δ*pldA*#T82/pCE139/pII99, $Hyg^R$, $Gen^R$ | This study |
| Δ*pldA*#T82 RacA-GFP#T1 | Δ*pldA*#T82/pPN154/pII99, $Hyg^R$, $Gen^R$ | This study |
| Δ*pldA*#T82 RacA-GFP#T12 | Δ*pldA*#T82/pPN154/pII99, $Hyg^R$, $Gen^R$ | This study |
| Δ*pldA*#T82 PA molecular probe#T10 | Δ*pldA*#T82/pBH50/pII99, $Hyg^R$, $Gen^R$ | This study |
| Δ*pldA*#T82 Vps52-GFP#T6 | Δ*pldA*#T82/pKG55, pII99, $Hyg^R$, $Gen^R$ | This study |
| Δ*pldA*#T82 Vps52-GFP#T7 | Δ*pldA*#T82/pKG55, pII99, $Hyg^R$, $Gen^R$ | This study |
| Δ*pldA*#T82 Fim-GFP#T3 | Δ*pldA*#T82/pBH66, pII99, $Hyg^R$, $Gen^R$ | This study |
| Δ*pldA*#T82 Fim-GFP#T7 | Δ*pldA*#T82/pBH66, pII99, $Hyg^R$, $Gen^R$ | This study |
| Δ*pldA*#T82 Fim-GFP#T13 | Δ*pldA*#T82/pBH66, pII99, $Hyg^R$, $Gen^R$ | This study |
| Δ*pldA*#T82 GFP-Exo70#T4 | Δ*pldA*#T82/pBH67, pII99, $Hyg^R$, $Gen^R$ | This study |
| Δ*pldA*#T82 GFP-Exo70#T6 | Δ*pldA*#T82/pBH67, pII99, $Hyg^R$, $Gen^R$ | This study |
| Δ*pldA*#T82 GFP-Exo70#T13 | Δ*pldA*#T82/pBH67, pII99, $Hyg^R$, $Gen^R$ | This study |
| Δ*pldA*#T82 GFP-Rab5#T10 | Δ*pldA*#T82/pBH68, pII99, $Hyg^R$, $Gen^R$ | This study |

| Organism/Strain | Characteristics | Reference |
|---|---|---|
| Δ*pldA*#T82 GFP-Rab5#T12 | Δ*pldA*#T82/pBH68, pII99, $Hyg^R$, $Gen^R$ | This study |
| Δ*pldA*#T82 GFP-Rab5#T15 | Δ*pldA*#T82/pBH68, pII99, $Hyg^R$, $Gen^R$ | This study |
| Δ*pldA*#T82 Sec3-GFP#T1 | Δ*pldA*#T82/pBH69, pII99, $Hyg^R$, $Gen^R$ | This study |
| Δ*pldA*#T82 Sec3-GFP#T10 | Δ*pldA*#T82/pBH69, pII99, $Hyg^R$, $Gen^R$ | This study |
| Δ*pldA*#T82 GFP-Rab11#T2 | Δ*pldA*#T82/pBH71, pII99, $Hyg^R$, $Gen^R$ | This study |
| Δ*pldB*#T26 complementation #T1 | Δ*pldB*#T26/pBH53 $Hyg^R$, $Gen^R$ | This study |
| Δ*pldB*#T26 complementation #T4 | Δ*pldB*#T26/pBH53 $Hyg^R$, $Gen^R$ | This study |
| Δ*pldB*#T26 complementation #T13 | Δ*pldB*#T26/pBH53 $Hyg^R$, $Gen^R$ | This study |
| Δ*pldB*#T26 GFP-NoxA#T1 | Δ*pldB*#T26/pGFP-NoxA/pBH12, $Hyg^R$, $Gen^R$ | This study |
| Δ*pldB*#T26 GFP-NoxA#T2 | Δ*pldB*#T26/pGFP-NoxA/pBH12, $Hyg^R$, $Gen^R$ | This study |
| Δ*pldB*#T26 GFP-NoxA#T3 | Δ*pldB*#T26/pGFP-NoxA/pBH12, $Hyg^R$, $Gen^R$ | This study |
| Δ*pldB* T26 NoxR-GFP#T1 | Δ*pldB*#T26/pNPP9, $Hyg^R$, $Gen^R$ | This study |
| Δ*pldB*#T26 Cdc24-GFP#T1 | Δ*pldB*#T26/pNPP6, $Hyg^R$, $Gen^R$ | This study |
| Δ*pldB*#T26 Cdc24-GFP#T2 | Δ*pldB*#T26/pNPP6, $Hyg^R$, $Gen^R$ | This study |
| Δ*pldB*#T26 BemA-GFP#T1 | Δ*pldB*#T26/pCE139, $Hyg^R$, $Gen^R$ | This study |
| Δ*pldB*#T26 BemA-GFP#T3 | Δ*pldB*#T26/pCE139/pBH12, $Hyg^R$, $Gen^R$ | This study |
| Δ*pldB*#T26 BemA-GFP#T5 | Δ*pldB*#T26/pCE139/pBH12, $Hyg^R$, $Gen^R$ | This study |
| Δ*pldB*#T26 RacA-GFP#T1 | Δ*pldB*#T26/pPN154, $Hyg^R$, $Gen^R$ | This study |
| Δ*pldB*#T26 RacA-GFP#T2 | Δ*pldB*#T26/pPN154, $Hyg^R$, $Gen^R$ | This study |
| Δ*pldB*#T26 PA molecular probe#T1 | Δ*pldB*#T26/pBH50, $Hyg^R$, $Gen^R$ | This study |
| Δ*pldB*#T26 PA molecular probe#T2 | Δ*pldB*#T26/pBH50, $Hyg^R$, $Gen^R$ | This study |
| Δ*pldB*#T26 PA molecular probe#T3 | Δ*pldB*#T26/pBH50, $Hyg^R$, $Gen^R$ | This study |

| Organism/Strain | Characteristics | Reference |
|---|---|---|
| Δ*pldB*#T26 GFP-PldB#T5 | Δ*pldB*#T26/pBH54/pBH12, $Hyg^R$, $Gen^R$ | This study |
| Δ*pldB*#T26 GFP-PldB#T6 | Δ*pldB*#T26/pBH54/pBH12, $Hyg^R$, $Gen^R$ | This study |
| Δ*pldB*#T26 GFP-PldB#T9 | Δ*pldB*#T26/pBH54/pBH12, $Hyg^R$, $Gen^R$ | This study |
| Δ*pldB*#T26 PldB-mCherry#T1 | Δ*pldB*#T26/pBH52, $Hyg^R$, $Gen^R$ | This study |
| Δ*pldB*#T26 PldB-mCherry#T2 | Δ*pldB*#T26/pBH52, $Hyg^R$, $Gen^R$ | This study |
| Δ*pldB*#T26 PldB-mCherry#T3 | Δ*pldB*#T26/pBH52, $Hyg^R$, $Gen^R$ | This study |
| Δ*pldB*#T26 GFP#T1 | Δ*pldB*#T26/pBH28, pBH12, $Hyg^R$, $Gen^R$ | This study |
| Δ*pldB*#T26 mCherry#T2 | Δ*pldB*#T26/pCE126, pBH12, $Hyg^R$, $Gen^R$ | This study |
| Δ*pldB*#T26 mCherry#T3 | Δ*pldB*#T26/pCE126, pBH12, $Hyg^R$, $Gen^R$ | This study |
| Δ*pldB*#T26 Vps52-GFP#T7 | Δ*pldB*#T26/pKG55, $Hyg^R$, $Gen^R$ | This study |
| Δ*pldB*#T26 Fim-GFP#T1 | Δ*pldB*#T26/pBH66, $Hyg^R$, $Gen^R$ | This study |
| Δ*pldB*#T26 Fim-GFP#T5 | Δ*pldB*#T26/pBH66, $Hyg^R$, $Gen^R$ | This study |
| Δ*pldB*#T26 Fim-GFP#T7 | Δ*pldB*#T26/pBH66, $Hyg^R$, $Gen^R$ | This study |
| Δ*pldB*#T26 GFP-Exo70#T2 | Δ*pldB*#T26/pBH67, $Hyg^R$, $Gen^R$ | This study |
| Δ*pldB*#T26 GFP-Exo70#T6 | Δ*pldB*#T26/pBH67, $Hyg^R$, $Gen^R$ | This study |
| Δ*pldB*#T26 GFP-Exo70#T10 | Δ*pldB*#T26/pBH67, $Hyg^R$, $Gen^R$ | This study |
| Δ*pldB*#T26 GFP-Rab5#T1 | Δ*pldB*#T26/pBH68, $Hyg^R$, $Gen^R$ | This study |
| Δ*pldB*#T26 GFP-Rab5#T2 | Δ*pldB*#T26/pBH68, $Hyg^R$, $Gen^R$ | This study |
| Δ*pldB*#T26 GFP-Rab5#T2.2 | Δ*pldB*#T26/pBH68, $Hyg^R$, $Gen^R$ | This study |
| Δ*pldB*#T26 Sec3-GFP#T6 | Δ*pldB*#T26/pBH69, $Hyg^R$, $Gen^R$ | This study |
| Δ*pldB*#T26 Sec3-GFP#T8 | Δ*pldB*#T26/pBH69, $Hyg^R$, $Gen^R$ | This study |
| Δ*pldB*#T26 Sec3-GFP#T11 | Δ*pldB*#T26/pBH69, $Hyg^R$, $Gen^R$ | This study |
| Δ*pldB*#T26 GFP-Rab11#T1 | Δ*pldB*#T26/pBH71, $Hyg^R$, $Gen^R$ | This study |

| **Organism/Strain** | **Characteristics** | **Reference** |
|---|---|---|
| Δ*pldB*#T87 complementation#T1 | Δ*pldB*#T87/pBH53, $Hyg^R$, $Gen^R$ | This study |
| Δ*pldB*#T87 complementation#T4 | Δ*pldB*#T87/pBH53, $Hyg^R$, $Gen^R$ | This study |
| Δ*pldB*#T87 complementation#T10 | Δ*pldB*#T87/pBH53, $Hyg^R$, $Gen^R$ | This study |
| Δ*pldB*#T87 GFP-NoxA#T1 | Δ*pldB*#T87/pGFP-NoxA/pBH12, $Hyg^R$, $Gen^R$ | This study |
| Δ*pldB*#T87 NoxR-GFP#T4 | Δ*pldB*#T87/pNPP9/pBH12, $Hyg^R$, $Gen^R$ | This study |
| Δ*pldB*#T87 Cdc24-GFP#T1 | Δ*pldB*#T87/pNPP6/pBH12, $Hyg^R$, $Gen^R$ | This study |
| Δ*pldB*#T87 Cdc24-GFP#T2 | Δ*pldB*#T87/pNPP6/pBH12, $Hyg^R$, $Gen^R$ | This study |
| Δ*pldB*#T87 Cdc24-GFP#T3 | Δ*pldB*#T87/pNPP6/pBH12, $Hyg^R$, $Gen^R$ | This study |
| Δ*pldB*#T87 BemA-GFP#T2 | Δ*pldB*#T87/pCE139/pBH12, $Hyg^R$, $Gen^R$ | This study |
| Δ*pldB*#T87 RacA-GFP#T1 | Δ*pldB*#T87/pPN154/pBH12, $Hyg^R$, $Gen^R$ | This study |
| Δ*pldB*#T87 RacA-GFP#T2 | Δ*pldB*#T87/pPN154/pBH12, $Hyg^R$, $Gen^R$ | This study |
| Δ*pldB*#T87 RacA-GFP#T3 | Δ*pldB*#T87/pPN154/pBH12, $Hyg^R$, $Gen^R$ | This study |
| Δ*pldB*#T87 PA molecular probe#T1 | Δ*pldB*#T87/pBH50, $Hyg^R$, $Gen^R$ | This study |
| Δ*pldB*#T87 PA molecular probe#T2 | Δ*pldB*#T87/pBH50, $Hyg^R$, $Gen^R$ | This study |
| Δ*pldB*#T87 PA molecular probe#T3 | Δ*pldB*#T87/pBH50, $Hyg^R$, $Gen^R$ | This study |
| Δ*pldB*#T87 GFP-PldB#T1 | Δ*pldB*#T87/pBH54/pBH12, $Hyg^R$, $Gen^R$ | This study |
| Δ*pldB*#T87 GFP-PldB#T2 | Δ*pldB*#T87/pBH54/pBH12, $Hyg^R$, $Gen^R$ | This study |
| Δ*pldB*#T87 Vps52-GFP#T2 | Δ*pldB*#T87/pKG55, $Hyg^R$, $Gen^R$ | This study |
| Δ*pldB*#T87 Vps52-GFP#T6 | Δ*pldB*#T87/pKG55, $Hyg^R$, $Gen^R$ | This study |
| Δ*pldB*#T87 GFP-Exo70#T5 | Δ*pldB*#T87/pBH67, $Hyg^R$, $Gen^R$ | This study |
| Δ*pldB*#T87 GFP-Exo70#T7 | Δ*pldB*#T87/pBH67, $Hyg^R$, $Gen^R$ | This study |
| Δ*pldB*#T87 GFP-Rab5#T1 | Δ*pldB*#T87/pBH68, $Hyg^R$, $Gen^R$ | This study |

| Organism/Strain | Characteristics | Reference |
|---|---|---|
| Δ*pldB*#T87 GFP-Rab5#T2 | Δ*pldB*#T87/pBH68, $Hyg^R$, $Gen^R$ | This study |
| Δ*pldB*#T87 GFP-Rab5#T3 | Δ*pldB*#T87/pBH68, $Hyg^R$, $Gen^R$ | This study |
| Δ*pldB*#T87 GFP-Rab5#T4 | Δ*pldB*#T87/pBH68, $Hyg^R$, $Gen^R$ | This study |
| Δ*pldB*#T87 GFP-Rab11#T2 | Δ*pldB*#T87/pBH71, $Hyg^R$, $Gen^R$ | This study |
| ***L. perenne*** | | |
| *L. perenne* cv. Samson | - | AgResearch |
| **Plasmid** | **Characteristics** | **Reference** |
| PN1862 (pSF15.15) | pSP72 containing 1.4-kb *Hind*III P*trpC-hph* from pCB1004 cloned into *Sma*I site. $Amp^R$; $Hyg^R$; *Nco*I-free P*trpC-hph* | S. Foster |
| PN4111 (pPN94) | pSF15.15 containing 0.8-kb *Sal*I/*Xba*I *tef* promoter in *Xho*I/*Xba*I site and 0.6-kb *EcoR*I/*Bgl*II T*trpC* in *Eco*RI/*Bgl*II site | Takemoto *et al.* 2006 |
| PN4183 (pRS426) | *ori*(f1)-l*acZ*-T7 promoter-MCS (*Kpn*I-*Sac*I)-T3 promoter-*lacI*-*ori*(pMB1)-$amp^R$-*ori* (2 micron), *URA3;* $Amp^R$ | Christianson *et al.* 1992 |
| PN1687 (pII99) | P*trpC*-*nptII*-T*trpC*, $Amp^R$/$Gen^R$ | Lara-Ortíz *et al.* 2003 |
| PN4241 (pBV579) | pAN583 containing a 0.1 kb *Bsr*GI/*Bam*HI fragment containing NLS (three tandem repeats of the nuclear localisation signal from simian virus large T-antigen) from pEBFP2-Nuc | Khang *et al*. 2010 |
| pUC57 | 5' *pldA* flank | This study |
| pPN82 | pBlueScriptII® KS(+) containing 1.4-kb *Hind*III P*trpC-hph* and P*gpd*-EGFP-T*trpC*; $Amp^R$ /$Hyg^R$ | Tanaka *et al*. 2006 |
| pGADT7 | GAL4-AD, SV40-NLS, LEU2, $Amp^R$ | Takara Bio USA Inc. |
| pGBKT7 | GAL4-BD, SV40-NLS, TRP1, $Kan^R$ | Takara Bio USA Inc. |
| pGADT7-7 | pGBKT& containing SV40large T-$antigen_{aa86-708}$ cDNA, $Amp^R$ | Takara Bio USA Inc. |
| pGBKT7-p53 | pGBKT7 containing murine $p53_{aa72-390}$ cDNA, $Kana^R$ | Takara Bio USA Inc. |
| pPN112 | pGAD-T7containing a *noxR* insert, $Amp^R$ | Takemoto *et al.*, 2011 |

| Organism/Strain | Characteristics | Reference |
|---|---|---|
| pPN113 | pGBK-T7 containing a *noxR* insert, Kan$^R$ | Takemoto *et al.*, 2011 |
| pYR6 | pGAD-T7containing a *racA* insert, Amp$^R$ | Becker *et al.*, (in preparation) |
| pYR7 | pGBK-T7 containing a *racA* insert, Kan$^R$ | Becker *et al.*, (in preparation) |
| pPN160 | pGAD-T7containing a *cdc24* insert, Amp$^R$ | Takemoto *et al.*, 2011 |
| pPN173 | pGBK-T7 containing a *cdc24* insert, Kan$^R$ | Takemoto *et al.*, 2011 |
| pYR8 | pGAD-T7containing a *cdc42* insert, Amp$^R$ | Becker *et al.*, (in preparation) |
| pYR9 | pGBK-T7 containing a *cdc42* insert, Kan$^R$ | Becker *et al.*, (in preparation) |
| pPN172 | pGBK-T7 containing a *bemA* insert, Kan$^R$ | Takemoto *et al.*, 2011 |
| pPN154 | pPN94 containing 0.7-kb *Xba*I/*Xho*I *egfp* and 0.6-kb *Xho*I/*Not*I *racA* in XbaI/NotI site | Takemoto *et al.*, 2011 |
| pNPP6 | pPN94 containing 3.0-kb *Xba*I/*Cla*I *cdc*24 and 0.7-kb *Cla*I/*Not*I 3GA-*egfp* in *Xba*I/*Not*I site. | Takemoto *et al.*, 2011 |
| pNPP9 | pPN94 containing 1.6-kb *Eco*RI/*Cla*I *noxR* and 0.7-kb ClaI/NotI 3GA-*egfp* in *EcoR*I/*Not*I site. | Takemoto *et al.*, 2011 |
| pCE139 | pRS425 containing *bemA-egfp* | Eaton, pers. comm. |
| pGFP-NoxA | pRS425 containing *egfp-noxA* | Eaton, pers. comm. |
| pCE131 | pRS426 containing *pldA* deletion construct, Hyg$^R$ | This study |
| pCE121 | pRS426 containing *pldB* deletion construct, Gen$^R$ | This study |
| pCE129 | pRS426 containing a P*gdp-pldA*-T*trpC* overexpression construct | This study |
| pCE126 | pRS426 containing P*gpd*-mCherry-T*trpC* insert | Candy, 2019 |
| pKG55 | pPN94 containing P*tef-vps52-gfp*-T*trpC* (*vps52*: 2167 bp), Hyg$^R$ | Green, pers. comm. |
| pBH12 | pPN94 containing 723 bp *B. cinerea tub* terminator insert amplified from pNR1 downstream of *hph*; Hyg$^R$ | Hassing *et al.*, 2019 |
| pBH16 | pBH12 containing mCherry-NLS amplified from pBV579, Hyg$^R$ | This study |
| pBH28 | pBH12 containing *nptII* replacing *hph* and *gfp;* Gen$^R$ | Hassing *et al.*, 2019 |
| pBH37 | pBH28 containing P*gpd-gfp-pldA*-T*trpC*, Gen$^R$ | This study |

| Organism/Strain | Characteristics | Reference |
|---|---|---|
| pBH38 | pBH12 containing P*gpd-pldA-mCherry*-T*trpC*, Hyg$^R$ | This study |
| pBH48 | pRS426 containing a P*gdp-pldB*-T*trpC* overexpression construct | This study |
| pBH50 | pBH12 containing the PA-localisation construct (*gfp*, bp 151-273 of *S. cerevisiae SPO20*), Hyg$^R$ | This study |
| pBH52 | pBH12 containing P*gpd-pldB-mCherry*-T*trpC*, Hyg$^R$ | This study |
| pBH53 | pBH12 containing a *pldB* complementation construct | This study |
| pBH54 | pBH28 containing P*gpd-gfp-pldB*-T*trpC*, Gen$^R$ | This study |
| pBH60 | pGADT7 containing a *pldB* insert, Amp$^R$ | This study |
| pBH61 | pGBK-T7 containing a *pldB* insert, Kan$^R$ | This study |
| pBH62 | pGBK-T7 containing a *mssD* insert, Kan$^R$ | This study |
| pBH63 | pGAD-T7 containing a *mssD* insert, Amp$^R$ | This study |
| pBH64 | pGAD-T7 containing a *pkcA* insert, Amp$^R$ | This study |
| pBH65 | pGBK-T7 containing a *pkcA* insert, Kan$^R$ | This study |
| pBH66 | pBH12 containing P*tef-fim-gfp*-T*trpC* (*fim*: 2231 bp), Hyg$^R$ | This study |
| pBH67 | pBH12 containing P*tef-gfp-exo70*-T*trpC* (*exo70*: 1955 bp), Hyg$^R$ | This study |
| pBH68 | pBH12 containing P*tef-gfp-rab5*-T*trpC* (*rab5*: 966 bp), Hyg$^R$ | This study |
| pBH69 | pBH12 containing P*tef-sec3-gfp*-T*trpC* (*sec3*: 4618 bp), Hyg$^R$ | This study |
| pBH71 | pBH12 containing P*tef-gfp-rab11*-T*trpC* (*rab11*: 953 bp), Hyg$^R$ | This study |

**Table S2:** Primer sequences.

| Name | Sequence (5'-3') | Purpose |
|---|---|---|
| PldA1 | GCGGATAACAATTTCACACAGGAAACAGCCCCTA GACCTGACGGAAGG | *pldA* deletion, 3' fragment |
| PldA2 | AAATGCTCCTTCAATATCAGTTCCAAGCTCCCTCTG AGTTGGTGACGAC | *pldA* deletion, 3' fragment |
| PldA5 | CCAGCACTCGTCCGAGGGCAAAGGAATAGAAGTC ATGTTAGCCGTTGCTG | *pldA* deletion, 5' fragment |
| PldA6 | GTAACGCCAGGGTTTTCCCAGTCACGACATGTTTT CGGATTCGGCG | *pldA* deletion, 5' fragment |
| PldA7 | ACAGCTACCCCGCTTGAGCAGACATCACCATGTCC AAGACTGGGGAAGTTTC | *pldA* overexpression |

| Name | Sequence (5'-3') | Purpose |
|---|---|---|
| PldA8 | AGATTCGTCAAGCTGTTTGATGATTTCAGTTAGGT GTACACACTTTCTGTCC | *pldA* overexpression |
| PldA9 | TTTGGTCGTTGGGACGCTCGC | *pldA* sequencing |
| PldA10 | TCAGCAGCCTTCCTTTGTGAG | *pldA* sequencing |
| PldA11 | ACAAGTTGAGCGAATCTAAGC | Verification of *pldA* deletion |
| PldA12 | AAGTCGTCACCAACTCAGAGG | Verification of *pldA* deletion |
| PldA13 | CCCATTTTACCACTTGGCGTC | Verification of *pldA* deletion |
| PldA14 | CTCAAAGTAGCACACTCCTCG | Verification of *pldA* deletion |
| PldB1 | GCGGATAACAATTTCACACAGGAAACAGCAGTTTG GCCCTGTTTGTTGC | *pldB* deletion, 3' fragment |
| PldB2 | CCAAGCCCAAAAAGTGCTCCTTCAATATCCCAGCA TGGAAGGCAAATCG | *pldB* deletion, 3' fragment |
| PldB3 | CTCGAAAATCATTCCTACTAAGATGGGTACTTGTA GGCAGTGGTGAGCT | *pldB* deletion, 5' fragment |
| PldB4 | GGTAACGCCAGGGTTTTCCCAGTCACGACCATAGA CGCTGTATCCGCCA | *pldB* deletion, 5' fragment |
| PldB5 | ACAGCTACCCCGCTTGAGCAGACATCACCATGCCT TTTGATTTCAGCATG | *pldB* overexpression, 5' fragement |
| PldB6 | AGATTCGTCAAGCTGTTTGATGATTTCAGCTAGTT GCTGTGGAACGATC | *pldB* overexpression, 3' fragement |
| PldB7 | CTGCTATCAGTCAGAAGTGGCGTCTTGACAGACTG TTGC | *pldB* overexpression, 3' fragement |
| PldB8 | ACGCCACTTCTGACTGATAGCAGCCGGCCGGCGCA TGTAAAGTTCAGG | *pldB* overexpression, 5' fragement |
| PldB9 | CTTCCGCTCGTTTTGGTAGTC | *pldB* sequencing |
| PldB10 | ACTCCGCTTGCCATACCATCC | *pldB* sequencing |
| PldB11 | GACTCCATAAATGAAATGCTC | *pldB* sequencing |
| PldB12 | CACTGGTTGAGCGTATCATTC | *pldB* sequencing |
| PldB13 | CCTTGGACACATCGGCAGGTC | *pldB* sequencing |
| PldB14 | ACAACCACTTGACCGTCCAGC | Verification of *pldB* deletion |
| PldB15 | TTGAGTTGCCAGAACCCAGAG | Verification of *pldB* deletion |
| PldB16 | GCAACGTGTATCTTTAGCTCC | Verification of *pldB* deletion |
| PldB17 | ATAAGTGTGCTGAGATTGCAC | Verification of *pldB* deletion |
| BH59 | TATCCACGCCCTCCTACATC | Verification of *pldA* deletion |

| Name | Sequence (5'-3') | Purpose |
|---|---|---|
| BH60 | GTTGACGGCAATTTCGATG | Verification of *pldA* deletion |
| BH75 | CAGTGAGCGAGGAAGCGGAAGGCTTGCTTAGCTT GATATCTG | pBH12 amplification |
| BH76 | CTGAAATCATCAAACAGCTTG | pBH12 amplification |
| BH78 | GTGTCACCTAAATCGTATGTG | *pldB* complementation construct |
| BH79 | GGAGGTGGAGGTTCTGGTGGAGGTGGATCTATGGT GAGCAAGGGCGAGGA | *mCherry* with linker forward |
| BH114 | GTGACACTATAGAACTCGACGAATTCCCTTGTATC TCTACA | Amplification of P*gdp*+*gfp* forward |
| BH191 | CATAGATCCACCTCCACCAGAACCTCCACCTCCCT TGTACAGCTCGTCCATGC | Amplification of P*gdp*+*gfp* with linker reverse |
| BH192 | GGAGGTGGAGGTTCTGGTGGAGGTGGATCTATGTC CAAGACTGGGGAAGTTTC | *pldA* with linker forward |
| BH193 | GTCGAGTTCTATAGTGTCACC | Amplification of vector backbone |
| BH196 | GGTGATGTCTGCTCAAGCGG | Amplification of vector backbone |
| BH197 | CTTCCGCTTCCTCGCTCACTG | Amplification of vector backbone |
| BH198 | CCGCTTGAGCAGACATCACCATGTCCAAGACTGGG GAAGTTTC | *pldA* forward with overhang to P*gdp* |
| BH199 | CAAGCTGTTTGATGATTTCAGTTAAGATCTGTACA GCTCGTCCATGCCG | *mCherry* reverse overhang to pBH12 |
| BH200 | CATAGATCCACCTCCACCAGAACCTCCACCTCCGG TGTACACACTTTCTGTCCA | *pldA* reverse with overhang to linker |
| BH202 | CATAGATCCACCTCCACCAGAACCTCCACCTCCGT TGCTGTGGAACGATCAGTTG | *pldB* reverse with linker |
| BH206 | CATAGATCCACCTCCACCAGAACCTCCACCTCCGG TGTACACACTTTCTGTCCA | *npt2* reverse for split marker deletion |
| BH207 | CATAGATCCACCTCCACCAGAACCTCCACCTCCGG TGTACACACTTTCTGTCCA | *npt2* forward for split marker deletion |
| BH220 | CACATACGATTTAGGTGACACTAGCAACGTGTATC TTTAGCTC | *pldB* forward for complementation |
| BH222 | GACGCCACTAAGACTAGTTGACTGAAATCATCAAA CAGCTTG | amplification of pBH50 backbone |
| BH223 | CACCTGCTCCGGCACCAGCGCCCTTGTACAGCTCG TCCATGCC | amplification of pBH50 backbone |
| BH224 | CCGCTTGAGCAGACATCACCATGTCACCGGCAGAA CATGG | *pldB* reverse with overhang to P*gdp* |

| Name | Sequence (5'-3') | Purpose |
|---|---|---|
| BH235 | GACCGGCAGATCTATTGTATACCCCTTCTGCTTTTG CTGGAA | *pldB* reverse for complementation |
| BH236 | GTATACAATAGATCTGCCGGTC | backbone amplification forward without terminator |
| BH239 | TTCTGGTGGAGGTGGATCTATGTCACCGGCAGAAC ATGGTAA | *pldB* forward overhang to linker |
| BH241 | ATGGCCATGGAGGCCAGTGAATTCATGTCACCGGC AGAACATGG | *pldB* forward for pGAD-T7 |
| BH242 | CATCTGCAGCTCGAGCTCGATGGATCCCTAGTTGT AGATTGACAGCGG | *pldB* reverse for pGAD-T7 |
| BH243 | GCATATGGCCATGGAGGCCGAATTCATGTCACCGG CAGAACATGG | *pldB* forward for pGBK-T7 |
| BH244 | TGCTAGTTATGCGGCCGCTGCAGGCTAGTTGTAGA TTGACAGCGG | *pldB* reverse for pGBK-T7 |
| BH245 | GCATATGGCCATGGAGGCCGAATTCATGCCCTCGT TTCTCAACGAC | *mssD* forward for pGAD-T7 |
| BH246 | TGCTAGTTATGCGGCCGCTGCAGGCTAAACCATGC GTACGCCGCTG | *mssD* reverse for pGAD-T7 |
| BH247 | ATGGCCATGGAGGCCAGTGAATTCATGCCCTCGTT TCTCAACGAC | *mssD* forward for pGBK-T7 |
| BH248 | CATCTGCAGCTCGAGCTCGATGGATCCCTAAACCA TGCGTACGCCGCTG | *mssD* reverse for pGBK-T7 |
| BH249 | ATGGCCATGGAGGCCAGTGAATTCATGAGCGAAG ACGAAAAAAAG | *pkcA* forward for pGAD-T7 |
| BH250 | CATCTGCAGCTCGAGCTCGATGGATCCTCATTCAA AATCCGCGGTGTAC | *pkcA* reverse for pGAD-T7 |
| BH251 | GCATATGGCCATGGAGGCCGAATTCATGAGCGAA GACGAAAAAAAG | *pkcA* forward for pGBK-T7 |
| BH252 | TGCTAGTTATGCGGCCGCTGCAGGTCATTCAAAAT CCGCGGTGTAC | *pkcA* reverse for pGBK-T7 |
| BH256 | GGAGGTGGAGGTTCTGGTGGAGGTGGATCTATGGT GAGCAAGGGCGAGGAG | amplification of *gfp* for C-terminal fusion protein |
| BH257 | CATACATCACCGTCAAACCTCATGAATGTCCTCAA ATTGC | *fim* forward for pBH67 |
| BH258 | AGATCCACCTCCACCAGAACCTCCACCTCCCATCT TCTCATGCGTCGCC | *fim* reverse for pBH67 |
| BH263 | CATACATCACCGTCAAACCTCATGGTGAGCAAGGG CGAG | *gfp* forward for N-terminal fusion protein |
| BH264 | GGAGGTGGAGGTTCTGGTGGAGGTGGATCTATGGC AGCACGAGGCCCT | *rab5* forward for pBH68 |

| Name | Sequence (5'-3') | Purpose |
|---|---|---|
| BH265 | CAAGCTGTTTGATGATTTCAGTTAACAGCTGCACG GCCCGCT | *rab5* reverse for pBH68 |
| BH266 | GGAGGTGGAGGTTCTGGTGGAGGTGGATCTATGGC GGTTGGCTTATCAAATAC | *exo70* forward for pBH67 |
| BH267 | CAAGCTGTTTGATGATTTCAGTTATGAAGCCAGAC TCAGG | *exo70* reverse for pBH67 |
| BH268 | CATACATCACCGTCAAACCTCATGGACCGCGCAAA TGGGG | *sec3* forward for pBH69 |
| BH269 | AGATCCACCTCCACCAGAACCTCCACCTCCTCGGA ACGCCGTCTTCACGTC | *sec3* reverse for pBH69 |
| BH276 | GATCTGCTCTTTCTTTTCTC | sequencing forward out of P*tef* |
| BH277 | GATCACATGGTCCTGCTG | sequencing forward out of *gfp* |
| BH278 | GAACTTCAGGGTCAGCTTG | sequencing reverse out of *gfp* |
| BH279 | GGAGGTGGAGGTTCTGGTGGAGGTGGATCTATGGC CAACGACGAATACGATG | *rab11* forward for pBH71 |
| BH280 | CAAGCTGTTTGATGATTTCAGTTAACAGCACTTGC CACCCTTG | *rab11* reverse for pBH71 |
| TtrpC-F | CTGAAATCATCAAACAGCTTGACG | backbone for overexpression construct |
| pRS426-TtrpC-R | GCGGATAACAATTTCACACAGGAAACAGCCCATCT TAGTAGGAATGATTTTCG | backbone for overexpression construct |
| pRS426-Pgdp-F | GTAACGCCAGGGTTTTCCCAGTCACGACGAATTCC CTTGTATCTCTACA | backbone for overexpression construct |
| pRS426-Pgdp-R | GGTGATGTCTGCTCAAGCGGGGTAG | backbone for overexpression construct |
| pRS426_F | GCTGTTTCCTGTGTGAAATTG | *pRS426* backbone |
| pRS426_R | GGGTTTTCCCAGTCACGAC | *pRS426* backbone |
| hph_F | AGCTTGGAACTGATATTGAAGG | P*trpC-hph* for deletion constructs |
| hph_R | CGTCCGAGGGCAAAGGAATAG | P*trpC-hph* for deletion constructs |
| nptIIF | GATATTGAAGGAGCACTTTTTG | P*trpC-npt2*-T*trpC* for deletion constructs |

| Name | Sequence (5'-3') | Purpose |
|---|---|---|
| nptIIR | CTACCCATCTTAGTAGGAATG | P*trpC-npt2*-T*trpC* for deletion constructs |
| **Primers used for RT-qPCR.** | | |
| BH185 | CGTCAAAGCTGGCCACATTT | *pldA* for |
| BH186 | TGCAAGAACTCCATGGGCAT | *pldA* rev |
| BH187 | CGGCTTTCCTATCGACGACA | *pldB* for |
| BH188 | ACCATTTTGCTCCTCCGTGA | *pldB* rev |

**Methods S1:** Statistical analysis.

**Methods S2** Construct design.

For generation of the *pldA* replacement construct (pCE131) a 1,439 bp PCR fragment 5' of *pldA* and a 1292 bp PCR fragment 3' of *pldA* were amplified using Phusion polymerase with the primer combination PldA6/PldA5 and PldA1/PldA2. The 5' fragment was unable to be amplified directly from *E. festucae* genomic DNA so was synthesized by GenScript as an insert in pUC57. For the generation of the *pldB* replacement construct, pCE121, a 1,133 bp PCR fragment 5' of *pldB* and a 943 bp PCR fragment 3' of *pldB* were amplified as described making use of the primer combinations PldB4/PldB3 and PldB1/PldB2. P*trpC-hph* (1394 bp) cassette for the *pldA* replacement construct was amplified from pSF15.15 with the primer combination hph_F/hph_R and P*trpC-npt2*-T*trpC* (1,741 bp) for the *pldB* replacement construct from pII99 with the primers nptIIF/nptIIR. The backbone pRS426 was amplified with the primers pRS426_F/pRS426_R.

The *pldB* complementation construct (pBH53) was generated from three PCR fragments: the backbone was amplified from pBH12 with BH236/BH78 (4,540 bp) and the two *pldB* fragments were amplified with BH220/PldB8 (3,843 bp) and PldB7/BH235 (3,335 bp) from genomic DNA. The first of the *pldB* fragments included 1548 bp of the promoter region and the latter 357 bp of the terminator region.

For the generation of the *pldA* overexpression construct (pCE129), *pldA* (2,906 bp) was amplified from WT genomic DNA with the primers PldA7/PldA8. For the *pldB* overexpression construct pBH48, *pldB* was amplified in two fragments with the following combinations:

PldB5/PldB8 (2,774 bp) and PldB6/PldB7 (2,985 bp). The pRS426 backbone was amplified as described above and P*gpd* and T*trpC* with pRS426-Pgdp-F/pRS426-Pgdp-R (2,310 bp) and TtrpC-F/ pRS426-TtrpC-R (569 bp), from pPN82 and pII99 respectively.

The mCherry fusion protein encoding constructs were assembled from three backbone fragments and the *pldA* (pBH38) or *pldB* (pBH52) coding region. These included: a 2,817 bp fragment amplified from pBH16 with BH76/BH75, containing T*trpC* and a *hph* cassette, from a 4,608 bp fragment amplified from pBH28 with BH197/196 containing genes for replication in *E. coli* and P*gpd* from a 765 bp fragment amplified from pBV579 with BH79/BH199 encoding *mCherry*. *pldA* was amplified from gDNA with BH198/BH200 (2,956 bp) and *pldB* in two fragments with BH224/PldB8 and PldB7/BH202 (2,986 bp). For the N-terminal GFP fusion protein constructs the backbone was amplified in two fragments from pBH28. One fragment was 4,810 bp in length and was amplified with BH76/BH193 and the other one was 3,080 bp in length and was amplified with BH114/BH191. Genomic DNA was used to amplify *pldA* (2,965 bp) with BH192/PldA8 and *pldB* (5,198 bp) in two fragments with BH239/PldB8 and PldB7/PldB6. Gibson assembly resulted in the plasmids pBH37 (GFP-PldA) and pBH54 (GFP-PldB). All designed fusion protein constructs encode a $(GGGS)_2$ linker between the protein of interest and the fluorophore.

The Y2H vectors were generated as follows: *mssD* (EfM3.031950) was amplified from cDNA with the primers BH247/BH248 (2,940 bp) for insertion into pGADT7 (pBH63) and with BH245/BH246 (2,938 bp) for insertion into pGBKT7 (pBH62); *pkcA* (EfM3.006500), was amplified from cDNA with the primers BH249/BH250 (3,474 bp) for insertion into pGADT7 (pBH64) and with BH251/BH252 (3,472 bp) for insertion into pGBKT7 (pBH65); *pldB* was amplified from genomic DNA as amplifications from cDNA were unsuccessful, however the position of introns had previously been confirmed via PCR on short fragment cDNA. The gene was amplified in two fragments with the primers BH241/PldB8 (2,292 bp) and PldB7/BH242 (2,924 bp) for insertion into pGADT7 (pBH60) and with BH243/PldB8 (2,268 bp) and PldB7/BH244 (2,921 bp) for insertion into pGBKT7 (pBH61). The backbone vectors were amplified in *E. coli* and subsequently pGADT7 was restriction enzyme digested with *Eco*RI and *Bam*HI and pGBKT7 with *Eco*RI and *Pst*I. The Gibson-assembly reaction was performed following the gel extraction and purification of all fragments.

For the generation of the PA biosensor (pBH50) the genetic sequence of PA binding region of Spo20 (aa 51-91) including a 5xGA linker and a stop codon (total 156 bp) was ordered from GenScript. The backbone was amplified in two fragments from pBH28 with the primers BH222/BH75 (2,838 bp) and BH197/BH223 (5,326 bp).

Constructs for the localisation of Fimbin, Rab5, Rab11, Sec3 and Exo70, were generated by Gibson assembly from three fragments, respectively. The vector backbone was amplified from pBH12 with pBH76/pBH77 (5870 bp), *gfp* for C-terminal fusion proteins with BH256/BH115 (771 bp) and *gfp* for N-terminal fusion proteins with BH263/BH191 (768 bp). Primers for the amplification of *gfp* included overhangs to the vector backbone and the 2x(GGGGS) linker. The gene *fim* was amplified from WT genomic DNA with BH257/258 (2282 bp), *exo70* with BH266/267 (1997 bp), *rab5* with BH264/BH265 (1008 bp), *sec3* with BH268/BH267 (4669 bp) and *rab11* with BH279/BH280 (1004 bp). Assembly resulted in the plasmids pBH66, pBH67, pBH68, pBH69 and pBH71 respectively.
