## Supplemental Methods 1 for "Phosphatidic acid produced by phospholipase D is required for hyphal cell-cell fusion and fungal-plant symbiosis"

### Methods S1

#### Introduction

This document summarises and presents a reproducible record of the statistical analyses presented in the main body of the paper. Because several of the datasets produced in this work have unusual distributions or require correction for multiple testing, we focus on describing the rationale for the particular approaches we have applied.

##### Frequency of hyphal fusions among different strains.

We observed no hyphal fusions in our *pldB* deletion strains, while several such fusions can be found in all other strains. To quantify this phenomenon we counted the number of fusions observed in one field of view at 400-fold magnification. Each strain was analysed in triplicates, where one replicate consisted of the number of fusions observed in 10 fields of view.

```
fusions <- read.csv('clean_data/fusion_counts.csv')
ggplot(fusions, aes(genotype, n_fusions)) +
    geom_hline(yintercept=1.73) +
    stat_summary(aes(colour=is_pldb_KO)) +
    scale_colour_brewer(palette="Set1") +
    ylab("Number of hyphal fusions per field of view") +
    xlab("Genotypes") +
    coord_flip() +
    theme(axis.text.y=element_blank(),
          axis.ticks.y=element_blank())
```

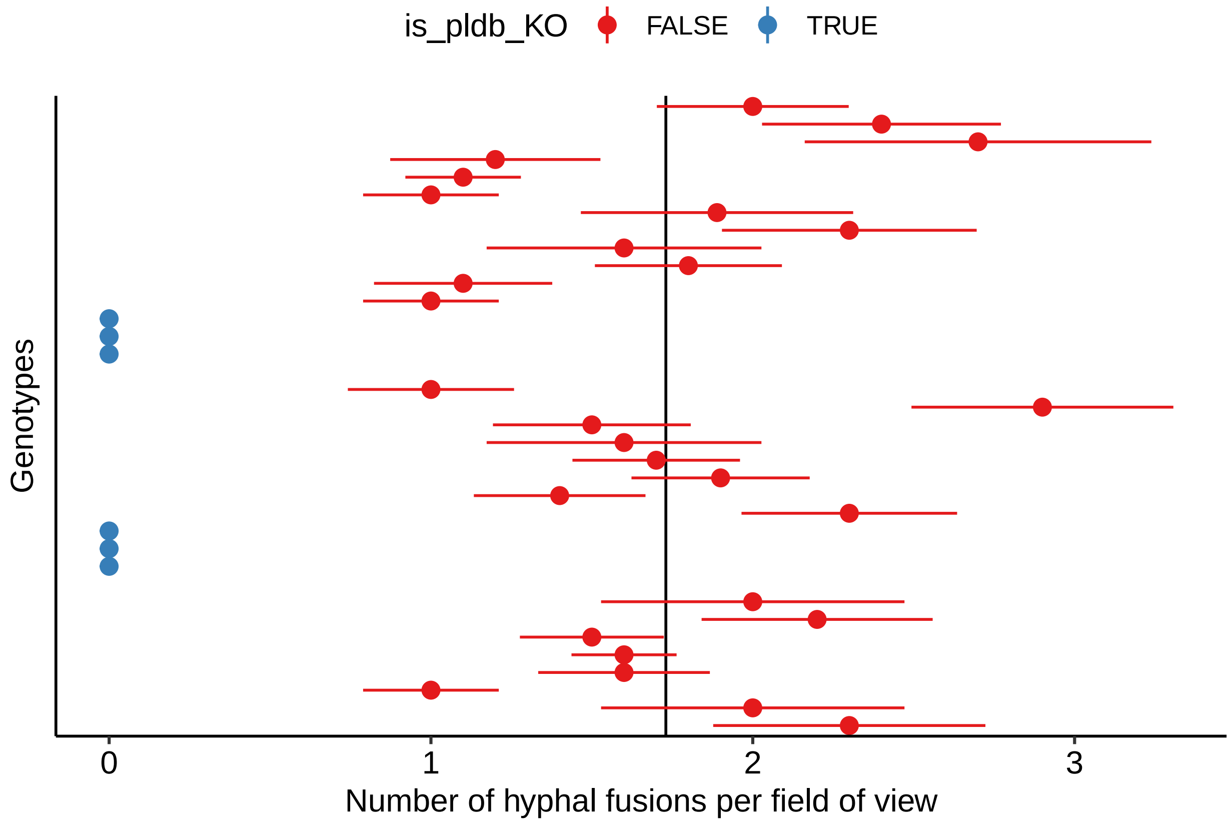


Excluding the *pldB* deletion strains, we observe the number of fusions counted per field of view is approximately Poisson distributed with a mean ($\lambda$) of ~ 1.734. We can illustrate this relationship by plotting the observed (bar plot) and expected (line) distributions of counts per observation.

```
all_fusions   <- fusions[!fusions$is_pldb_KO, "n_fusions"]
p <- barplot(table(all_fusions)/length(all_fusions),
             col="white",
             xlab="n.fusions",
             ylab="Proportion of observations")
lines(p[,1], dpois(0:7, lambda=1.734), lwd=3, col="steelblue", type="b")
```

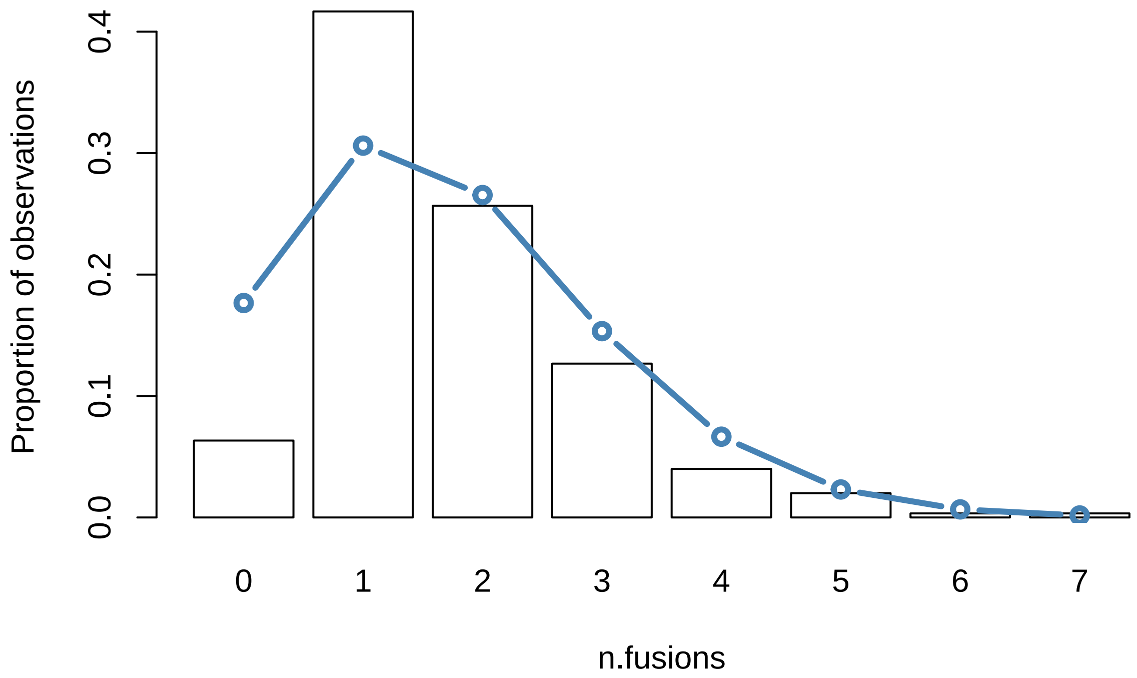


Using this relationship, we can construct a test of the null hypothesis that the *pldB deletions* lines produce hyphal fusions at the same rate as all other stains. Assuming the fusion counts per observation are Poisson-distributed, the probability of observing no fusions in 10 consecutive attempts is $Pois(x = 0, \lambda = 1.734)^{10}$, where $Pois$ is the Poisson density function. This gives a p-value:

```
dpois(0, lambda=1.734)^10
```

```
## [1] 2.946685e-08
```

Note, this test is likely to be conservative as the observed distribution appears to have fewer zeros than expected under the Poisson distribution.

##### Plant phenotypes.

To investigate whether the mutant strains influence the growth of the host plant we analysed two key plant phenotypes: the number and length of tillers (shoots) produced by grass infected by each strain.

**Example analyses**

The tiller data is reasonably straightforward to analyse. We fit a linear model with the plant phenotype as the explanatory value and strain as the dependant variable. We analyse the tiller length data from the *pldB* deletion strains as to demonstrate one analysis. First, we plot the data by strain.

```
tiller_lens <- read.csv("clean_data/pldB_tiller_lens.csv")

ggplot(tiller_lens, aes(genotype, value)) +
    geom_boxplot() +
    ylab("tiller length") +
    coord_flip()
```

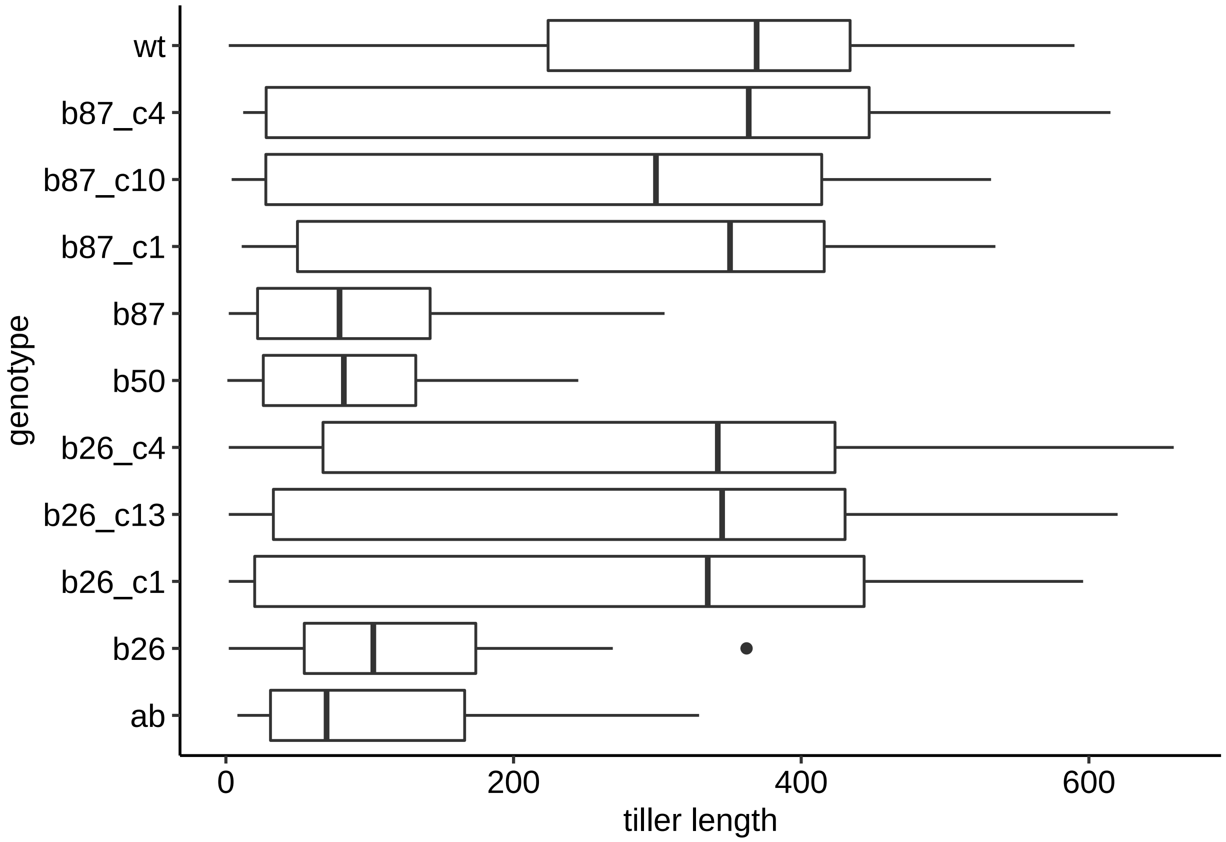


We can then fit the linear model and extract the result with `broom::tidy`.

```
library(broom)
tiller_lens$genotype <- relevel(tiller_lens$genotype, ref="wt")
fit <- lm(value ~ genotype,data=tiller_lens)
tidy_fit <- tidy(fit, conf.int= TRUE, conf.level=0.95)
#drop first row which is intercept
ggplot(tidy_fit[-1,], aes(term, estimate, ymax=conf.high, ymin=conf.low))  +
    geom_pointrange() +
```

```
    geom_hline(yintercept=0, lty=3, colour="red") +
    ylab("Difference in tiller length from WT") +
    xlab("Strain") +
    coord_flip()
```

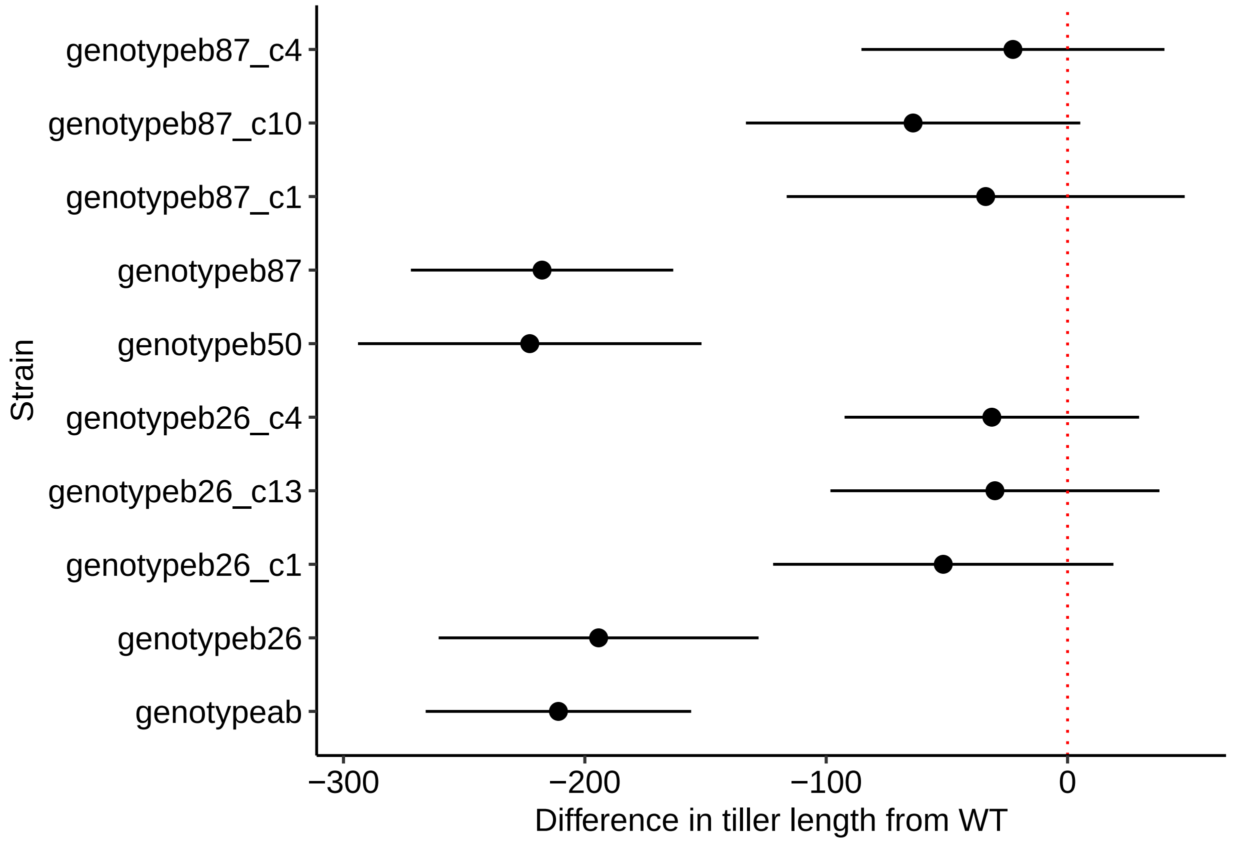


Here the point-ranges represent the estimate (+/- 95% confidence interval) of the difference in tiller lengths between each stain and the WT.

Finally, we can adjust the p-values in our results table to reflect the fact we have preformed multiple comparisons with the WT strain. Note, p-values represented as zero in these tables represent values less than $1 \times 10^{-8}$.

```
tidy_fit$p_adjusted <- p.adjust(tidy_fit$p.value)
knitr::kable(tidy_fit, digits = c(1,2,2,2,8,2,2,8) )
```

| term | estimate | std.error | statistic | p.value | conf.low | conf.high | p_adjusted |
|---|---|---|---|---|---|---|---|
| (Intercept) | 309.03 | 24.01 | 12.87 | 0.00000000 | 261.85 | 356.20 | 0.00000000 |
| genotypeab | -210.95 | 28.00 | -7.54 | 0.00000000 | -265.95 | -155.95 | 0.00000000 |
| genotypeb26 | -194.29 | 33.74 | -5.76 | 0.00000001 | -260.56 | -128.02 | 0.00000010 |
| genotypeb26_c1 | -51.49 | 35.89 | -1.43 | 0.15190200 | -121.99 | 19.00 | 0.75950999 |
| genotypeb26_c13 | -30.09 | 34.70 | -0.87 | 0.38633053 | -98.25 | 38.08 | 1.00000000 |
| genotypeb26_c4 | -31.37 | 31.06 | -1.01 | 0.31288432 | -92.38 | 29.64 | 1.00000000 |

| term | estimate | std.error | statistic | p.value | conf.low | conf.high | p_adjusted |
|---|---|---|---|---|---|---|---|
| genotypeb50 | -222.82 | 36.23 | -6.15 | 0.00000000 | -293.99 | -151.65 | 0.00000001 |
| genotypeb87 | -217.73 | 27.67 | -7.87 | 0.00000000 | -272.08 | -163.38 | 0.00000000 |
| genotypeb87_c1 | -33.92 | 41.98 | -0.81 | 0.41945600 | -116.38 | 48.54 | 1.00000000 |
| genotypeb87_c10 | -64.00 | 35.26 | -1.81 | 0.07010754 | -133.27 | 5.27 | 0.42064521 |
| genotypeb87_c4 | -22.63 | 31.95 | -0.71 | 0.47911914 | -85.41 | 40.14 | 1.00000000 |

**All other datasets**

Because separate experiments (including different WT plants) were run for deletion and over-expression strains, we analyse datasets separately (in each case comparing each mutant genotype to a WT strain that grown under the same conditions at the same time). We analyse each data set and provide a full results table in the Appendix of this report. These results include seperate analyses for the tiller-length data and the number of tiller per plant.

#### Hyphal counts per intercellular space

During growth, the spaces between plant cells are typically occupied by a single *Epichloë* hypha. Though all *Epichloë* strains occasionally form more than one hypha per intercellular space, we noticed this phenomenon appeared to be more frequent in *pldB* deletion strains. To illustrate this effect, we plot the distribution of hyphae-per-space across all genotypes.

```
hyphae <- read.csv("clean_data/hyphae_counts.csv")
ggplot(hyphae, aes(n.hyphae, fill=genotype)) +
    geom_bar(colour='black') +
    facet_wrap(~strain, nrow=3) +
    scale_fill_brewer(palette="Set1")
```

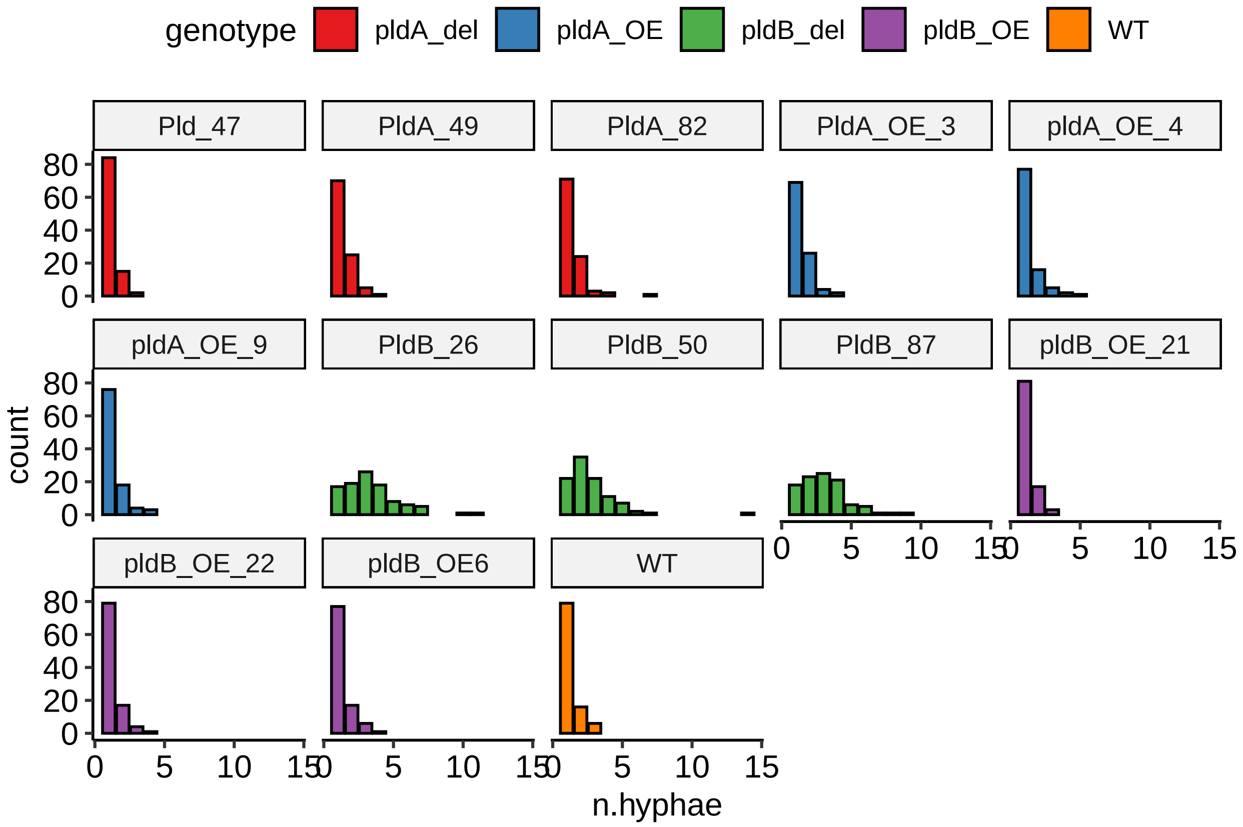


This distribution of this this data is unusual, in that (in most strains) the majority of intercellular spaces contain exactly one hypha. Most parametric tests will not accommodate this "one-biased" distribution. Instead we used a robust approach, namely the Dunn's test for multiple comparisons, to test for significant differences in the number of hyphae per intercellular space between all strains.

The table produced by the `dunn.test` function provided corrected p-values for all pair-wise comparisons in this study.

```
library(dunn.test)
dunn.test(hyphae$n.hyphae, hyphae$genotype, method="bonferroni" )
```

```
##   Kruskal-Wallis rank sum test
##
## data: x and group
## Kruskal-Wallis chi-squared = 421.6109, df = 4, p-value = 0
##
##
##                             Comparison of x by group
##                                   (Bonferroni)
## Col Mean-|
## Row Mean |    pldA_del    pldA_OE    pldB_del    pldB_OE
## ---------+-------------------------------------------
```

```
##   pldA_OE |  -0.408829
##           |     1.0000
##           |
## pldB_del |  -16.32028  -15.91145
##           |    0.0000*    0.0000*
##           |
##   pldB_OE |   0.815986   1.224816   17.13627
##           |     1.0000     1.0000    0.0000*
##           |
##        WT |   0.542151   0.831238   12.08233  -0.034837
##           |     1.0000     1.0000    0.0000*     1.0000
##
## alpha = 0.05
## Reject Ho if p <= alpha/2
```

#### Superoxide Staining

When treated with nitroblue tetrazolium, the hyphal tips of WT *Epichloë* cells typically stain dark blue. Some cells, however, generate abnormal staining patterns with the blue either patchy or entirely absent from the hyphal tip. Abnormal staining patterns appear to be more common in *pldB* deletion strains than in other strains.

```
staining <- read.csv("clean_data/staining.csv")
staining$genotype <- factor(
            staining$genotype, levels= rev(levels(staining$genotype))
)
staining$staining_type <- factor(
            staining$staining_type, levels=c("normal", "abnormal")
)

ggplot(staining, aes(replicate, n, fill=staining_type)) +
    geom_col(colour='black') +
    facet_grid(~ genotype) +
    scale_fill_brewer()
```

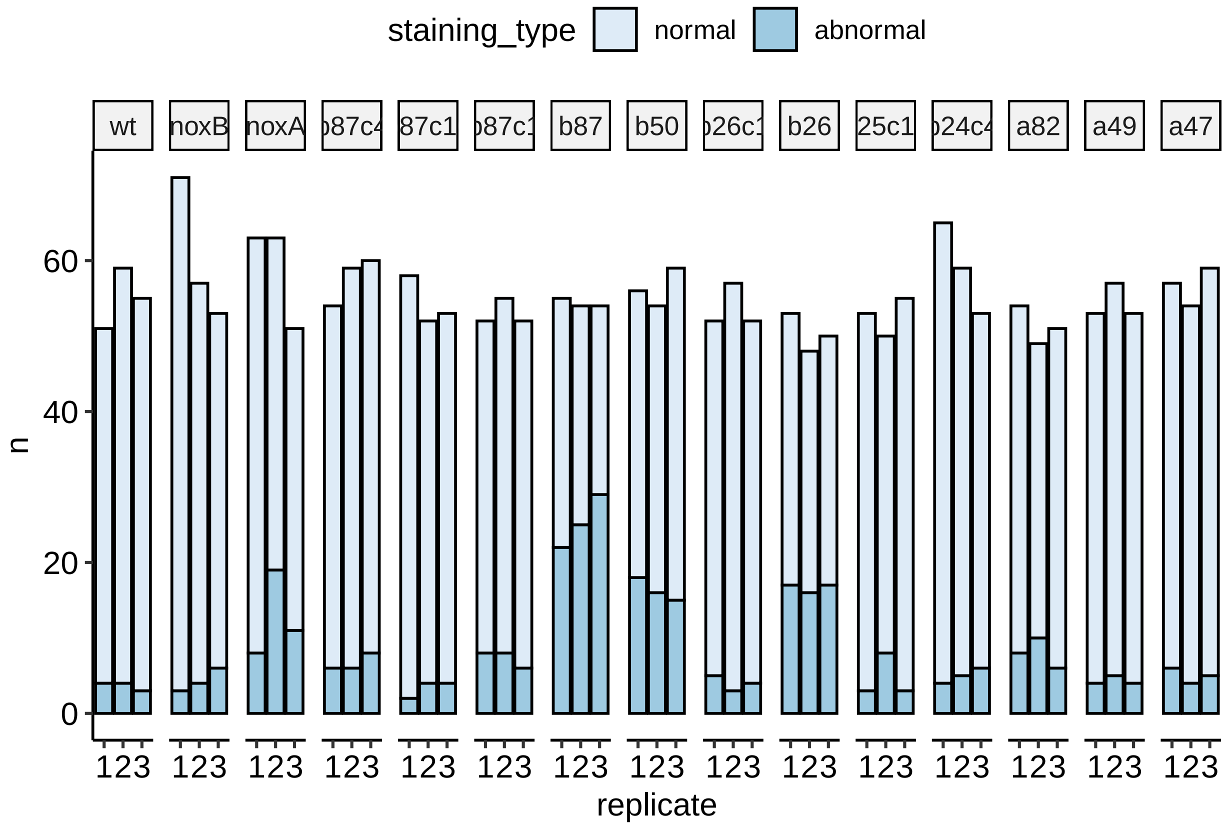


We use a generalized linear model with binomial link function (equivalent to a logistic regression) to test the null hypotheses that each strain gives rise to abnormally-staining cells at a rate equal to that of WT strains.

Here we fit the model and display odds-ratios for each each strain compared to WT.

```
by_geno <-  aggregate(n ~ replicate + genotype , data=staining, FUN=c)
fit <- glm(n ~ genotype, family=binomial, data=by_geno)
tidy_fit <- broom::tidy(fit, exponentiate=TRUE, conf.int=TRUE)

ggplot(tidy_fit[-1,], aes(term, estimate, ymax=conf.high, ymin=conf.low))  +
     geom_pointrange() +
     geom_hline(yintercept=1, lty=3, colour="red") +
     ylab("OR compared to WT") +
     xlab("Strain") +
     coord_flip()
```

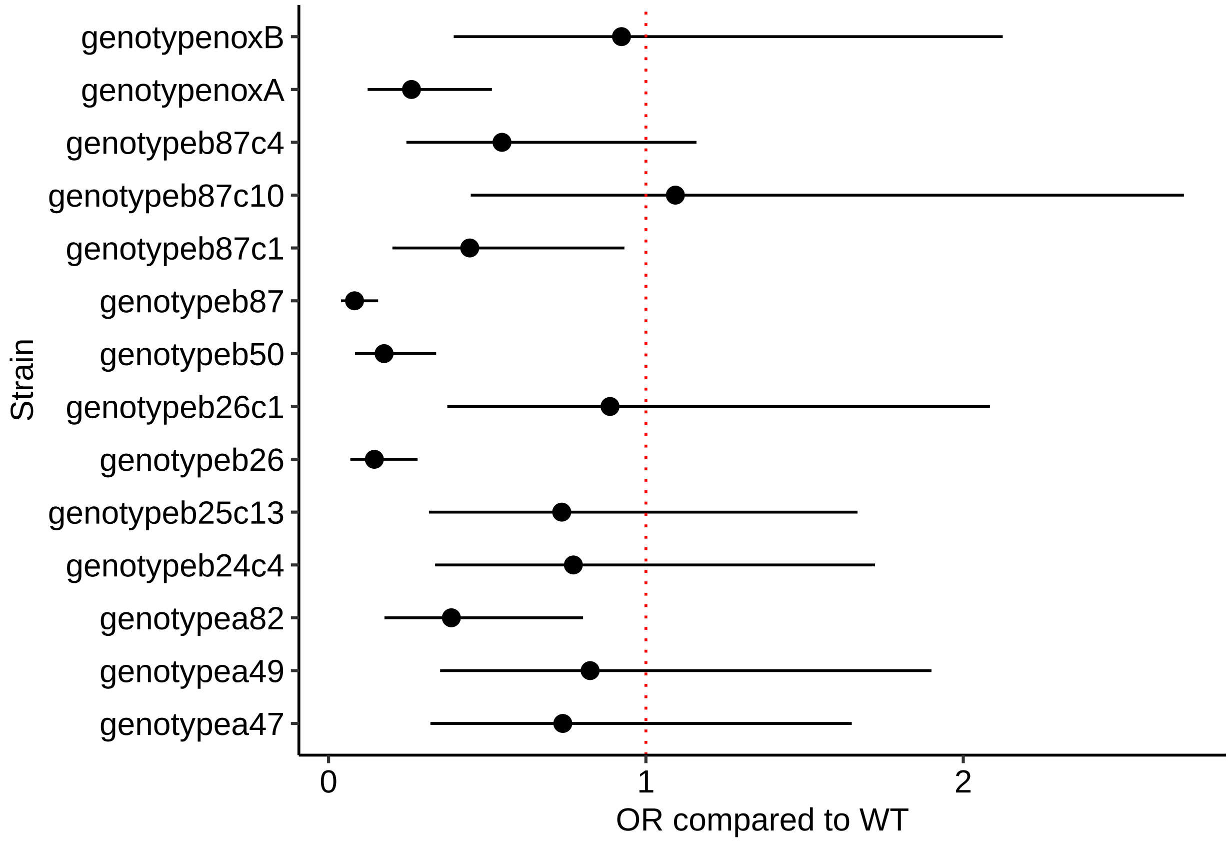


For an odds ratio, a value of 1 is equivalent to no difference between groups, so strains with error-bars not crossing the red line show a significant difference from WT. As above, we can correct the p-values arising from this test to account for the fact we have tested multiple independent strains in this work, then print the complete results table.

```
tidy_fit$adjusted_pvalue <- p.adjust(tidy_fit$p.value)
knitr::kable(tidy_fit,  digits = c(1,2,2,2,8,2,2,8))
```

| term | estimate | std.error | statistic | p.value | conf.low | conf.high | adjusted_pvalue |
|---|---|---|---|---|---|---|---|
| (Intercept) | 14.00 | 0.31 | 8.46 | 0.00000000 | 7.97 | 27.42 | 0.00000000 |
| genotypenoxB | 0.92 | 0.42 | -0.19 | 0.85047140 | 0.39 | 2.12 | 1.00000000 |
| genotypenoxA | 0.26 | 0.36 | -3.71 | 0.00020766 | 0.12 | 0.51 | 0.00228426 |
| genotypeb87c4 | 0.55 | 0.39 | -1.54 | 0.12347797 | 0.25 | 1.16 | 0.98782377 |
| genotypeb87c10 | 1.09 | 0.45 | 0.20 | 0.84412008 | 0.45 | 2.70 | 1.00000000 |
| genotypeb87c1 | 0.44 | 0.39 | -2.09 | 0.03656141 | 0.20 | 0.93 | 0.32905267 |
| genotypeb87 | 0.08 | 0.35 | -7.17 | 0.00000000 | 0.04 | 0.16 | 0.00000000 |
| genotypeb50 | 0.17 | 0.36 | -4.91 | 0.00000092 | 0.08 | 0.34 | 0.00001101 |
| genotypeb26c1 | 0.89 | 0.43 | -0.28 | 0.78162049 | 0.37 | 2.08 | 1.00000000 |
| genotypeb26 | 0.14 | 0.36 | -5.43 | 0.00000006 | 0.07 | 0.28 | 0.00000075 |
| genotypeb25c13 | 0.73 | 0.42 | -0.74 | 0.46212345 | 0.32 | 1.67 | 1.00000000 |
| genotypeb24c4 | 0.77 | 0.41 | -0.63 | 0.52937563 | 0.34 | 1.72 | 1.00000000 |
| genotypea82 | 0.39 | 0.38 | -2.48 | 0.01318623 | 0.18 | 0.80 | 0.13186235 |

| term | estimate | std.error | statistic | p.value | conf.low | conf.high | adjusted_pvalue |
|---|---|---|---|---|---|---|---|
| genotypea49 | 0.82 | 0.43 | -0.45 | 0.64944772 | 0.35 | 1.90 | 1.00000000 |
| genotypea47 | 0.74 | 0.41 | -0.74 | 0.46208846 | 0.32 | 1.65 | 1.00000000 |

### Appendix

This appendix includes the results of linear models fitted to each plant phenotype dataset. To analyse these datasets we introduce a function, `fit and summarise`, that automates the tests described in the "Plant Phenotypes" section above.

```
fit_and_summarise <- function(fname, stringsAsFactors=FALSE, kable=TRUE){
    #read data and set WT to reference cat.
    mutant_data <- read.csv(fname)
    mutant_data$genotype <-  as.factor(toupper(mutant_data$genotype))
    mutant_data$genotype <- relevel(mutant_data$genotype, ref="WT")
    # fit model and make tidy data
    fit <- lm(value ~ genotype,data=mutant_data)
    tidy_fit <- tidy(fit, conf.int= TRUE, conf.level=0.95)
    # adjust p-values and return results table
    tidy_fit$adjusted_pvalue <- p.adjust(tidy_fit$p.value)
    if(kable){
        return(knitr::kable(tidy_fit,
                            digits = c(1,2,2,2,8,2,2,8)))
    }
    tidy_fit
}
```

We use this function to test each dataset.

#### pldB deletion strains

##### Tiller Length

```
fit_and_summarise("clean_data/pldB_tiller_lens.csv")
```

| term | estimate | std.error | statistic | p.value | conf.low | conf.high | adjusted_pvalue |
|---|---|---|---|---|---|---|---|
| (Intercept) | 309.03 | 24.01 | 12.87 | 0.00000000 | 261.85 | 356.20 | 0.00000000 |
| genotypeAB | -210.95 | 28.00 | -7.54 | 0.00000000 | -265.95 | -155.95 | 0.00000000 |
| genotypeB26 | -194.29 | 33.74 | -5.76 | 0.00000001 | -260.56 | -128.02 | 0.00000010 |
| genotypeB26_C1 | -51.49 | 35.89 | -1.43 | 0.15190200 | -121.99 | 19.00 | 0.75950999 |
| genotypeB26_C13 | -30.09 | 34.70 | -0.87 | 0.38633053 | -98.25 | 38.08 | 1.00000000 |

| term | estimate | std.error | statistic | p.value | conf.low | conf.high | adjusted_pvalue |
|---|---|---|---|---|---|---|---|
| genotypeB26_C4 | -31.37 | 31.06 | -1.01 | 0.31288432 | -92.38 | 29.64 | 1.00000000 |
| genotypeB50 | -222.82 | 36.23 | -6.15 | 0.00000000 | -293.99 | -151.65 | 0.00000001 |
| genotypeB87 | -217.73 | 27.67 | -7.87 | 0.00000000 | -272.08 | -163.38 | 0.00000000 |
| genotypeB87_C1 | -33.92 | 41.98 | -0.81 | 0.41945600 | -116.38 | 48.54 | 1.00000000 |
| genotypeB87_C10 | -64.00 | 35.26 | -1.81 | 0.07010754 | -133.27 | 5.27 | 0.42064521 |
| genotypeB87_C4 | -22.63 | 31.95 | -0.71 | 0.47911914 | -85.41 | 40.14 | 1.00000000 |

###### Tiller Number

```
fit_and_summarise("clean_data/pldB_tiller_num.csv")
```

| term | estimate | std.error | statistic | p.value | conf.low | conf.high | adjusted_pvalue |
|---|---|---|---|---|---|---|---|
| (Intercept) | 3.36 | 0.93 | 3.63 | 0.00047137 | 1.52 | 5.21 | 0.00424233 |
| genotypeAB | 11.35 | 1.49 | 7.63 | 0.00000000 | 8.40 | 14.30 | 0.00000000 |
| genotypeB26 | 1.89 | 1.43 | 1.32 | 0.19008792 | -0.95 | 4.72 | 1.00000000 |
| genotypeB26_C1 | -0.64 | 1.31 | -0.49 | 0.62861587 | -3.24 | 1.97 | 1.00000000 |
| genotypeB26_C13 | 0.04 | 1.34 | 0.03 | 0.97846779 | -2.63 | 2.71 | 1.00000000 |
| genotypeB26_C4 | 2.04 | 1.34 | 1.52 | 0.13308764 | -0.63 | 4.71 | 1.00000000 |
| genotypeB50 | 1.47 | 1.56 | 0.94 | 0.34883500 | -1.63 | 4.57 | 1.00000000 |
| genotypeB87 | 12.49 | 1.49 | 8.40 | 0.00000000 | 9.54 | 15.45 | 0.00000000 |
| genotypeB87_C1 | -1.36 | 1.38 | -0.99 | 0.32645322 | -4.11 | 1.38 | 1.00000000 |
| genotypeB87_C10 | -0.16 | 1.34 | -0.12 | 0.90333471 | -2.83 | 2.51 | 1.00000000 |
| genotypeB87_C4 | 0.33 | 1.26 | 0.26 | 0.79476308 | -2.17 | 2.83 | 1.00000000 |

##### pldA deletion strains

###### Tiller Length

```
fitted_tiller_num <- fit_and_summarise("clean_data/pldA_tiller_num.csv",
                                       kable= FALSE)

knitr::kable(fitted_tiller_num[1:4,],  digits = c(1,2,2,2,8,2,2,8))
```

| term | estimate | std.error | statistic | p.value | conf.low | conf.high | adjusted_pvalue |
|---|---|---|---|---|---|---|---|
| (Intercept) | 7.00 | 1.40 | 4.98 | 0.00001585 | 4.15 | 9.85 | 0.00011097 |
| genotypePLDA_KO47 | -0.50 | 1.90 | -0.26 | 0.79417162 | -4.36 | 3.36 | 1.00000000 |
| genotypePLDA_KO49 | -0.67 | 1.90 | -0.35 | 0.72803993 | -4.52 | 3.19 | 1.00000000 |
| genotypePLDA_KO82 | 0.25 | 2.11 | 0.12 | 0.90622843 | -4.02 | 4.52 | 1.00000000 |

##### Tiller Number

```
fitted_tiller_len <- fit_and_summarise("clean_data/pldA_tiller_lens.csv",
                                       kable= FALSE
)
knitr::kable(fitted_tiller_len[1:4,],  digits = c(1,2,2,2,8,2,2,8))
```

| term | estimate | std.error | statistic | p.value | conf.low | conf.high | adjusted_pvalue |
|---|---|---|---|---|---|---|---|
| (Intercept) | 206.89 | 25.58 | 8.09 | 0.0000000 | 156.54 | 257.23 | 0 |
| genotypePLDA_KO47 | 13.22 | 35.23 | 0.38 | 0.7078589 | -56.14 | 82.57 | 1 |
| genotypePLDA_KO49 | 19.48 | 35.45 | 0.55 | 0.5830715 | -50.30 | 89.27 | 1 |
| genotypePLDA_KO82 | 5.87 | 38.00 | 0.15 | 0.8772840 | -68.92 | 80.67 | 1 |

#### pldA over-expression strains

##### Tiller Length

```
knitr::kable(fitted_tiller_len[c(1,5:7),],  digits = c(1,2,2,2,8,2,2,8))
```

| term | estimate | std.error | statistic | p.value | conf.low | conf.high | adjusted_pvalue |
|---|---|---|---|---|---|---|---|
| (Intercept) | 206.89 | 25.58 | 8.09 | 0.0000000 | 156.54 | 257.23 | 0 |
| genotypePLDA_OE3 | 26.30 | 34.63 | 0.76 | 0.4481826 | -41.87 | 94.48 | 1 |
| genotypePLDA_OE4 | 17.11 | 34.27 | 0.50 | 0.6179351 | -50.35 | 84.58 | 1 |
| genotypePLDA_OE9 | -7.86 | 31.64 | -0.25 | 0.8041184 | -70.14 | 54.43 | 1 |

##### Tiller Number

```
fit_and_summarise("clean_data/OE_tiller_num.csv")
```

| term | estimate | std.error | statistic | p.value | conf.low | conf.high | adjusted_pvalue |
|---|---|---|---|---|---|---|---|
| (Intercept) | 2.17 | 0.25 | 8.79 | 0.00000026 | 1.64 | 2.69 | 0.00000106 |
| genotypeBOE21 | 0.17 | 0.35 | 0.48 | 0.63941477 | -0.58 | 0.91 | 1.00000000 |
| genotypeBOE22 | -0.67 | 0.49 | -1.35 | 0.19625252 | -1.72 | 0.38 | 0.58875757 |
| genotypeBOE6 | 0.03 | 0.37 | 0.09 | 0.92855151 | -0.75 | 0.81 | 1.00000000 |

#### pldB over-expression strains

##### Tiller Length

```r
fit_and_summarise("clean_data/OE_tiller_lens.csv")
```

| term | estimate | std.error | statistic | p.value | conf.low | conf.high | adjusted_pvalue |
|---|---|---|---|---|---|---|---|
| (Intercept) | 168.92 | 28.98 | 5.83 | 0.00000107 | 110.20 | 227.65 | 0.00000429 |
| genotypeBOE21 | 54.79 | 40.25 | 1.36 | 0.18163798 | -26.76 | 136.34 | 0.54491394 |
| genotypeBOE22 | 35.08 | 66.93 | 0.52 | 0.60334866 | -100.54 | 170.69 | 1.00000000 |
| genotypeBOE6 | 15.44 | 42.81 | 0.36 | 0.72038553 | -71.30 | 102.18 | 1.00000000 |

##### Tiller number

```r
knitr::kable(fitted_tiller_num[c(1,5:7),],  digits = c(1,2,2,2,8,2,2,8))
```

| term | estimate | std.error | statistic | p.value | conf.low | conf.high | adjusted_pvalue |
|---|---|---|---|---|---|---|---|
| (Intercept) | 7.00 | 1.40 | 4.98 | 0.00001585 | 4.15 | 9.85 | 0.00011097 |
| genotypePLDA_OE3 | 0.17 | 1.90 | 0.09 | 0.93066930 | -3.69 | 4.02 | 1.00000000 |
| genotypePLDA_OE4 | -1.50 | 1.79 | -0.84 | 0.40781225 | -5.13 | 2.13 | 1.00000000 |
| genotypePLDA_OE9 | 1.12 | 1.79 | 0.63 | 0.53386484 | -2.51 | 4.76 | 1.00000000 |
